# Fundamental limits on the rate of bacterial growth

**DOI:** 10.1101/2020.10.18.344382

**Authors:** Nathan M. Belliveau, Grifin Chure, Christina L. Hueschen, Hernan G. Garcia, Jane Kondev, Daniel S. Fisher, Julie A. Theriot, Rob Phillips

## Abstract

Recent years have seen an experimental deluge interrogating the relationship between bacterial growth rate, cell size, and protein content, quantifying the abundance of proteins across growth conditions with unprecedented resolution. However, we still lack a rigorous understanding of what sets the scale of these quantities and when protein abundances should (or should not) depend on growth rate. Here, we seek to quantitatively understand this relationship across a collection of *Escherichia coli* proteomic data covering ≈ 4000 proteins and 36 growth rates. We estimate the basic requirements for steady-state growth by considering key processes in nutrient transport, cell envelope biogenesis, energy generation, and the central dogma. From these estimates, ribosome biogenesis emerges as a primary determinant of growth rate. We expand on this assessment by exploring a model of proteomic regulation as a function of the nutrient supply, revealing a mechanism that ties cell size and growth rate to ribosomal content.

## Introduction

The observed range of bacterial growth rates is enormously diverse. In natural environments, some microbial organisms may double only once per year (***Mikucki et al., 2009***) while in comfortable laboratory conditions, growth can be rapid with several divisions per hour (***Schaechter et al., 1958***). This six order-of-magnitude difference in time scales of growth encompasses different microbial species and lifestyles, yet even for a single species such as *Escherichia coli*, the growth rate can be modulated over a large scale by tuning the type and amount of nutrients in the growth medium (***Liu et al., 2005***). This remarkable plasticity in growth rate illustrates the intimate relationship between environmental conditions and the rates at which cells convert nutrients into new cellular material – a relationship that has remained a major topic of inquiry in bacterial physiology for over a century (***Jun et al., 2018***).

A key discovery in bacterial physiology of the past 70 years was the identification of bacterial “growth laws” (***Schaechter et al., 1958***); empirical relationships that relate the bacterial growth rate to the protein and RNA composition of the intracellular milieu in a number of different species. Over the past decade, a flurry of work (***Molenaar et al., 2009***; ***Scott et al., 2010***; ***Klumpp and Hwa, 2014***; ***Basan et al., 2015***; ***Dai et al., 2016***; ***Erickson et al., 2017***) has examined these growth laws at a quantitative level, developing a series of phenomenological models from which the growth laws naturally emerge. In parallel, a “molecular revolution” in biology has yielded an increasingly refined molecular census of the cell, particularly for bacteria such as the microbial workhorse *E. coli* (***Schmidt et al., 2016***; ***Davidi et al., 2016***). In light of the now expansive trove of quantitative biological data, we can revisit several of the evergreen questions about bacterial growth and physiology that were originally raised by microbiologists in the middle of the 20th century. Specifically, what biological processes are the primary determinants for how quickly bacterial cells can grow and reproduce? Why do cells modulate the absolute numbers and relative ratios of their molecular constituents in response to changes in growth rate or nutrient availability?

In this work, we begin by considering these two questions from two distinct angles. First, as a result of an array of high-quality proteome-wide measurements of *E. coli* under diverse growth conditions, we have generated a census that allows us to explore how the number of key molecular players change as a function of growth rate. Here, we have assembled a singular data set of protein copy numbers using measurements collected over the past decade via mass spectrometry (***Schmidt et al., 2016***; ***Peebo et al., 2015***; ***Valgepea et al., 2013***) or ribosomal profiling (***Li et al., 2014***) of the composition of the *E. coli* proteome across a gamut of growth rates. Due to notable changes in cell size and cellular composition as a function of growth rate (***Bremer and Dennis, 2008***; ***Taheri-Araghi et al., 2015***), as well as differences in normalization and standardization schemes used in each experimental work, substantial care was taken to ensure consistency on a per cellular basis (see the Appendix for a detailed analysis and additional discussion). To our knowledge, this compiled and curated dataset represents the most comprehensive view to date of the *E. coli* proteome, covering ≈ 4000 proteins and 36 unique growth rates, with the observed abundance of any given protein being directly comparable between data sets and across growth rates. This allows us to interrogate the *E. coli* specific physiology underlying the observed abundances while minimizing the effects of experimental noise as ≈ 75% of the proteins are observed in at least two separate datasets.

Second, by compiling molecular turnover rate measurements for many of the fundamental processes associated with bacterial growth, we make quantitative order-of-magnitude estimates of key cellular processes in nutrient transport, cell envelope biogenesis, energy generation, and the central dogma (schematized in ***Figure 1***) to determine whether our current understanding of the kinetics of these processes are sufficient to explain the magnitude of the observed protein copy numbers across conditions (see ***Box 1*** describing the philosophy behind this approach). The census, combined with these estimates, provide a window into the question of whether the rates of central processes such as energy generation or DNA synthesis vary systematically as a function of cell growth rate by altering protein copy number, and in particular, whether any of these processes may be limiting growth.

**Figure 1.**
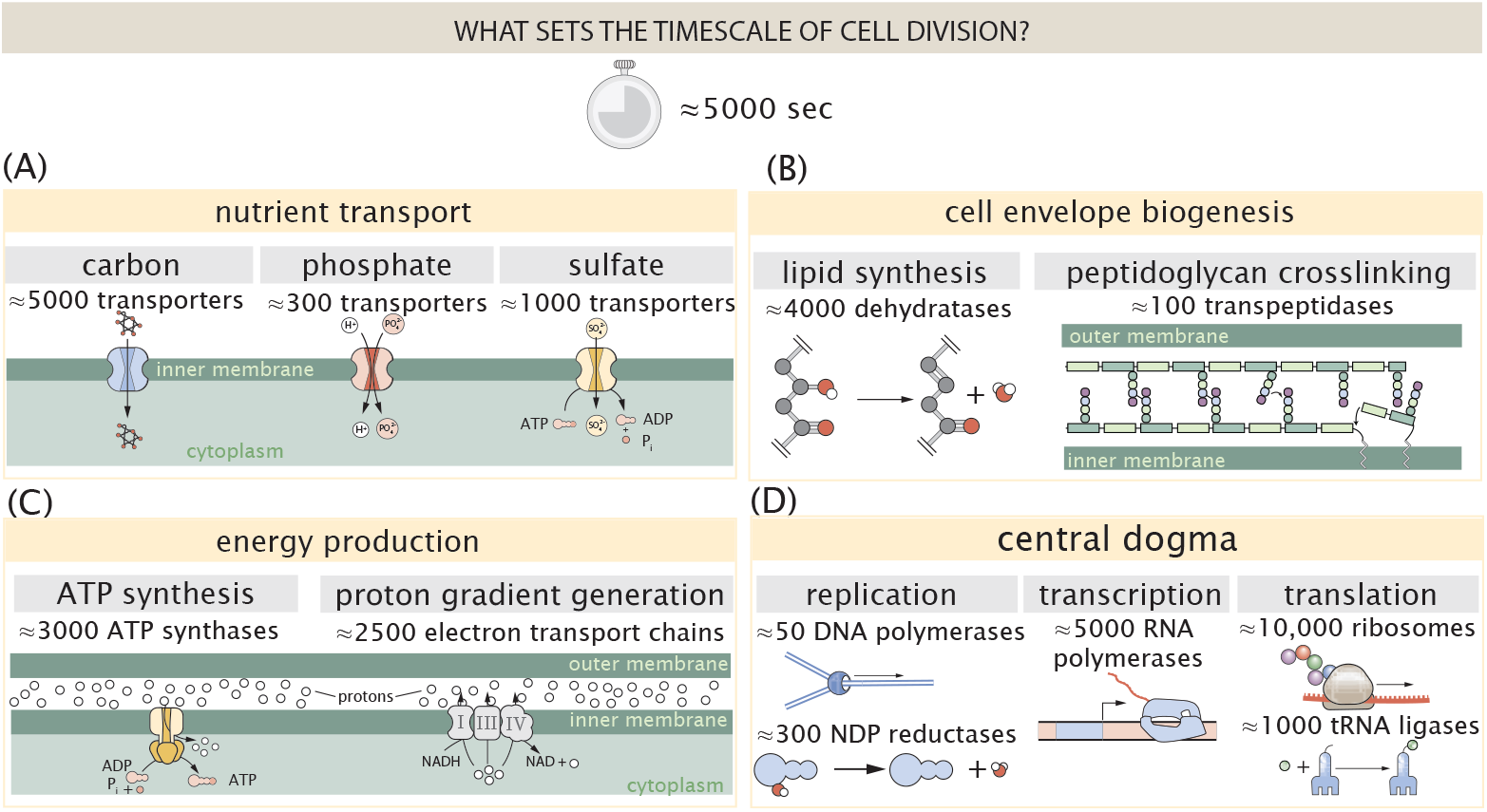
Transport and synthesis processes necessary for cell division. We consider an array of processes necessary for a cell to double its molecular components, broadly grouped into four classes. These categories are (A) nutrient transport across the cell membrane, (B) cell envelope biogenesis, (C) energy production (namely, ATP synthesis), and (D) processes associated with the central dogma. Numbers shown are the approximate number of complexes of each type observed at a growth rate of 0.5 hr^−1^, or a cell doubling time of ≈ 5000 s.

Throughout our estimates, we consider an archetypal growth rate of ≈ 0.5 hr^−1^ corresponding to a doubling time of ≈ 5000 seconds, as the data sets examined here heavily sample this growth regime. While we formulate point estimates for the protein abundances at this division time, we also consider how these values will vary at other growth rates due to changes in cell size, surface area, and chromosome copy number (***Taheri-Araghi et al., 2015***; ***Harris and Theriot, 2018***). For the majority of the processes considered, we find that the protein copy numbers appear tuned for the task of cell doubling across a continuum of growth rates. Thus, our understanding of the kinetics of various biological processes is sufficient to quantitatively explain the observed abundances of these proteins.

From these estimates, it emerges that translation, particularly the synthesis of ribosomal proteins is a plausible candidate that limits the rate of cellular growth in *E. coli*. We reach this conclusion by considering that ribosome synthesis is 1) a rate limiting step for the *fastest* bacterial division, and 2) the main determinant of bacterial growth rate across nutrient conditions associated with moderate to fast growth rates. In addition, a strict dependence between the maximal growth rate and ribosomal mass fraction coincides with the regime where the growth laws appear most valid (***Amir, 2017***; ***Scott et al., 2010***). This enables us to suggest that the long-observed correlation between growth rate and cell size (***Schaechter et al., 1958***; ***Si et al., 2017***) can be simply attributed to the increased absolute number of ribosomes per cell under conditions supporting extremely rapid growth. To better understand how the observed alterations in absolute protein abundances (and in particular, changes in ribosome copy number) influence growth rate across different nutrient conditions, we consider a minimal model of cellular growth. Our conclusions from these analyses provide important insight into how *E. coli* regulates growth across conditions of differing nutrient availability and identifies fundamental constraints in bacterial growth more broadly.

### Nutrient Transport

We begin by considering the critical transport processes diagrammed in ***Figure 1*(A)**. In order to build new cellular mass, the molecular and elemental building blocks must be scavenged from the environment in different forms. Carbon, for example, is acquired via the transport of carbohydrates and sugar alcohols with some carbon sources receiving preferential treatment in their consumption (***Monod, 1947***). Phosphorus, sulfur, and nitrogen, on the other hand, are harvested primarily in the forms of inorganic salts, namely phosphate, sulfate, and ammonium/ammonia (***Jun et al., 2018***; ***Assentoft et al., 2016***; ***Stasi et al., 2019***; ***Antonenko et al., 1997***; ***Rosenberg et al., 1977***; ***Willsky et al., 1973***). All of these compounds have different membrane permeabilities (***Phillips, 2018***) and most require some energetic investment either via ATP hydrolysis or through the proton electrochemical gradient to bring the material across the hydrophobic cell membrane.

##### Box 1. The Rules of Engagement for Order-Of-Magnitude Estimates

This work relies heavily on “back-of-the-envelope” estimates to understand the growth-rate dependent abundances of molecular complexes. This moniker arises from the limitation that any estimate should be able to fit on the back of a postage envelope. As such, we must draw a set of rules governing our precision and sources of key values.

**The rule of “one, few, and ten”**. The philosophy behind order-of-magnitude estimates is to provide an estimate of the appropriate scale, not a prediction with many significant digits (***Mahajan, 2010***). We therefore define three different scales of precision in making estimates. The scale of “one” is reserved for values that range between 1 and 2. For example, If a particular process has been experimentally measured to transport 1.87 protons for a process to occur, we approximate this process to require 2 protons per event. The scale of “few” is reserved for values ranging between 3 and 7. For example, we will often use Avogadro’s number to compute the number of molecules in a cell given a concentration and a volume. Rather than using Avogadro’s number as 6.02214 × 10^23^, we will approximate it as 5 × 10^23^. Finally, the scale of “ten” is reserved for values which we know within an order of magnitude. If a particular protein complex is present at 883 copies per cell, we say that it is present in approximately 103 copies per cell. These different scales will be used to arrive at simple estimates that report the expected scale of the observed data. Therefore, the estimates presented here should not be viewed as hard-and-fast predictions of precise copy numbers, but as approximate lower (or upper) bounds for the number of complexes that may be needed to satisfy some cellular requirement.

Furthermore, we use equality symbols (=) sparingly and frequently defer to an approximation (≈) symbol when reporting an estimate, indicating that we are confident in the estimate to within a factor of a few.

**The BioNumbers Database as a source for values**. In making our estimates, we often require approximate values for key cellular properties, such as the elemental composition of the cell, the average dry mass, or approximate rates of synthesis. We rely heavily on the BioNumbers Database (bionumbers.hms.harvard.edu, **Milo et al. (2010)**) as a repository for such information. Every value we draw from this database has an associated BioNumbers ID number, abbreviated as BNID, and we provide this reference in grey-boxes in each figure.

**Uncertainty in the data sets and the accuracy of an estimate**. The data sets presented in this work are the products of careful experimentation with the aim to report, to the best of their ability, the absolute copy numbers of proteins in the cell. These data, collected over the span of a few years, come from different labs and use different internal standards, controls, and even techniques (discussed further in the Appendix Section “Experimental Details Behind Proteomic Data”). As a result, there is notable disagreement in the measured copy numbers for some complexes across data sets. In assessing whether our estimates could explain the observed scales and growth-rate dependencies, we also considered the degree of variation between the different data sets. For example, say a particular estimate undercuts the observed data by an order of magnitude. If all data sets agree within a factor of a few of each other, we revisit our estimate and consider what me may have missed. However, if the data sets themselves disagree by an order of magnitude, we determine that our estimate is appropriate given the variation in the data.

**Point versus continuum estimates**. For each estimate performed in this work, we begin with a simple order-of-magnitude estimate for the abundance of the complex in question at an archetypal growth rate of around 0.5 hr^−1^, followed by a more refined estimate across a continuum of growth rates from around 0.05 to 2.0 hr^−1^. The former estimate, always outlined in the associated figure and indicated as a translucent brown point in the corresponding plots, will rely on making coarse-grained approximations of cell mass, cell volume, and/or typical surface areas. The continuum estimates, displayed as a grey curve on the various plots, relax these assertions and incorporate empirical findings from the literature of how cell masses, volumes, and surface areas scale with the cellular growth rate. Thus, it is possible for the point estimate at a growth rate of 0.5 hr^−1^ may not perfectly agree with the continuum estimate at 0.5 hr^−1^. We emphasize that both the point and continuum estimates are not hard predictions for the complex abundance, but rather reflect an order-of-magnitude estimates. While the proteomic measurements will often not fall directly on the curve corresponding to the continuum estimate, we do not view this as a failure of the approach. Rather, we gauge the accuracy of our estimates by examining whether the 1) the point *and* continuum estimates are within the same order of magnitude as the observations and 2) whether the continuum estimate qualitatively captures the growth-rate dependence observed in the data. We direct the reader to the Appendix Section “Extending Estimates to a Continuum of Growth Rates” for a more in-depth discussion of point versus continuum estimates.

The elemental composition of *E. coli* has received much quantitative attention over the past half century (***Neidhardt et al., 1991***; ***Taymaz-Nikerel et al., 2010***; ***Heldal et al., 1985***; ***Bauer and Ziv, 1976***), providing us with a starting point for estimating how many atoms of each element must be scavenged from the environment. A synthesis of these studies presents an approximate dry mass composition of ≈ 50% carbon (BNID: 100649; see ***Box 1*** for explanation of BNID references), ≈ 15% nitrogen (BNID: 106666), ≈ 3% phosphorus (BNID: 100653), and 1% sulfur (BNID: 100655) with the remainder being attributable to oxygen, hydrogen, and various transition metals. Here we use this stoichiometric breakdown to estimate the abundance and growth rate dependence of a variety of transporters responsible for carbon uptake, and provide more extensive investigation of the other critical elements - phosphorus, sulfur, and nitrogen - in the Appendix Section “Additional Estimates of Fundamental Biological Processes”.

Using ≈ 0.3 pg as the typical *E. coli* dry mass at a growth rate of ≈ 0.5 hr^−1^ (BNID: 103904), coupled with the approximation that ≈ 50% of this mass is carbon, we estimate that ≈ 1 × 10^10^ carbon atoms must be brought into the cell in order to double all of the carbon-containing molecules [***Figure 2***(A, top)]. Typical laboratory growth conditions provide carbon as a single class of sugar (such as glucose, galactose, or xylose) often transported cross the cell membrane by a transporter complex specific to that particular sugar. One such mechanism of transport is via the PTS system, which is a highly modular system capable of transporting a diverse range of sugars with high specificity (***Escalante et al., 2012***). The glucose-specific component of this system transports ≈ 200 glucose molecules (≈ 1200 carbon atoms) per second per transporter (BNID: 114686). Making the assumption that this is a typical sugar transport rate for the PTS system, coupled with the need to transport ≈ 1 × 10^10^ carbon atoms, we then expect on the order of ≈ 2000 transporters must be expressed per cell in order to bring in enough carbon atoms [***Figure 2***(A, top)].

**Figure 2.**
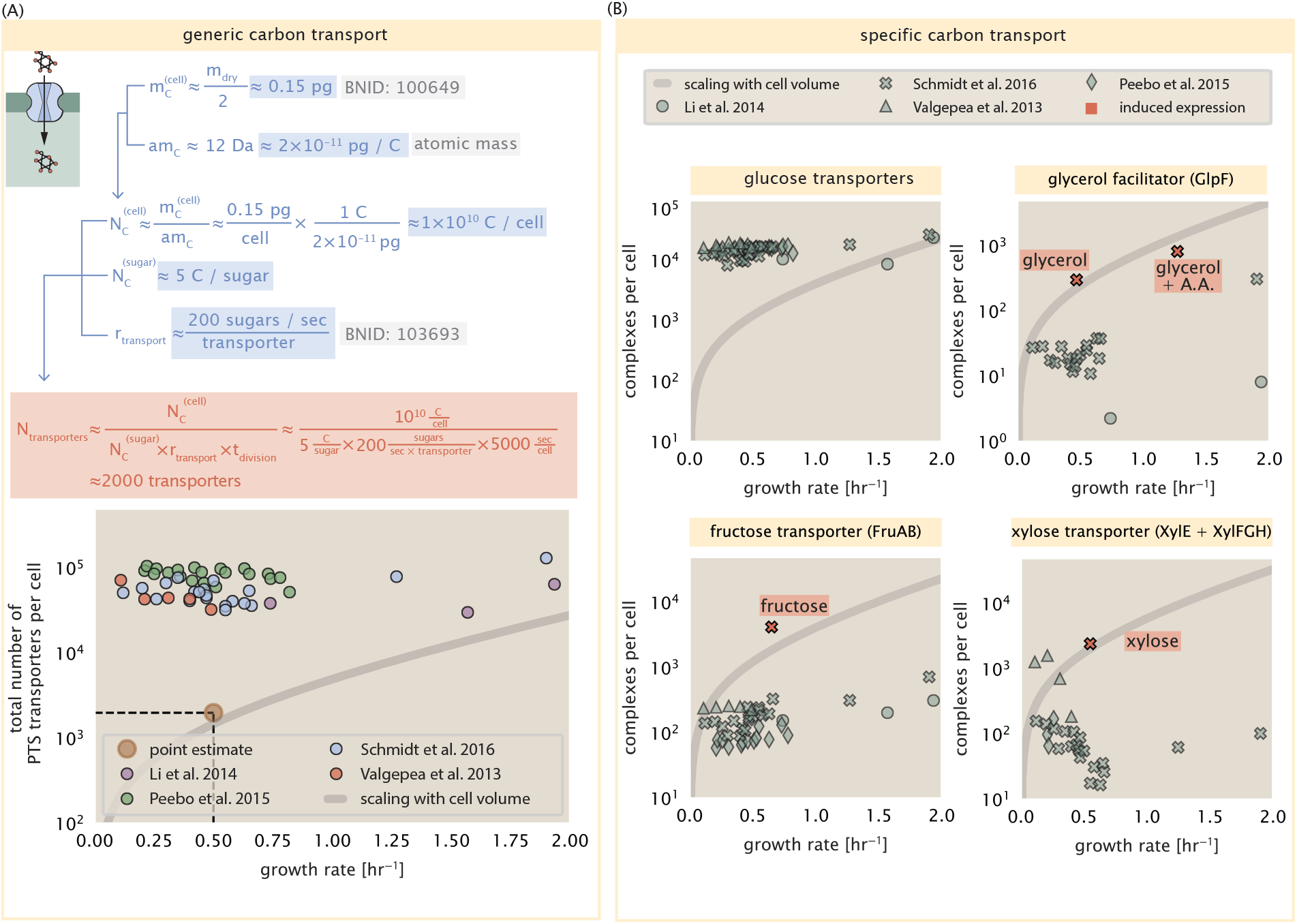
The abundance of carbon transport systems across growth rates. (A) A simple estimate for the minimum number of generic carbohydrate transport systems (top) assumes. 1 ≈ 10^10^ C are needed to complete division, each transported sugar contains.≈ 5 C, and each transporter conducts sugar molecules at a rate of. ≈ 200 per second. Bottom plot shows the estimated number of transporters needed at a growth rate of ≈ 0.5 per hr (light-brown point and dashed lines). Colored points correspond to the mean number of complexes involved in carbohydrate import (complexes annotated with the Gene Ontology terms GO:0009401 and GO:0098704) for different growth conditions across different published datasets. (B) The abundance of various specific carbon transport systems plotted as a function of the population growth rate. The rates of substrate transport differ between these transporter species. To compute the continuum growth rate estimate (grey line), we used the following transport rates for each transporter species: 200 glucose S^−1^ (BNID: 103693), 2000 glycerol S^−1^ (***Lu et al., 2003***), 200 fructose.S^−1^ (assumed to be similar to PtsI, BNID: 103693), and 50 xylose S^−1^ (assumed to be comparable to LacY, BNID:103159). Red points and highlighted text indicate conditions in which the only source of carbon in the growth medium induces expression of the transport system. Grey lines in (A) and (B) represents the estimated number of transporters per cell at a continuum of growth rates. **Figure 2 - Figure supplement 1**. Estimates and observed abundances of phosphate and sulfate transporters.

We find, however, that the experimental measurements exceed this by several fold (***Figure 2***(A), bottom), implying that the cell is capable of transporting more carbon atoms than strictly needed for biosynthesis. We can also abstract this calculation to consider any particular growth rate given knowledge of the cell density and volume as a function of growth rate (described further in the Appendix Section “Extending Estimates to a Continuum of Growth Rates”). This abstraction, shown as a grey line in ***Figure 2*(A)**, reveals an excess of transporters even at faster growth rates. This contrasts with our observations for uptake of phosphorus and sulfur, which turn out to align well with our expectations across different growth conditions (***Figure 2-Figure Supplement 1*** and discussed further in the Appendix Section “Additional Estimates of Fundamental Biological Processes”).

It is important to note that so far we have neglected any specifics of the regulation of the carbon transport system. Using the diverse array of growth conditions available in the data, we can explore how individual carbon transport systems depend on specific carbon availability. In ***Figure 2***(B), we show the total number of carbohydrate transporters specific to different carbon sources. A striking observation, shown in the top-left plot of ***Figure 2***(B), is the constancy in the expression of the glucose-specific transport systems, an observation that stands in contrast with other species of transporters. Additionally, we note that the total number of glucose-specific transporters is tightly distributed at ≈ 1 × 10^4^ per cell, the approximate number of transporters needed to sustain rapid growth of several divisions per hour. Interestingly, this illustrates that *E. coli* maintains a substantial number of complexes present for transporting glucose regardless of growth condition, which is known to be the preferential carbon source (***Monod, 1947***; ***Liu et al., 2005***; ***Aidelberg et al., 2014***).

Many metabolic operons are regulated with dual-input logic gates that are only expressed when glucose concentrations are low and the concentration of other carbon sources are elevated (***Gama-Castro et al., 2016***; ***Zhang et al., 2014***; ***Gama-Castro et al., 2016***; ***Belliveau et al., 2018***; ***Ireland et al., 2020***). Points colored in red in ***Figure 2***(B) (labeled by red text-boxes) correspond to growth conditions in which the specific carbon source (glycerol, xylose, or fructose) is present as the sole source of carbon. The grey lines in ***Figure 2***(B) show the estimated number of transporters needed at each growth rate to satisfy the cellular carbon requirement, adjusted for the specific carbon source in terms of number of carbon atoms per molecule and the rate of transport for the particular transporter species. These plots show that, even in the absence of the particular carbon source, expression of the transporters is maintained on the order of ≈ 1 × 10^2^ per cell. The low but non-zero abundances may reflect the specific regulatory logic involved, requiring that cells are able to transport some minimal amount of an alternative carbon source in order to induce expression of these alternative carbon-source systems when needed (***Laxhuber et al., 2020***).

### Limits on Transporter Expression

If acquisition of nutrients was a limiting process in cell division under the typical growth conditions explored here, the growth rate could be theoretically increased simply by expressing more transporters, but is this feasible at a physiological level? A way to approach this question is to compute the amount of space in the bacterial membrane that could be occupied by nutrient transporters. Considering a rule-of-thumb for the surface area of *E. coli* of about 5 μm^2^ (BNID: 101792), we expect an areal density for 2000 transporters to be approximately a few hundred transporters per μm^2^. For a typical transporter occupying about 50 nm2, this amounts to about only ≈ 1% of the total inner membrane surface area (***Szenk et al., 2017***). In contrast, bacterial cell membranes typically have densities of ≈ 1 × 10^5^ proteins/μm^2^ (***Phillips, 2018***), with roughly 60 % of the surface area occupied by protein (BNID: 100078), implying that the cell could easily accommodate more transporters. There are, however, additional constraints on the space that can be devoted to nutrient uptake due to occupancy by proteins involved in processes like cell wall synthesis and energy production, and we will consider this further in the coming sections.

## Cell Envelope Biogenesis

In contrast to nutrient transporters, which support the synthesis of biomolecules throughout the cell and therefore need to scale with the cell size, here we must consider the synthesis of components that will need to scale with the surface area of the cell. *E. coli* is a rod-shaped bacterium with a remarkably robust length-to-width aspect ratio of ≈ 4:1 (***Harris and Theriot, 2018***; ***Ojkic et al., 2019***). Assuming this surface area is approximately the same between the inner and outer membranes of *E. coli*, and the fact that each membrane is itself is a lipid bilayer (or, a bilayer with lipopolysaccharides decorating the outer membrane), our rule-of-thumb of 5 μm^2^ per surface suggests a total membrane surface area of ≈ 20μm^2^ (see the Appendix Section “Estimation of Cell Size and Surface Area” for a description of the calculation of cell surface area as a function of cell size). In this section, we will estimate the number of key protein complexes needed to synthesize the lipids as well as the complexes involved in assembling the peptidoglycan scaffold that makes up the cell envelope.

### Lipid Synthesis

The dense packing of the membrane with proteins means that the cell membranes are not composed entirely of lipid molecules, with only ≈ 40 % of the membrane area occupied by lipids or lipopolysaccharide, both of which have fatty acid chains of similar length (BNID: 100078). Using a rule-of-thumb of 0.5 nm2 as the surface area of the typical lipid (BNID: 106993), we can estimate ≈ 2 × 10^7^ lipids per cell, which is in close agreement with experimental measurements (BNID: 100071, 102996).

The membranes of *E. coli* are composed of a variety of different lipids, each of which are unique in their structures and biosynthetic pathways (***Sohlenkamp and Geiger, 2016***). Recently, a combination of stochastic kinetic modeling (***Ruppe and Fox, 2018***) and *in vitro* kinetic measurements (***Ranganathan et al., 2012***; ***Yu et al., 2011***) has revealed remarkably slow steps in the fatty acid synthesis pathways which may serve as the rate limiting reactions for making new membrane fatty acids (that become components of a variety of membrane lipids) in *E. coli*. One such step is the removal of hydroxyl groups from the fatty-acid chain by ACP dehydratase that leads to the formation of carbon-carbon double bonds. This reaction, catalyzed by proteins FabZ and FabA (***Yu et al., 2011***), has been estimated to have kinetic turnover rates of ≈ 1 dehydration per second per enzyme (***Ruppe and Fox, 2018***). Thus, given this rate and the need to synthesize ≈ 2 × 10^7^ lipids over 5000 seconds, one can estimate that a typical cell requires ≈ 4000 ACP dehydratases. This is in reasonable agreement with the experimentally observed copy numbers of FabZ and FabA (***Figure 3***(A)). Furthermore, we can extend this estimate to account for the change in membrane surface area as a function of the growth rate (grey line in ***Figure 3***(A)), which in contrast to our observations with glucose uptake, indeed captures the observed growth rate dependent expression of these two enzymes.

**Figure 3.**
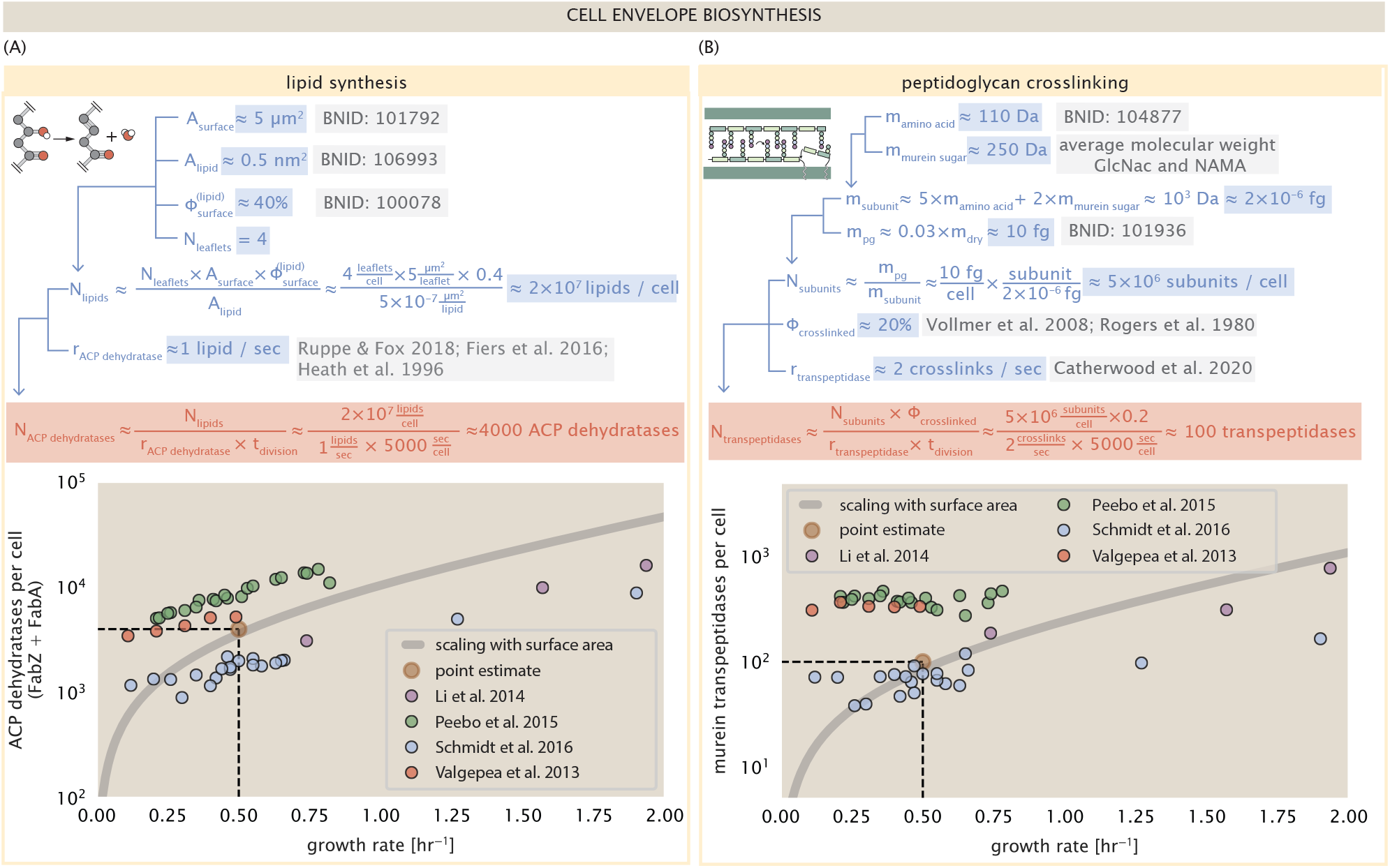
(A) Top panel shows an estimation for the number of ACP dehydratases necessary to form functional phospholipids, which is assumed to be a rate-limiting step on lipid synthesis. The rate of ACP dehydratases was inferred from experimental measurements via a stochastic kinetic model described in ***Ruppe and Fox (2018)***. Bottom panel shows the experimentally observed complex copy numbers using the stoichiometries [FabA]_2_ and [FabZ]_2_. (B) An estimate for the number of peptidoglycan transpeptidases needed to complete maturation of the peptidoglycan. The mass of the murein subunit was estimated by approximating each amino acid in the pentapeptide chain as having a mass of ≈ 110 Da and each sugar in the disaccharide having a mass of ≈ 250 Da. The *in vivo* rate of transpeptidation in *E. coli* was taken from recent analysis by Catherwood et al. (2020). The bottom panel shows experimental measurements of the transpeptidase complexes in *E. coli* following the stoichiometries [MrcA]_2_, [MrcB]_2_, [MrdA]_1_, and [MrdB]_1_. Grey curves in each plot show the estimated number of complexes needed to satisfy the synthesis requirements scaled by the surface area as a function of growth rate.

### Peptidoglycan Synthesis

The exquisite control of bacteria over their cell shape is due primarily to a stiff, several nanometer thick meshwork of polymerized disaccharides that makes up the cell wall termed the peptidoglycan. The formation of the peptidoglycan is an intricate process involving many macromolecular players (***Shi et al., 2018***; ***Morgenstein et al., 2015***), whose coordinated action synthesizes the individual subunits and integrates them into the peptidoglycan network that maintains cell shape and integrity even in the face of large-scale chemical and osmotic perturbations (***Harris and Theriot, 2018***; ***Shi et al., 2018***). Due to the extensive degree of chemical crosslinks between glycan strands, the entire peptidoglycan is a single molecule comprising ≈ 3% of the cellular dry mass (BNID: 1019360), making it the most massive molecule in *E. coli*. The polymerized unit of the peptidoglycan is a N-acetylglucosamine and N-acetylmuramic acid disaccharide, of which the former is functionalized with a short pentapeptide. With a mass of ≈ 1000 Da, this unit, which we refer to as a murein subunit, is polymerized to form long strands in the periplasm which are then attached to each other via their peptide linkers. Together, these quantities provide an estimate of ≈ 5 × 10^6^ murein subunits per cell.

There are various steps which one could consider *a priori* to be a limiting process in the synthesis of peptidoglycan, including the biosynthesis steps that occur in the cytoplasm, the transglycosylation reaction which adds new subunits to the glycan strands, and the formation of the peptide crosslinks between strands (***Shi et al., 2018***; ***Morgenstein et al., 2015***; ***Lovering et al., 2012***; ***Barreteau et al., 2008***). Despite the extensive mechanistic characterization of these components, *quantitative* characterization of the individual reaction rates along the entire kinetic pathway of remain scarce and make identification of any particularly slow steps difficult. However, such measurements have recently been made for the crosslinking machinery [transpeptidases, ***Catherwood et al.*** (***2020***)] of the peptidodglycan which provides lateral structural integrity to the peptidoglycan shell. As the primary mechanism of subunit integration occurs by a complex with both transglycosylation and transpeptidation activities (***Shi et al., 2018***) and that the measured turnover of transpeptidases being rather slow (≈ 2 crosslinking reactions per second) we therefore consider only the transpeptidation reaction in this work. We believe that, in lieu of other quantitative measurements, crosslinking represents a reasonable candidate for a rate-limiting step in growth as it is vital for cell size and shape homeostasis.

In principle, each murein subunit can be involved in such a crosslink. In some microbes, such as in Gram-positive bacterium *Staphylococcus aureus*, the extent of crosslinking can be large with > 90% of pentapeptides forming a connection between glycan strands. In *E. coli*, however, a much smaller proportion (≈ 20%) of the peptides are crosslinked, resulting in a weaker and more porous cell wall (***Vollmer et al., 2008***; ***Rogers et al., 1980***). The formation of these crosslinks occurs primarily during the polymerization of the murein subunits and is facilitated by a family of transpeptidase enzymes. The four primary transpeptidases of *E. coli* have only recently been quantitatively characterized *in vivo*, via liquid chromatography mass spectrometry, which revealed a notably slow kinetic turnover rate of ≈ 2 crosslinking reactions formed per second per enzyme as noted above (***Catherwood et al., 2020***).

Assembling these quantities permits us to make an estimate that on the order of ≈ 100 transpeptidases per cell are needed for complete maturation of the peptidoglycan, given a division time of ≈ 5000 seconds; a value that is comparable to experimental observations [***Figure 3***(B)]. Expanding this estimate to account for the changing mass of the peptidoglycan as a function of growth rate [grey line in ***Figure 3***(B)] predicts an order-of-magnitude increase in the abundance of the transpeptidases when the grow rate is increased by a factor of four. Here, however, the measured complex abundances across the different proteomic data sets show systematic disagreements and obfuscates any significant dependence on growth rate.

### Limits on Cell Wall Biogenesis

While the processes we have considered represent only a small portion of proteins devoted to cell envelope biogenesis, we find it unlikely that they limit cellular growth in general. The relative amount of mass required for lipid and peptidoglycan components will decrease at faster growth rates due to a decrease in the cell’s surface area to volume ratio (***Ojkic et al., 2019***). Furthermore, despite the slow catalytic rate of FabZ and FabA in lipid synthesis, experimental data and recent computational modeling has shown that the rate of fatty-acid synthesis can be drastically increased by increasing the concentration of FabZ (***Yu et al., 2011***; ***Ruppe and Fox, 2018***). With a proteome size of ≈ 3×10^6^ proteins, a hypothetical 10-fold increase in expression from 4000 to 40,000 ACP dehydratases would result in a paltry ≈ 1% increase in the size of the proteome. In the context of peptidoglycan synthesis, we note that our estimate considers only the transpeptidase enzymes that are involved in lateral and longitudinal elongation of the peptidoglycan. This neglects the presence of other transpeptidases that are present in the periplasm and also involved in remodeling and maturation of the peptidoglycan. It is therefore possible that if this was setting the speed limit for cell division, the simple expression of more transpeptidases would be sufficient to maintain the structural integrity of the cell wall.

## Energy Production

Cells consume and generate energy predominantly in the form of nucleoside triphosphates (NTPs) in order to grow. The highenergy phosophodiester bonds of (primarily) ATP power a variety of cellular processes that drive biological systems away from thermodynamic equilibrium. We therefore turn to the synthesis of ATP as a potential process that may limit growth, which will also require us to consider the maintenance of the electrochemical proton gradient that powers it.

### ATP Synthesis

Hydrolysis of the terminal phosphodiester bond of ATP into ADP (or alternatively GTP and GDP) and an inorganic phosphate provides the thermodynamic driving force in a wide array of biochemical reactions. One such reaction is the formation of peptide bonds during translation, which requires ≈ 2 ATPs for the charging of an amino acid to the tRNA and ≈ 2 GTPs for the formation of each peptide bond. Assuming the ATP costs associated with error correction and post-translational modifications of proteins are negligible, we can make the approximation that each peptide bond has a net cost of ≈ 4 ATP (BNID: 101442). Formation of GTP from ATP is achieved via the action of nucleoside diphosphate kinase, which catalyzes this reaction without an energy investment (*Lascu and Gonin, 2000*). We therefore consider all NTP requirements of the cell to be functionally equivalent to being exclusively ATP. In total, the energetic costs of peptide bond formation consumes ≈ 80% of the cells ATP budget [BNID: 107782; 106158; 101637; 111918, ***Lynch and Marinov*** (***2015***); ***Stouthamer*** (***1973***)]. This pool of ATP is primarily produced by the F_1_-F_0_ ATP synthase – a membrane-bound rotary motor which under ideal conditions can yield ≈ 300 ATP per second [BNID: 114701; ***Weber and Senior*** (***2003***)].

To estimate the total number of ATP equivalents consumed during a cell cycle, we will make the approximation that there are ≈ 3 × 10^6^ proteins per cell with an average protein length of ≈ 300 peptide bonds (BNID: 115702; 108986; 104877). Taking these values together, coupled with an estimate of ≈ 4 ATP equivalents per peptide bond, we find that the typical *E. coli* cell consumes ≈ 5 × 109 ATP per cell cycle on protein synthesis alone. Assuming that each ATP synthases operates at its maximal speed (300 ATP per second per synthase), ≈ 3000 ATP synthases are needed to keep up with the energy demands of the cell. This estimate is comparable with the experimental observations, shown in ***Figure 4***(A). Since this estimate assumes all ATP is synthesized via ATP synthase and neglects synthesis via fermentative metabolism, this may explain why at the fastest growth rates (≈ 2 hr^−1^), our continuum estimate predicts more synthase than is experimentally observed (data points below the gray line in ***Figure 4***(A) at fast growth rates). In particular, at faster growth rates, *E. coli* enters a type of overflow metabolism where non-respiratory routes for ATP synthesis become more pronounced and provide the remaining ATP demand (***Molenaar et al., 2009***; ***Zhuang et al., 2011***; ***Szenk et al., 2017***).

**Figure 4.**
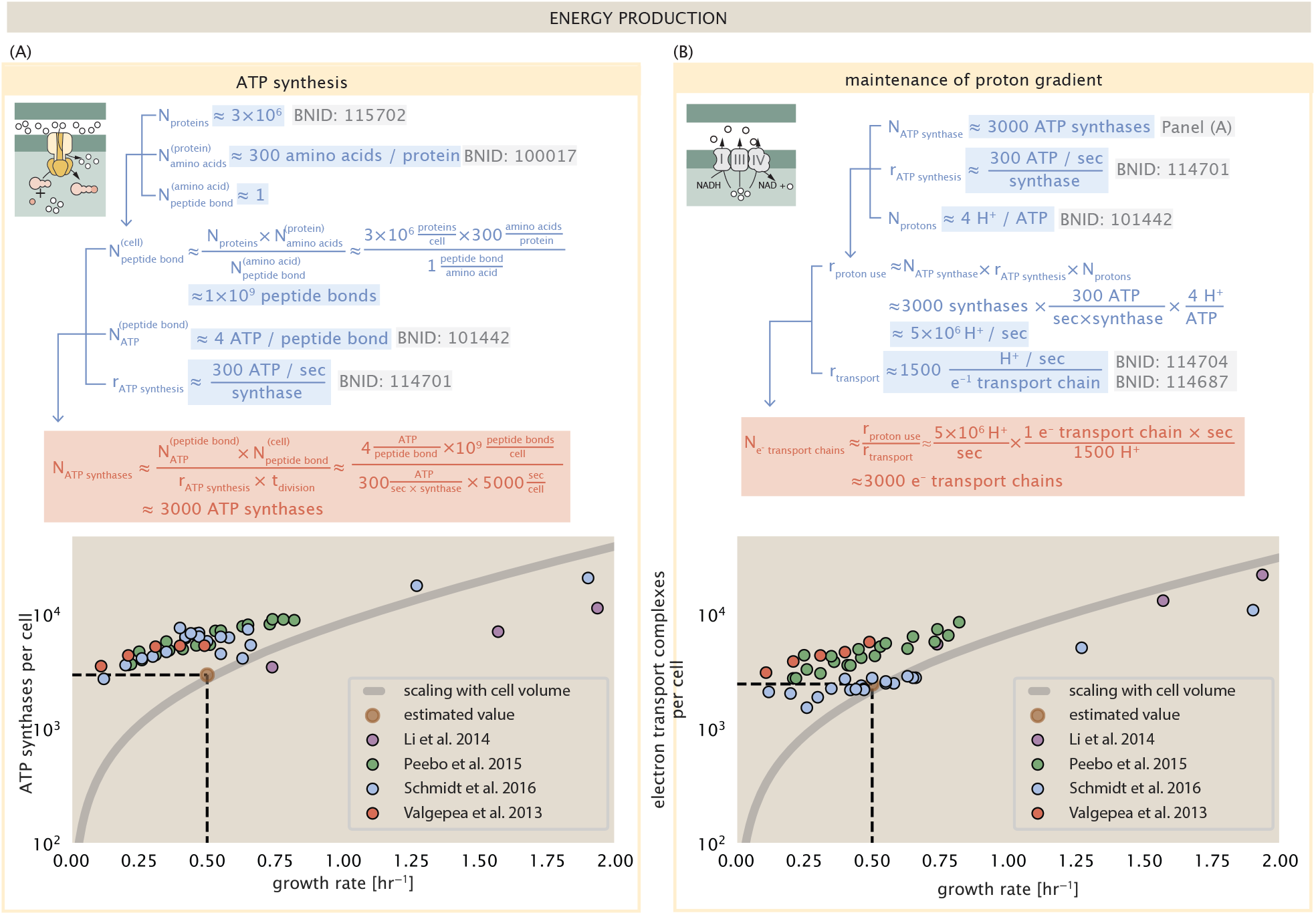
The abundance of F_1_-F_0_ ATP synthases and electron transport chain complexes as a function of growth rate. (A) Estimate of the number of F_1_-F_0_ ATP synthase complexes needed to accommodate peptide bond formation and other NTP dependent processes. Points in plot correspond to the mean number of complete F_1_-F_0_ ATP synthase complexes that can be formed given proteomic measurements and the subunit stoichiometry [AtpE]_10_[AtpF]_2_[AtpB][AtpC][AtpH][AtpA]_3_[AtpG][AtpD]_3_. (B) Estimate of the number of electron transport chain complexes needed to maintain a membrane potential of −200 mV given estimate of number of F_1_-F_0_ ATP synthases from (A). Points in plot correspond to the average number of complexes identified as being involved in aerobic respiration by the Gene Ontology identifier GO:0019646 that could be formed given proteomic observations. These complexes include cytochromes *bd1* ([CydA][CydB][CydX][CydH]), *bdII* ([AppC][AppB]), *bo*_3_,([CyoD][CyoA][CyoB][CyoC]) and NADH:quinone oxioreducase I ([NuoA][NuoH][NuoJ][NuoK][NuoL][NuoM][NuoN][NuoB][NuoC][NuoE][NuoF][NuoG][NuoI]) and II ([Ndh]). Grey lines in both (A) and (B) correspond to the estimate procedure described, but applied to a continuum of growth rates. We direct the reader to the Supporting Information for a more thorough description of this approach.

### Generating the Proton Electrochemical Gradient

In order to produce ATP, the F_1_-F_0_ ATP synthase itself must consume energy. Rather than burning through its own product (and violating thermodynamics), this intricate macromolecular machine has evolved to exploit the electrochemical potential established across the inner membrane through cellular respiration. This electrochemical gradient is manifest by the pumping of protons into the intermembrane space via the electron transport chains as they reduce NADH. In *E. coli*, this potential difference is ≈ −200 mV (BNID: 102120). A simple estimate of the inner membrane as a capacitor with a working voltage of −200 mV reveals that ≈ 2 × 10^4^ protons must be present in the intermembrane space. However, each rotation of an ATP synthase shuttles ≈ 4 protons into the cytosol (BNID: 103390). With a few thousand ATP synthases producing ATP at their maximal rate, the potential difference would be rapidly abolished in a few milliseconds if it were not being actively maintained.

The electrochemistry of the electron transport complexes of *E. coli* have been the subject of intense biochemical and biophysical study (***Ingledew and Poole, 1984***; ***Khademian and Imlay, 2017***; ***Cox et al., 1970***; ***Henkel et al., 2014***). A recent work (***Szenk et al., 2017***) examined the respiratory capacity of the *E. coli* electron transport complexes using structural and biochemical data, revealing that each electron transport chain rapidly pumps protons into the intermembrane space at a rate of ≈ 1500 protons per second (BIND: 114704; 114687). Using our estimate of the number of ATP synthases required per cell [***Figure 4***(A)], coupled with these recent measurements, we estimate that ≈ 3000 electron transport complexes would be necessary to facilitate the ≈ 5 × 10^6^ protons per second diet of the cellular ATP synthases. This estimate is in agreement with the number of complexes identified in the proteomic datasets [plot in ***Figure 4***(B)]. This suggests that every ATP synthase must be accompanied by ≈ 1 functional electron transport chain.

### Limits on Biosynthesis in a Crowded Membrane

Our estimates thus far have focused on biochemistry at the periphery of the cell, with the processes of nutrient transport, cell envelope biogenesis, and energy generation all requiring space to perform their biological functions. The cell’s surface area, however, does not scale as rapidly as cell size (***Harris and Theriot, 2018***) and there will be diminishing space available to support the proteomic requirements at faster growth rates. It is therefore necessary to consider the consequences of a changing surface area to volume ratio in our effort to identify limitations on growth. Here we use our analysis of ATP production to better understand this constraint.

In our estimate of ATP production above we found that a cell demands about 5 × 10^9^ ATP per cell cycle or ≈ 1 × 10^6^ ATP/s. With a cell volume of roughly 1 fL (BNID: 100004), this corresponds to about 2×10^10^ ATP per fL of cell volume, in line with previous estimates (***Stouthamer and Bettenhaussen, 1977***; ***Szenk et al., 2017***). In ***Figure 5*** (A) we plot this ATP demand as a function of the surface area to volume ratio in green, where we have considered a range of cell shapes from spherical to rod-shaped with an aspect ratio (length/width) equal to 4. In order to consider the maximum ATP that could be produced, we consider the amount of ATP that can be generated by a membrane filled with ATP synthase and electron transport complexes and a maximal production rate of about 3 ATP/ (nm2 s) (***Szenk et al., 2017***). This is shown in blue in ***Figure 5***(A), which shows that at least for the growth rates observed (right column in plot), the energy demand is roughly an order of magnitude less. Interestingly, ***Szenk et al.*** (***2017***) found that ATP production by respiration is less efficient than by fermentation on a per membrane area basis, due to the additional proteins of the electron transport chain. This suggests that, even under anaerobic growth, cells will have sufficient membrane space for ATP production.

**Figure 5.**
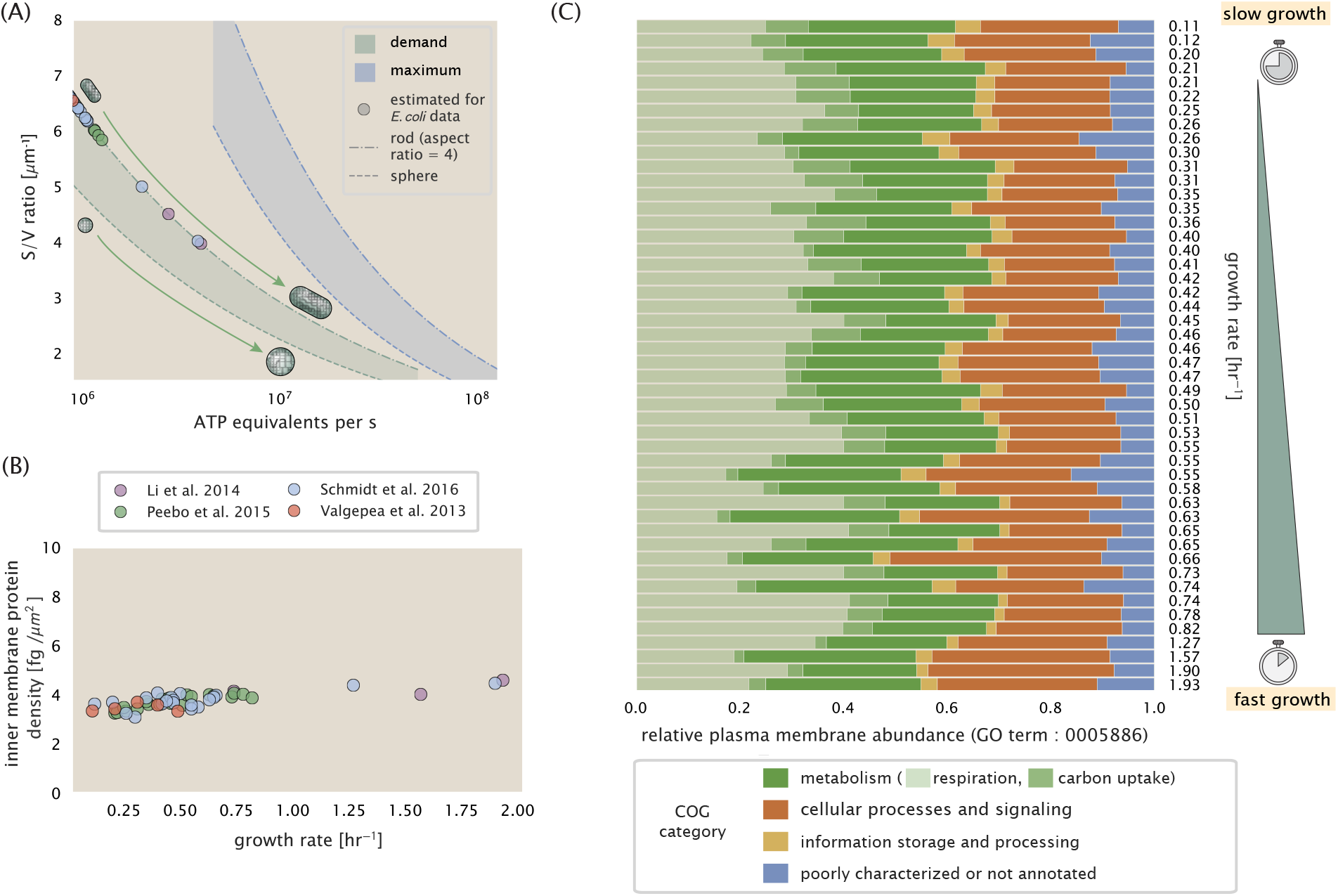
Inluence of cell size and surface area to volume ratio on ATP production and inner membrane composition. (A) Scaling of ATP demand and maximum ATP production through respiration as a function of surface area to volume ratio. Cell volumes of 0.5 fL to 50 fL were considered, with the dashed (- -) line corresponding to a sphere and the dash-dot line (-.) reflecting a rod-shaped bacterium like *E. coli* with a typical aspect ratio (length / width) of 4 (***Shi et al., 2018***). The ATP demand is calculated as 106 ATP/(μm^3^ s), while the maximum ATP production rate is taken to be 3 ATP / (nm^2^·s) (***Szenk et al., 2017***), with calculations of *E. coli* volume and surface area detailed in Appendix Section “Estimation of Cell Size and Surface Area”. In this calculation, 50% of the bacterial inner membrane is assumed to be protein, with the remainder lipid. (B) Total protein mass per μm^2^ calculated for proteins with inner membrane annotation (GO term: 0005886). (C) Relative protein abundances are grouped by their COG annotations (‘metabolic’, ‘cellular processes and signaling’, ‘information storage and processing’, and ‘poorly characterized or not annotated‘) for the data from ***Schmidt et al.***(***2016***). Metabolic proteins are further separated into respiration (F_1_-F_0_ ATP synthase, NADH dehydrogenase I, succinate:quinone oxidoreductase, cytochrome bo_3_ ubiquinol oxidase, cytochrome bd-I ubiquinol oxidase) and carbohydrate transport (GO term: GO:0008643). Note that the elongation factor EF-Tu can also associate with the inner membrane, but was excluded in this analysis due to its high relative abundance (roughly identical to the summed protein shown in part (B)).

Importantly, this analysis highlights that there will indeed be a maximum attainable cell size due to the limited capacity to provide resources as the cell increases in size. The maximum energy production in ***Figure 5***(A), however, does represent a somewhat unachievable limit since the inner membrane also includes other proteins like those we have considered for nutrient transport and cell wall biogenesis. To better understand the overall proteomic makeup of the inner membrane, we therefore used Gene Ontology (GO) annotations (***Ashburner et al., 2000***; ***The Gene Ontology Consortium, 2018***) to identify all proteins embedded or peripheral to the inner membrane (GO term: 0005886). Those associated but not membrane-bound include proteins like MreB and FtsZ that must nonetheless be considered as a vital component occupying space on the membrane. In ***Figure 5***(B), we find that the total protein mass per μm^2^ is nearly constant across growth rates. Interestingly, when we consider the distribution of proteins grouped by their Clusters of Orthologous Groups (COG) (***Tatusov et al., 2000***), the relative abundance of each category is nearly constant across growth rates. This suggests that no one process (energy production, nutrient uptake, etc.) is dominating even at fast growth rates [***Figure 5***(C)] and in line with our supposition that each of the processes we’ve considered so far are not fundamentally limiting the maximum growth rate. In contrast, when we apply such an analysis to cytosolic proteins (GO term: 0005829), we observe a clear change in the proteomic composition [***Figure 6***(A, B)]. In particular, with increasing growth rates there is a substantial increase in the relative protein mass associated with ‘information storage and processing’. This category includes proteins such as DNA polymerase, RNA polymerase, and ribosomes that are associated with the processes of the central dogma, whose increase is predominantly at the expense of ‘metabolic’ proteins as shown in ***Figure 6***(C). This notable anticorrelation shown in ***Figure 6***(D) is consistent with previous reports (***Schmidt et al., 2016***; ***Scott et al., 2010***; ***Zhu and Dai, 2019***). In the next section we therefore turn our attention to the processes of the central dogma.

**Figure 6.**
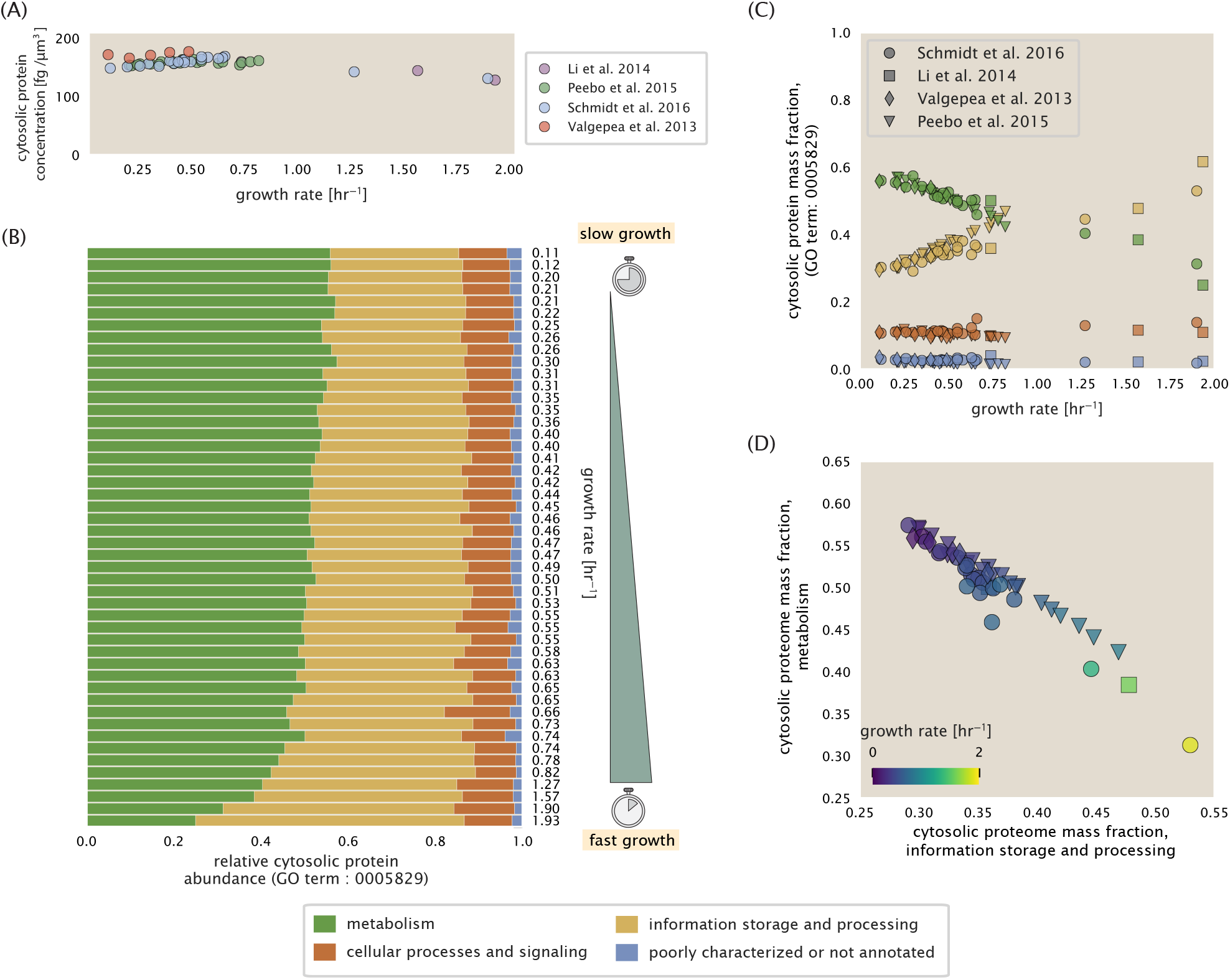
Characterization of cytosolic proteomic composition. (A) Total protein mass per μm^3^ calculated for cytosolic proteins (GO term: 0005886), with calculations of *E. coli* volume detailed in Appendix Section “Estimation of Cell Size and Surface Area”. (B) The distribution of cytosolic proteins is compared across the different growth conditions in ***Schmidt et al.*** (***2016***) by grouping their relative abundances by COG annotation (‘metabolic’, ‘cellular processes and signaling’, ‘information storage and processing’, ‘poorly characterized or not annotated’). (C) Relative cytosolic protein abundances, grouped by their COG annotations, are plotted as a function of growth rate. (D) The relative protein abundances associated with the ‘information storage and processing’ and ‘metabolic’ COG categories are plotted against each other and highlight that faster growth rates are associated with a larger mass fraction devoted to the ‘information storage and processing’ proteins of the central dogma.

## Processes of the Central Dogma

Up to this point, we have considered a variety of transport and biosynthetic processes that are critical to acquiring and generating new cell mass. While there are of course many other metabolic processes we could consider, we now turn our focus to some of the most important processes which *must* be undertaken irrespective of the growth conditions – those of the central dogma.

### DNA Replication

Most bacteria (including *E. coli*) harbor a single, circular chromosome and can have extra-chromosomal plasmids up to Ì 100 kbp in length. While we consider the starting material dNTPs in ***Figure 7-Figure Supplement 1*** and discussed further in the Appendix Section “Additional Process of the Central Dogma”, here we focus our quantitative thinking on the chromosome of *E. coli*, which harbors ≈ 5000 genes and ≈ 5 × 10^6^ base pairs.

To successfully divide and produce viable progeny, this chromosome must be faithfully replicated and segregated into each nascent cell. Replication is initiated at a single region of the chromosome termed the *oriC* locus where a pair of replisomes, each consisting of two DNA polymerase III, begin their high-fidelity replication of the genome in opposite directions (***Fijalkowska et al., 2012***). *In vitro* measurements have shown that DNA Polymerase III copies DNA at a rate of ≈ 600 nucleotides per second (BNID: 104120). Therefore, to replicate a single chromosome, two replisomes moving at their maximal rate would copy the entire genome in ≈ 4000 s. Thus, with a division time of 5000 seconds, there is sufficient time for a pair of replisome complexes to replicate the entire genome.

In rapidly growing cultures, bacteria like *E. coli* can initiate as many as 10 - 12 replication forks at a given time (***Bremer and Dennis, 2008***; ***Si et al., 2017***), we expect only a few DNA polymerases (≈ 10) are needed. However, as shown in ***Figure 7***, DNA polymerase III is nearly an order of magnitude more abundant. This discrepancy can be understood by considering its binding constant to DNA. *In vitro* characterization has quantified the *K*_*D*_ of DNA polymerase III holoenzyme to single-stranded and double-stranded DNA to be 50 and 200 nM, respectively (***Ason et al., 2000***). The right-hand plot in ***Figure 7*** shows that the concentration of DNA polymerase III across all data sets is within this range. Thus, its copy number appears to vary such that its concentration is approximately equal to the dissociation constant to the DNA. While the processes regulating the initiation of DNA replication are complex and involve more than just the holoenzyme, these data indicate that the kinetics of replication rather than the explicit copy number of the DNA polymerase III holoenzyme is the more relevant feature of DNA replication to consider. In light of this, the data in ***Figure 7*** suggests that for bacteria like *E. coli*, DNA replication does not represent a rate-limiting step in cell division. However, it is worth noting that for bacterium like *C. crescentus* whose chromosomal replication is initiated only once per cell cycle (***Jensen et al., 2001***), the time to double their chromosome indeed represents an upper limit to their growth rate.

**Figure 7.**
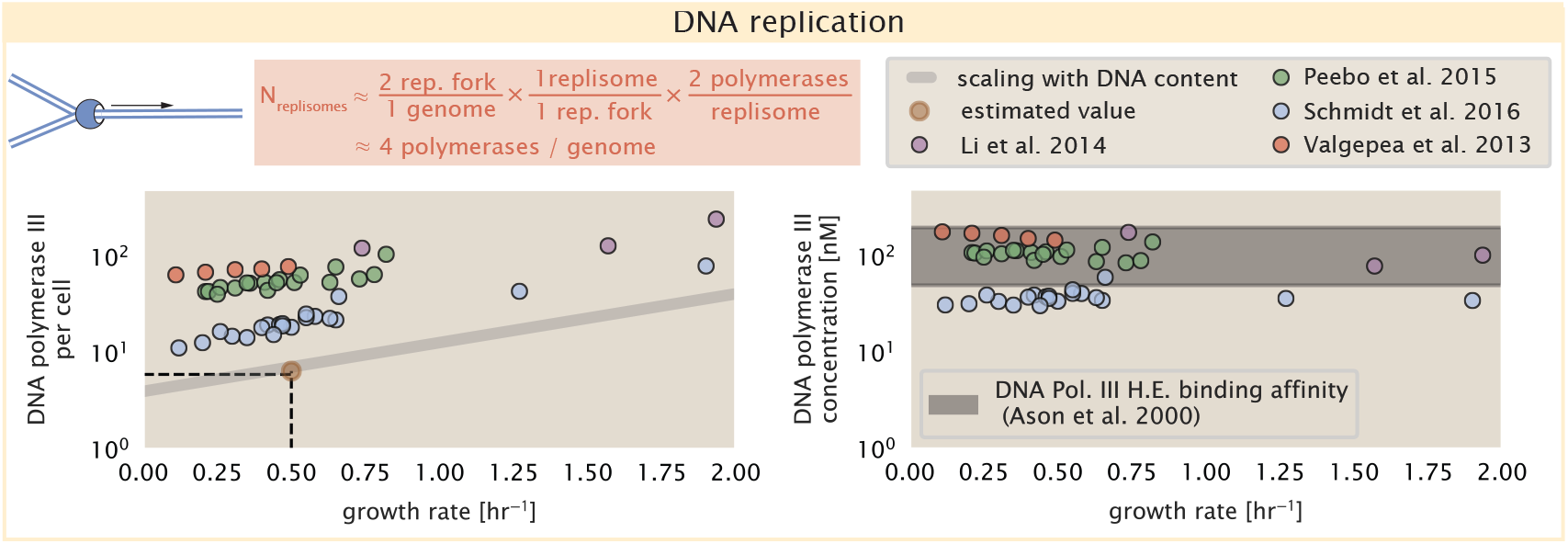
Complex abundance estimates for dNTP synthesis and DNA replication. An estimate for the minimum number of DNA polymerase holoenzyme complexes needed to facilitate replication of a single genome. Points in the left-hand plot correspond to the total number of DNA polymerase III holoenzyme complexes ([DnaE]_3_[DnaQ]_3_[HolE]_3_[DnaX]_5_[HolB][HolA][DnaN]_4_[HolC]_4_[HolD]_4_) per cell. Right-hand plot shows the effective concentration of DNA polymerase III holoenzyme (See Appendix Section “Estimation of Cell Size and Surface Area” for calculation of cell size). Grey lines in left-hand panel show the estimated number of complexes needed as a function of growth, the details of which are described in the Appendix. Figure 7-Figure supplement 1. Estimate and observations of the abundance of ribonucleotide reductase, a key component in dNTP synthesis.

### RNA Synthesis

We now turn our attention to the next stage of the central dogma – the transcription of DNA to form RNA. We consider three major groupings of RNA, namely the RNA associated with ribosomes (rRNA), the RNA encoding the amino-acid sequence of proteins (mRNA), and the RNA which links codon sequence to amino-acid identity during translation (tRNA).

rRNA serves as the catalytic and structural component of the ribosome, comprising approximately 2/3 of the total ribosomal mass, and is decorated with ≈ 50 ribosomal proteins. Each ribosome contains three rRNA molecules of lengths 120, 1542, and 2904 nucleotides (BNID: 108093), meaning each ribosome contains ≈ 4500 nucleotides overall. *In vivo* measurements of the kinetics of rRNA transcription have revealed that RNA polymerases are loaded onto the promoter of an rRNA gene at a rate of ≈ 1 per second (BNID: 111997, 102362). If RNA polymerases are constantly loaded at this rate, then we can assume that ≈ 1 functional rRNA unit is synthesized per second per rRNA operon. While *E. coli* possesses 7 of these operons per chromosome, the fact that chromosome replication can be parallelized means that the average dosage of rRNA genes can be substantially higher (up to ≈ 70 copies) at fast growth rates (***Dennis et al., 2004***). At a growth rate of ≈ 0.5 hr^−1^, however, the average cell has ≈ 1 copy of its chromosome and therefore approximately ≈ 7 copies of the rRNA operons, producing ≈ 7 rRNA units per second. With a 5000 second division time, this means the cell is able to generate around 3 × 10^4^ functional rRNA units, comparable within an order of magnitude to the number of ribosomes per cell.

How many RNA polymerases are then needed to constantly transcribe the required rRNA? If one polymerase is loaded once every two seconds on average (BNID: 111997), and the transcription rate is ≈ 40 nucleotides per second (BNID: 101094), then the typical spacing between polymerases will be ≈ 40 nucleotides. However, we must note that the polymerase itself has a footprint of ≈ 40 nucleotides (BNID: 107873), meaning that one could expect to find one RNA polymerase per 80 nucleotide stretch of an rRNA gene. With a total length of ≈ 4500 nucleotides per operon and 7 operons per cell, the number of RNA polymerases transcribing rRNA at any given time is then ≈ 500 per cell.

As outlined in ***Figure 8***, and discussed further the Appendix Section “Additional Process of the Central Dogma”, synthesis of mRNA and tRNA together require on the order of another ≈ 400 RNAP. Thus, in total, one would expect the typical cell to require ≈ 1000 RNAP to satisfy its transcriptional demands. As is revealed in ***Figure 8***(B), this estimate is about an order of magnitude below the observed number of RNA polymerase complexes per cell (≈ 5000 - 7000). The difference between the estimated number of RNA polymerase needed for transcription and these observations, however, are consistent with literature revealing that ≈ 80% of RNA polymerases in *E. coli* are not transcriptionally active (***Patrick et al., 2015***).

**Figure 8.**
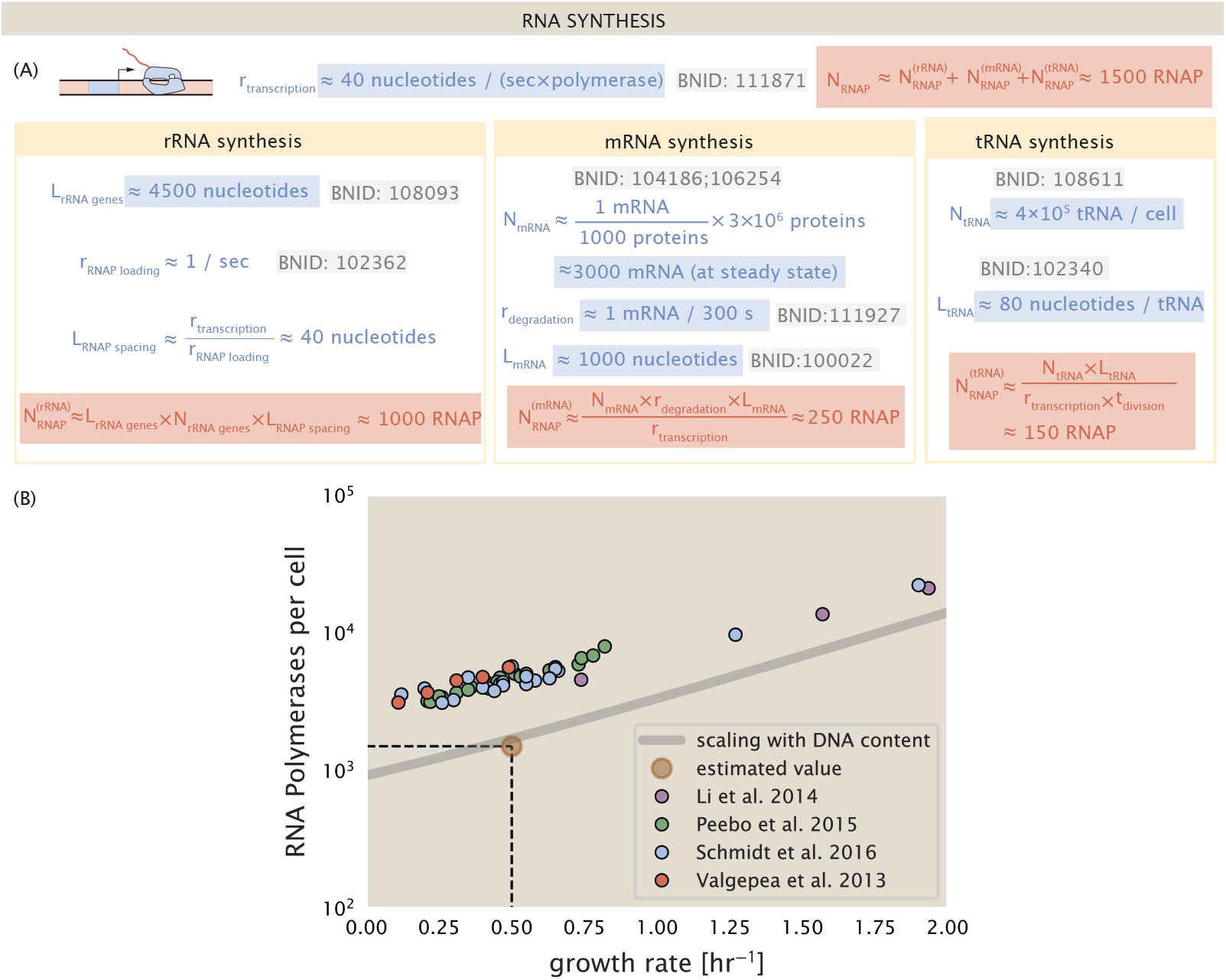
Estimation of the RNA polymerase demand and comparison with experimental data. (A) Estimations for the number of RNA polymerase needed to synthesize sufficient quantities of rRNA, mRNA, and tRNA from left to right, respectively. (B) The RNA polymerase core enzyme copy number as a function of growth rate. Colored points correspond to the average number RNA polymerase core enzymes that could be formed given a subunit stoichiometry of [RpoA]_2_[RpoC][RpoB]. **Figure 8-Figure supplement 1. Abundance and growth rate dependence of σ-70**.

Our estimates also neglect other mechanistic features of transcription and transcriptional initiation more broadly. For example, we acknowledge that a major fraction of the RNAP pool is non-specifically bound to DNA during its search for promoters from which to begin transcription (***Klumpp and Hwa, 2008***). Furthermore, we ignore the obstacles that RNA polymerase and DNA polymerase present to each other as they move along the DNA (***Finkelstein and Greene, 2013***). Additionally, while they represent the core machinery for transcription, RNA polymerase is not sufficient to initiate transcription. Initiation of transcription is often dependenton the presence of C-factors, protein cofactors that bind directly to the polymerase (***Browning and Busby, 2016***) and aid in promoter recognition. In ***Figure 8-Figure Supplement 1***, we show that the predicted RNA polymerase copy number indeed is more comparable with the abundance of σ-70 (RpoD), the primary sigma factor in *E. coli*. There therefore remains more to be investigated as to what sets the observed abundance of RNA polymerase in these proteomic data sets. However, we conclude that the observed RNA polymerase abundances are generally in excess of what appears to be needed for growth, suggesting that the synthesis of RNA polymerase themselves are not particularly limiting.

### Protein Synthesis

We conclude our dialogue between back-of-the-envelope estimates and comparison with the proteomic data by examining the final process in the central dogma – translation. In doing so, we will begin with an estimate of the number of ribosomes needed to replicate the cellular proteome. While the rate at which ribosomes translate is well known to depend on the growth rate [***Dai et al.*** (***2018***), a phenomenon we consider later in this work] we begin by making the approximation that translation occurs at a modest rate of ≈ 15 amino acids per second per ribosome (BNID: 100233). Under this approximation and our previous estimate of 10^9^ peptide bonds per cell at a growth rate of 0.5 hr^−1^, we can easily arrive at an estimate of ≈ 10^4^ ribosomes needed per cell to replicate the entire protein mass [***Figure 9***(A, top)]. This point estimate, as well as the corresponding estimate across a continuum of growth rates, proves to be notably comparable to the experimental observations, shown in the bottom panel of ***Figure 9***(A). While the ribosome is responsible for the formation of peptide bonds, we do not diminish the importance of charging tRNAs with their appropriate amino acid, a process with occurs with remarkable fidelity. In ***Figure 9-Figure Supplement 1*** we consider the process of ligating tRNAs to their corresponding amino acid, with further details provided in the Appendix Section “Additional Estimates of Fundamental Biological Processes,” with notable accord observed between the data and our quantitative expectations.

**Figure 9.**
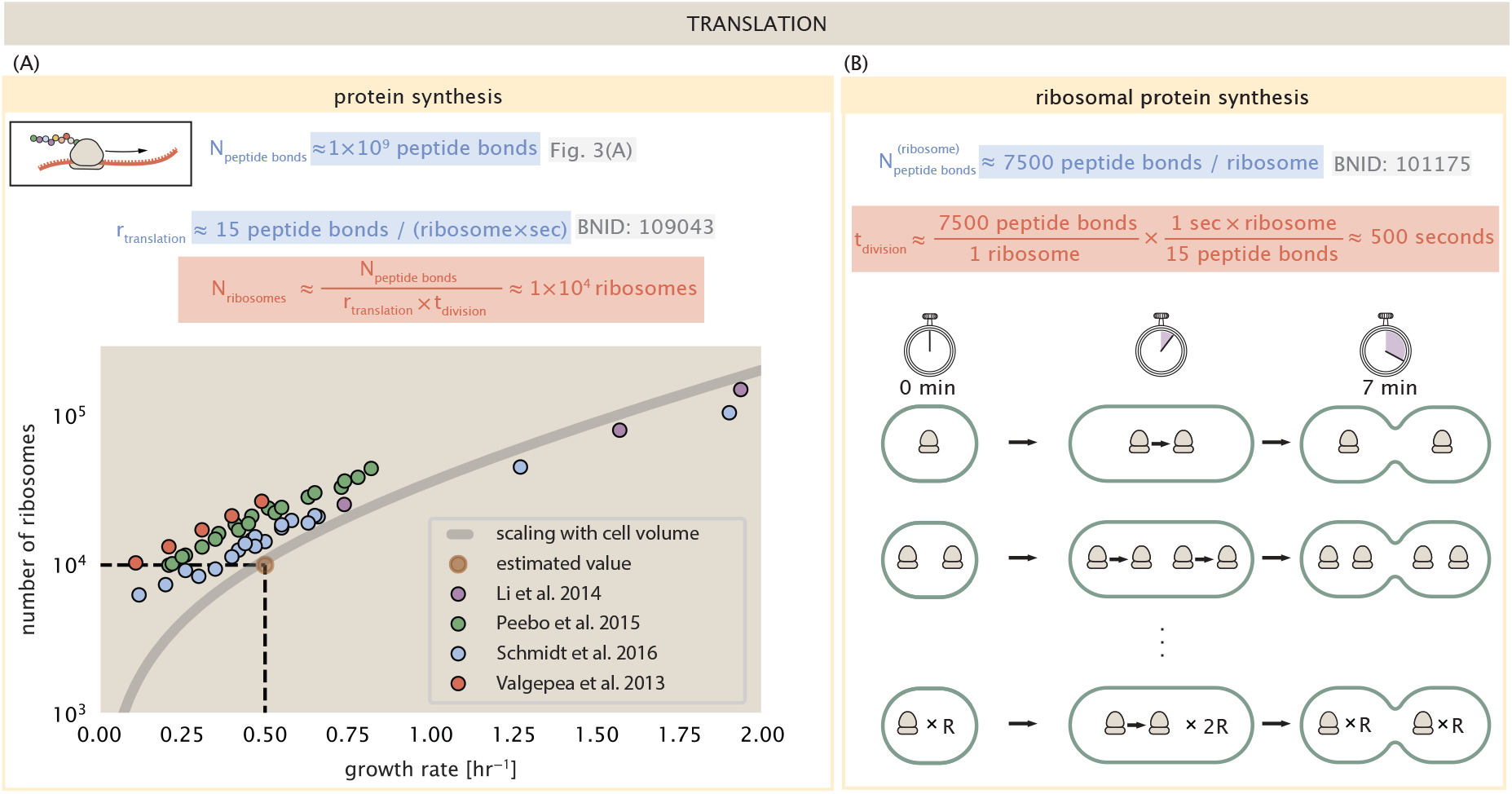
Estimation of the required number of ribosomes and the speed limit for bacterial replication. (A) Estimation of the number of ribosomes required to synthesize 10^9^ peptide bonds with an elongation rate of 15 peptide bonds per second. The average abundance of ribosomes is plotted as a function of growth rate. Our estimated values are shown for a growth rate of 0.5 hr^−1^. Grey lines correspond to the estimated complex abundance calculated at different growth rates. (B) Estimation for the time to replicate a ribosome. This rate is independent of the number of ribosomes R and instead is limited by the time required to double an individual ribosome. **Figure 9-Figure supplement 1. Estimate and observed abundance and growth rate dependence of tRNA ligases.**

Having completed our circuit through key processes of cellular growth outlined in ***Figure 1***, we can now take stock of our understanding of the observed growth rate dependence and abundances of various protein complexes. We note that, broadly speaking, these simple estimates have been reasonably successful in quantitatively describing the observations in the proteomic data, suggesting that the proteome is tuned in composition and absolute abundance to match the growth rate requirements without any one process representing a singular bottleneck or rate limiting step in division. However, in our effort to identify key limitations on growth, there are two notable observations that we wish to emphasize.

The first is a recurring theme throughout the estimates investigated here, which is that any inherent biochemical rate limitation can be overcome by expressing more proteins. We can view this as a parallelization of each biosynthesis task, which helps explain why bacteria tend to increase their protein content (and cell size) as growth rate increases (***Ojkic et al., 2019***). The second, and ultimately the most significant in defining the cellular growth rate, is that the synthesis of ribosomal proteins presents a special case where parallelization is *not* possible and thereby imposes a limit on the fastest possible growth rate. Each ribosome has ≈ 7500 amino acids across all of its protein components which must be strung together as peptide bonds through the action of another ribosome. Once again using a modest elongation rate of ≈ 15 amino acids per second, we arrive at an estimate of ≈ 500 seconds or ≈ 7 minutes to replicate a single ribosome. This limit, as remarked upon by others (***Dill et al., 2011***), serves as a hard theoretical boundary for how quickly a bacterium like *E. coli* can replicate. As each ribosome would therefore need to copy itself, this 7 minute speed limit is independent of the number of ribosomes per cell [***Figure 9***(B)], yet assumes that the only proteins that need to be replicated for division to occur are ribosomal proteins, a regime not met in biological reality. This poses an optimization problem for the cell – how are the translational demands of the entire proteome met without investing resources in the production of an excess of ribosomes?

This question, more frequently presented as a question of optimal resource allocation, has been the target of an extensive dialogue between experiment and theory over the past decade. In a now seminal work, ***Scott et al.*** (***2010***) present an elegant treatment of resource allocation through partitioning of the proteome into sectors – one of which being ribosome-associated proteins whose relative size ultimately defines the total cellular growth rate. In more recent years, this view has been more thoroughly dissected experimentally (***Klumpp and Hwa, 2014***; ***Basan et al., 2015***; ***Dai et al., 2018***, ***2016***; ***Erickson et al., 2017***) and together have led to a paradigm-shift in how we think of cellular physiology at the proteomic-level. However, the quantitative description of these observations is often couched in terms of phenomenological constants and effective parameters with the key observable features of expression often computed in relative, rather than absolute, abundances. Furthermore, these approaches often exclude or integrate away effects of cell size and chromosome content, which we have found through our estimates to have important connections to the observed cellular growth rate and proteomic content.

In the closing sections of this work, we explore how ribosomal content, total protein abundance, and chromosomal replication are intertwined in their control over the cellular growth rate. To do so, we take a more careful view of ribosome abundance, increasing the sophistication of our analysis by exchanging our order-of-magnitude estimates for a minimal mathematical model of growth rate control. This is defined by parameters with tangible connections to the biological processes underlying cellular growth and protein synthesis. Using this model, we interrogate how the size of the ribosome pool and its corresponding translational capacity enable cells to maintain a balance between the supply of amino acids via metabolism and catabolism and their consumption through the peptide bond formation required for growth.

### Maximum Growth Rate is Determined by the Ribosomal Mass Fraction

The 7 minute speed limit shown in ***Figure 9***(B) assumes all proteins in the cell are ribosomes. In order to connect this to the experimental data (and physiological reality more broadly), we first need to relax this assumption and determine a translation-limited growth rate. Here, we will assume that the cell is composed of *N*_pep_ peptide bonds and R ribosomes, whose precise values will depend on the growth rate *λ*. The protein subunits of each ribosomal protein sum to a total of ≈ 7500 amino acids as noted earlier, which we denote by *L_R_*. With an average mass of an amino acid of *m*_*AA*_ ≈ 110 Da (BNID: 104877), the total ribosomal mass fraction Φ_*R*_ is given by

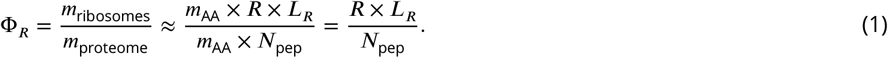

For exponentially growing cells (***Godin et al., 2010***), the rate of cellular growth will be related to the rate of protein synthesis via

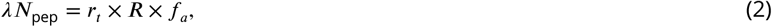

where *r*_*t*_ is the translation rate. Here, we’ve introduced a multiplicative factor *f*_*a*_ which represents the fraction of the ribosomes that are actively translating. This term allows us to account for immature or non-functional ribosomes or active sequestration of ribosomes through the action of the secondary messenger alarmone (p)ppGpp in poorer nutrient conditions (***Hauryliuk et al., 2015***).

Combining ***Equation 1*** and ***Equation 2*** results in an expression for a translation-limited growth rate, which is given by

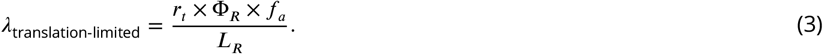

This result, derived in a similar manner by others (***Dennis et al., 2004***; ***Klumpp et al., 2013***), reflects mass-balance under steady state growth and has long provided a rationalization of the apparent linear increase in *E. coli*’s ribosomal content as a function of growth rate (***Maalœ, 1979***; ***Dennis et al., 2004***; ***Scott et al., 2010***; ***Dai et al., 2016***). The left-hand panel of ***Figure 10***(A) shows this growth rate plotted as a function of the ribosomal mass fraction. In the regime where all ribosomes are active (*f_a_* = 1) and the entire proteome is composed of ribosomal proteins (Φ_*R*_ = 1), indeed, we arrive at the maximum theoretical growth rate of *r_t_/L_R_*, and ≈ 7 min for *E. coli*.

**Figure 10.**
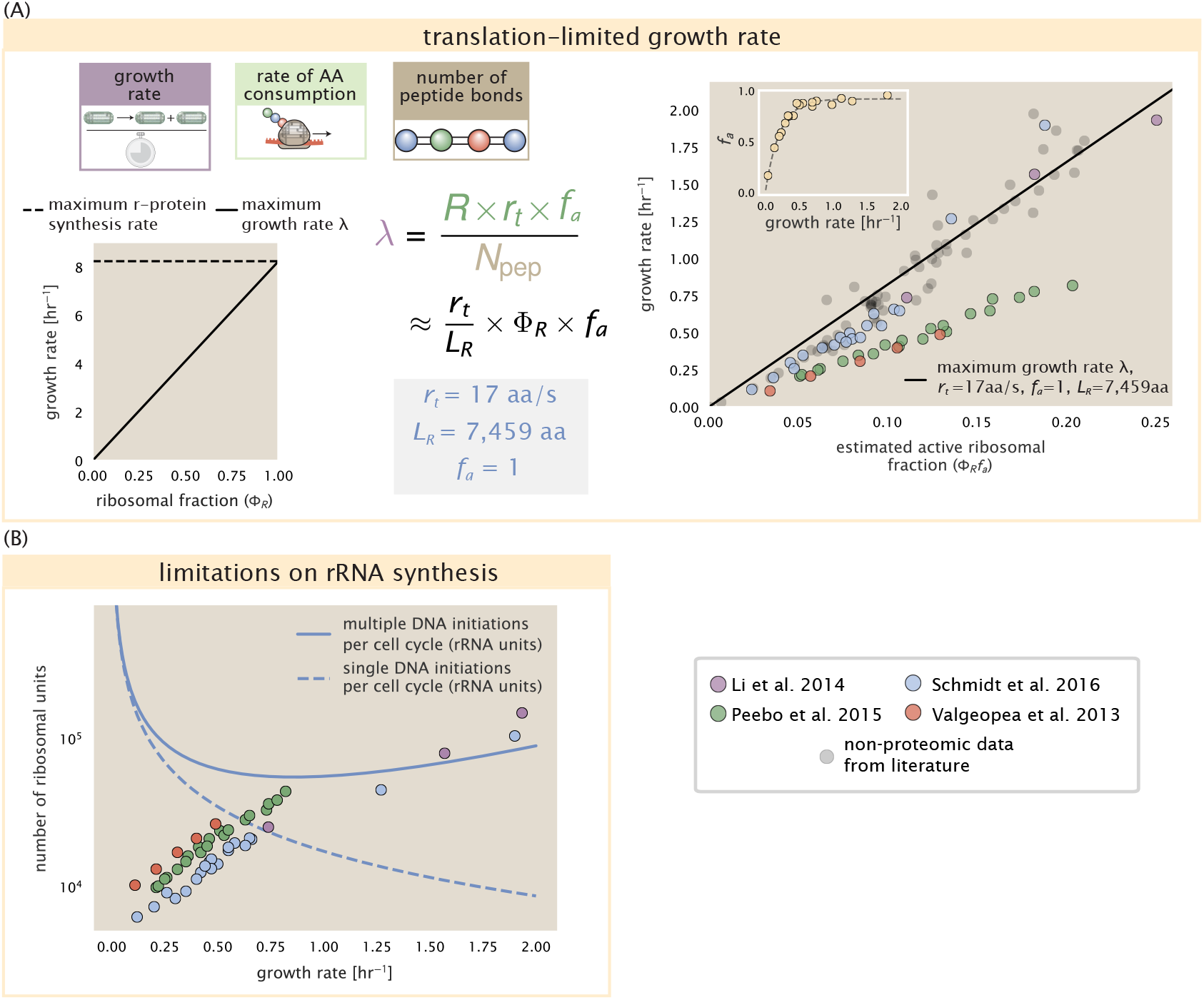
Translation-limited growth rate. (A) *left*: Translation-limited growth as a function of the ribosomal fraction. The solid line is calculated for an elongation rate of 17 aa per second. The dashed line corresponds to the maximum rate of ribosomal protein synthesis (≈ 7 min). *right*: Translation-limited growth rate as a function of the actively translating ribosomal fraction. The actively translating ribosomal fraction is calculated using the estimated values of *f_a_* from ***Dai et al.*** (***2016***) (shown in inset; see Section “Calculation of active ribosomal fraction" for additional detail). Gray data points show additional measurements from literature and considered further in the supplemental figure part (A). (B) Maximum number of rRNA units that can be synthesized as a function of growth rate. Solid curve corresponding to the rRNA copy number is calculated by multipyling the number of rRNA operons by the estimated number of 〈# ori〉 at each growth rate. The quantity was calculated using Equation 4 and the measurements from ***Si et al.*** (***2017***). The dashed line shows the maximal number of functional rRNA units produced from a single chromosomal initiation per cell cycle. Figure 10-Figure supplement 1. Comparison of Φ_*R*_*f*_*a*_ with literature and estimation of 〈# ori〉.

Connecting ***Equation 3*** to the proteomic data serving as the centerpiece of our work, however, requires knowledge of *f*_*a*_ at each growth rate as proteomic measurements only provide a measure of Φ_*R*_. Recently, ***Dai et al.*** (***2016***) determined *f*_*a*_ as a function of the growth rate (***Figure 10***(A), right-hand panel, inset), revealing that *f*_*a*_ ≈ 1 at growth rates above 0.75 hr^−1^ and *f*_*a*_ < 1 at slower growth rates. Using these data, we inferred the approximate active fraction (see the Appendix Section “Calculation of active ribosomal fraction”) at each growth rate and used this to compute Φ_*R*_ × *f*_*a*_ (***Figure 10***(A), colored points in right-hand panel). Importantly, these data largely skirt the translation-limited growth rate determined using ***Equation 3***, where we have taken *r*_*t*_ to be the maximal elongation rate of 17 amino acids per second measured by ***Dai et al.*** (***2016***). There is a notable discrepancy between the data collected in ***Schmidt et al.*** (***2016***); ***Li et al.*** (***2014***) and that collected from ***Valgepea et al.*** (***2013***); ***Peebo et al.*** (***2015***). When compared to other measurements (non-proteomic based) of the active ribosome mass fraction based on measurements of total RNA to total protein mass ratios (***Figure 10***(B), grey points in right-hand panel and further detailed in ***Figure 10-Figure Supplement 1***), the data from ***Valgepea et al.*** (***2013***) and ***Peebo et al.*** (***2015***) are notably different, suggesting there may be a systematic bias in these two sets of measurements.

Together, these results illustrate that the growth rates observed across the amalgamated data sets are indeed close to the translation-limited growth rate set by ribosomal activity, at least for the data reported in ***Schmidt et al.*** (***2016***) and ***Li et al.*** (***2014***). While this is a useful framework to consider how the relative abundance of ribosomes (compared to all other proteins) defines the growth rate, it is worth noting that as growth rate increases, so does the cell size and therefore so will the total proteomic mass (***Basan et al., 2015***). With a handle on how elongation rate and the total number of peptide bonds per proteome is related to the growth rate, we now expand this description to account for the increasing chromosomal content, cell size, and ribosome copy number at faster growth rates, enabling us to identify a potential bottleneck in the synthesis of rRNA.

### rRNA Synthesis Presents a Potential Bottleneck During Rapid Growth

Even under idealized experimental conditions, *E. coli* rarely exhibits growth rates above 2 hr^−1^ (***Bremer and Dennis, 2008***), which is still well-below the synthesis rate of a single ribosome, and below the maximum growth rates reported for several other bacteria (***Roller et al., 2016***). While we have considered potential limits imposed by translation of ribosomal *proteins*, here we consider potential limiting regimes specific to the synthesis of rRNA.

Due to multiple initiations of chromosomal replication per cell doubling, the effective number of rRNA operons increases with growth rate and will do so in proportion to the average number of chromosomal origins per cell, 〈# ori〉. This later parameter is set by how often replication must be initiated in order to keep up with the cell doubling time τ, whose time may be shorter than the cell cycle time τ_*cyc*_ (referring to the time from replication initiation to cell division) (***Dennis et al., 2004***; ***Ho and Amir, 2015***). This is quantified by

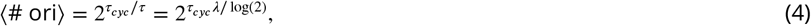

where the doubling time τ is related to the growth rate by τ = log(2)/*λ*. As the rRNA operons are predominantly located close to the origin of replication (BNID: 100352), we make the simplifying assumption that that the number of rRNA operons will be directly proportional to 〈# ori〉. We used the experimental measurements of τ (the timescale of chromosome replication and cell division) and (the timescale of a cell doubling) [***Figure 10 – Figure Supplement1***(B)] to calculate 〈# ori〉 with ***Equation 4*** as a function of growth rates. For growth rates above about 0.5 hr^−1^ *t*_*cyc*_ is approximately constant at about 70 minutes, implying an exponential increase in 〈# ori〉 and the rRNA operon copy number for growth rates above 0.5 hr^−1^.

Returning to our rule-of-thumb that one functional rRNA unit is produced per second per transcribing operon, we can estimate the maximum number of ribosomes that could be made as a function of growth rate (***Figure 10***(B), blue curve). Although we expect this estimate to significantly overestimate rRNA abundance at slower growth rates (*λ* < 0.5 hr^−1^), this provides a useful reference alongside the proteomic measurements, particularly in the regime of fast growth. For growth rates above about 1 hr^−1^ in particular, we find that cells will need to transcribe rRNA near their maximal rate. As a counter example, if *E. coli* did not initiate multiple rounds of replication, but could still replicate their chromosome within the requisite time limit, they would be unable to make enough rRNA for the observed number of ribosomes (dashed blue curve in ***Figure 10***(C)). The convergence between the maximum rRNA production and measured ribosome copy number suggests rRNA synthesis may begin to present a bottleneck at the fastest growth rates in *E. coli* due to the still-limited copies of rRNA genes.

### Rapid Growth Requires *E. coli* to Increase Both Cell Size and Ribosomal Mass Fraction

In the right-hand side of ***Figure 10***(B) we also find that above about 0.75 hr^−1^ the growth rate is determined solely by the ribosomal mass fraction Φ_*R*_, since *f*_*a*_ is close to 1, and *r*_*t*_ is near its maximal rate (***Dai et al., 2016***). While Φ_*R*_ will need to increase in order for cells to grow faster, the fractional dependence in ***Equation 3*** gives little insight into how this scaling is actually achieved by the cell.

It is now well-documented that *E. coli* cells add a constant volume per origin of replication, which is robust to a remarkable array of cellular perturbations (***Si et al., 2017***). Given the proteomic measurements featured in this work, we find that the ribosome copy number also scales in proportion to 〈# ori〉 (Figure 11(A)). However, an increase in ribosome abundance alone is not necessarily sufficient to increase growth rate and we also need to consider how Φ_*R*_ varies with 〈# ori〉. As shown in Figure 11(B), we find that the deviations in protein expression with 〈# ori〉 are largely restricted to regions of ribosomal protein genes. Here we have calculated the position-dependent protein expression across the chromosome by a running Gaussian average of protein copy number (20 kbp st. dev. averaging window) based on each gene’s transcriptional start site. These were median-subtracted to account for the change in total protein abundance with 〈# ori〉. This result suggests that Φ_*R*_ is also being tuned in proportion to 〈# ori〉 under nutrient-limited growth. Importantly, it is through this additional dependence on Φ_*R*_, combined with the exponential increase in 〈# ori〉 that was noted in the previous section, that E. coli exhibits an exponential increase in cell size with growth rate.

**Figure 11.**
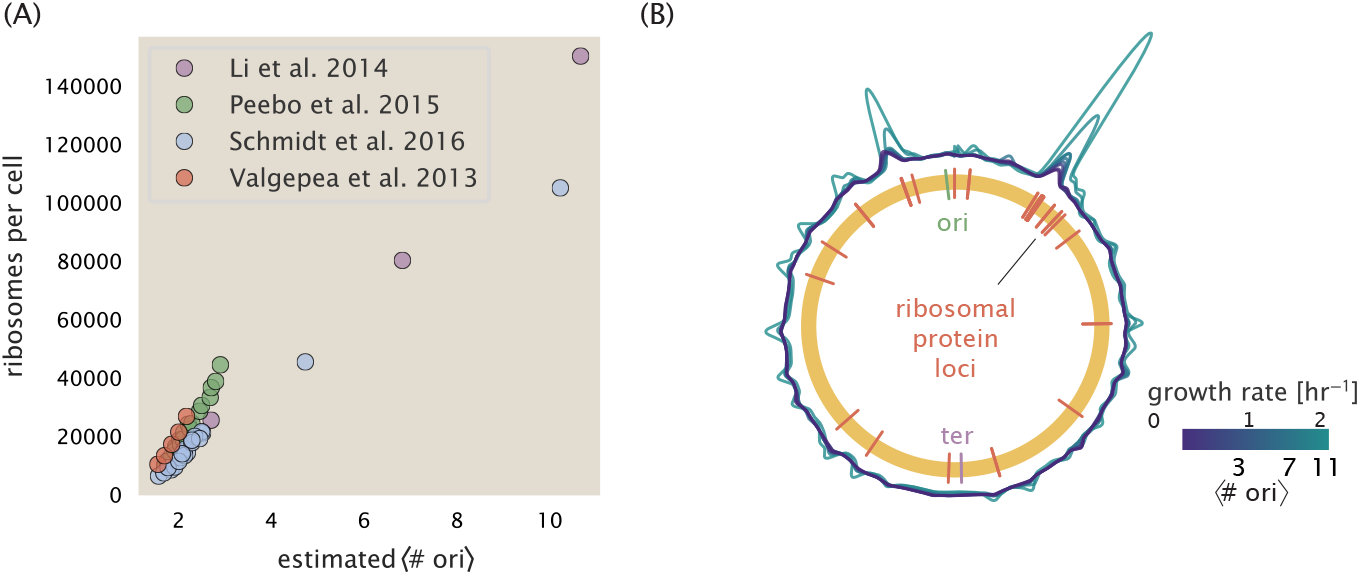
Cells increase both absolute ribosome abundance and *Φ_R_* with 〈# ori〉. (A) Plot of the ribosome copy number estimated from the proteomic data against the estimated 〈# ori〉 (see Appendix Section “Estimation of 〈# ori〉/ 〈# ori〉 and 〈# ori〉 for additional details). (B) A running Gaussian average (20 kbp st. dev.) of protein copy number is calculated for each growth condition considered by (Schmidt et al., 2016) based on each gene’s transcriptional start site. Since total protein abundance increases with growth rate, protein copy numbers are median-subtracted to allow comparison between growth conditions. 〈# ori〉 are estimated using the data in (A) and Equation 4.

## A Minimal Model of Nutrient-Mediated Growth Rate Control

While the preceding subsections highlight a dominant role for ribosomes in setting the growth rate, our analysis on the whole has emphasized how the total proteomic content changes in response to variable growth conditions and growth rate. In this final section we employ a minimal model of growth rate control to better understand how this interconnection between ribosomal abundance and total protein abundance influences the observed growth rate.

Here we propose that cells modulate their protein abundance in direct response to the availability of nutrients in their environment. As noted earlier, bacteria can modulate ribosomal activity through the secondary messenger molecules like (p)ppGpp in poorer nutrient conditions [***Figure 10***(C, inset); ***Dai et al.*** (***2016***)]. Importantly, these secondary messengers also cause global changes in transcriptional and translational activity (***Hauryliuk et al., 2015***; ***Zhu and Dai, 2019***; ***Büke et al., 2020***). In *E. coli*, amino acid starvation leads to the accumulation of de-acylated tRNAs at the ribosome’s A-site and a strong increase in (p)ppGpp synthesis activity by the enzyme RelA (***Hauryliuk et al., 2015***). Along with this, there is increasing evidence that (p)ppGpp also acts to inhibit the initiation of DNA replication (***Kraemer et al., 2019***), providing a potential mechanism for cells to lower 〈# ori〉 and maintain a smaller cell size in poorer nutrient conditions (***Fernández-Coll et al., 2020***).

To consider this quantitatively, we assume that cells modulate their proteome (total number of peptide bonds *N*_pep_, number of ribosomes ***R***, and ribosomal fraction Φ_*R*_) to better maximize their rate of peptide elongation *r*_*t*_. The elongation rate *r*_*t*_ will depend on how quickly the ribosomes can match codons with an amino-acyl tRNA, along with the subsequent steps of peptide bond formation and translocation. This ultimately depends on the cellular concentration of amino acids, which we treat as a single effective species, [*AA*]_eff_. In our model, we determine the the rate of peptide elongation *r*_*t*_ and achievable growth rate as simply depending on the supply of amino acids (and, therefore, also amino-acyl tRNAs), through a parameter *r_AA_* in units of AA per second, and the rate of amino acid consumption by protein synthesis (*r*_*t*_ × *R* × *f*_*a*_). This is shown schematically in ***Figure 12***(A) and derived in the Appendix Section “Derivation of Minimal Model for Nutrient-Mediated Growth Rate Control”. Given our observation that general protein synthesis and energy production are not limiting, we assume that other molecular players required by ribosomes such as elongation factors and GTP are available in sufficient abundance.

**Figure 12.**
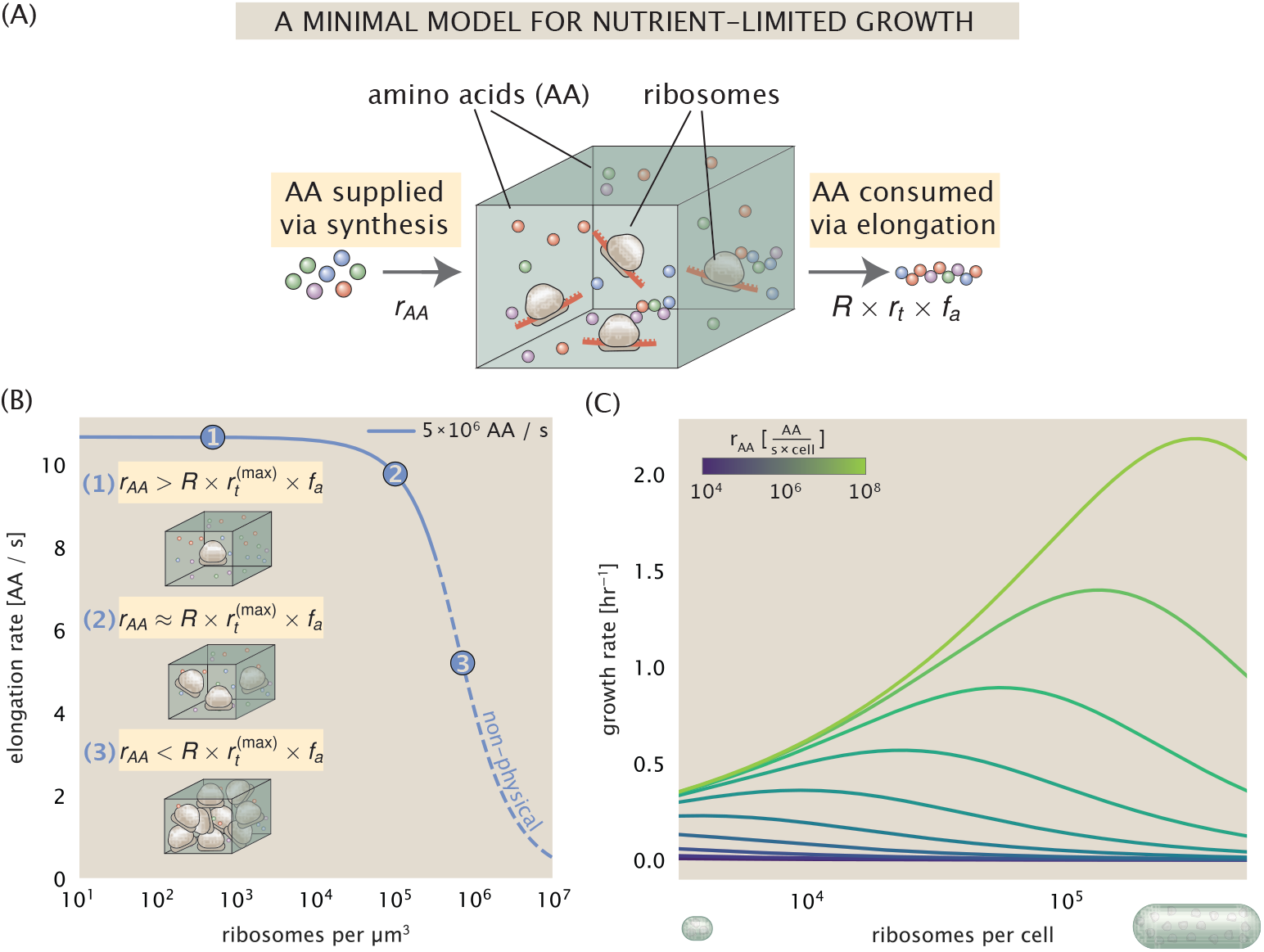
A minimal model of growth rate control under nutrient limitation. (A) We consider a unit volume of cellular material composed of amino acids (colored spheres) provided at a supply rate *r_AA_*. These amino acids are polymerized by a pool of ribosomes (brown blobs) at a rate *r*_*t*_ × *R* × *f*_*a*_, where *r*_*t*_ is the elongation rate, R is the ribosome copy number in the unit volume, and *f*_*a*_ is the fraction of those ribosomes actively translating. (B) The observed elongation rate is plotted as a function of the number of ribosomes. The three points correspond to three regimes of ribosome copy numbers and are shown schematically on the left-hand side. The region of the curve shown as dashed lines represents a non-physical copy number, but is shown for illustrative purposes. This curve was generated using an amino acid supply rate of 5 × 10^6^ AA / s, a maximal elongation rate of 17.1 AA / s, *f*_*a*_ = 1, and a unit cell volume of *V* = 1 fL. See Appendix Section “Derivation of Minimal Model for Nutrient-Mediated Growth Rate Control” for additional model details. (C) The cellular growth rate is plotted as a function of total cellular ribosome copy number for different cellular amino acid supply rates, with blue and green curves corresponding to low and high supply rates, respectively. As the ribosome copy number is increased, so too is the cell size and total protein abundance *N*_pep_. We direct the reader to the Supplemental Information for discussion on the inference of the relationship between cell size, number of peptide bonds, and ribosome copy number. **Figure 12-Figure supplement 1.** An interactive figure for exploration of the model parameter space.

In ***Figure 12***(B), we illustrate how the elongation rate will depend on the ribosomal copy number. Here, we have considered an arbitrarily chosen *r*_*AA*_ = 5 × 10^6^ AA s^−1^ μm^−3^ and *f*_*a*_ = 1 for a unit cell volume V = 1fL (we provide the interactive figure ***Figure 12-Figure Supplement 1*** which allows the user to explore different regimes of this parameter space). At low ribosome copy numbers, the observed elongation rate is dependent primarily on *[AA]_eff_* through *r_AA_* [as 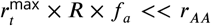, point (1) in ***Figure 12***(B)]. As the ribosome copy number is increased such that the amino acid supply rate and consumption rate are nearly equal [point (2) in ***Figure 12***(B)], the observed elongation rate begins to decrease sharply. When the ribosome copy number is increased even further, consumption at the maximum elongation rate exceeds the supply rate, yielding a significantly reduced elongation rate [point (3) in ***Figure 12***B)]. While the elongation rate will always be dominated by the amino acid supply rate at sufficiently low ribosome copy numbers, the elongation rate at larger ribosome abundances can be increased by tuning *f*_*a*_ such that not all ribosomes are elongating, reducing their total consumption rate.

### Optimal Ribosomal Content and Cell Size Depend on Nutrient Availability and Metabolic Capacity

To relate elongation rate to growth rate, we constrain the set of parameters based on our available proteomic measurements; namely, we restrict the values of *R*, *N*_pep_, and cell size to those associated with the amalgamated proteomic data (described in the Appendix Section “Estimation of Total Protein Content per Cell”). We then consider how changes in the nutrient conditions, through the parameter *r_AA′_*, influence the maximum growth rate as determined by ***Equation 3***. ***Figure 12***(C) shows how the growth rate depends on the rate of amino acid supply *r_AA_* as a function of the cellular ribosome copy number. A feature immediately apparent is the presence of a maximal growth rate increases with increasing *r_AA_*. Importantly, however, there is an optimum set of *R, N_pep_*, and cell size that are strictly dependent on the value of *r_AA_*. This shows that increasing the ribosomal concentration beyond the cell’s metabolic capacity will have the adverse consequence of depleting the supply of amino acids and lead to a concomitant decrease in the elongation rate *r*_*t*_ [***Figure 12***(B)] and growth rate. This helps us understand that while it is important for cells to increase their ribosomal content and cell size in order to increase growth rate, cells will better maximize their achievable growth rate by tuning these parameters according to the available nutrient conditions, since this is ultimately what allows cells to reach the peak for each curve shown in ***Figure 12***(C).

Also of note is the growth rate trends observed at low values of *r_AA_* [purple and blue lines in ***Figure 12***(C)], representative of growth in nutrient-poor media. In these conditions, there no longer exists a peak in growth, at least within the range of physiologically-relevant ribosome copy numbers considered here. This is a regime, associated with slower growth rates, where cells limit their pool of actively translating ribosomes by decreasing *f*_*a*_ (***Figure 10***(A), right-hand panel, inset; (***Dai et al., 2016***)), likely due to having excess ribosomes relative to the cell’s metabolic capacity. By reducing the fraction of actively translating ribosomes, we find that cells instead prioritize maintaining their pool of available amino acids [*AA*]_eff_ and increasing the achievable translation elongation rate.

## Discussion

Continued experimental and technological improvements have led to a treasure trove of quantitative biological data (*Hui et al., 2015*; *Schmidt et al., 2016*; *Si et al., 2017*; *Gallagher et al., 2020*; *Peebo et al., 2015*; *Valgepea et al., 2013*), and an ever advancing molecular view and mechanistic understanding of the constituents that support bacterial growth (*Taheri-Araghi et al., 2015*; *Morgenstein et al., 2015*; *Si et al., 2019*; *Karr et al., 2012*; *Kostinski and Reuveni, 2020*; *Macklin et al., 2020*). In this work we have compiled and curated what we believe to be the state-of-the-art knowledge on proteomic copy number across a broad range of growth conditions in *E. coli*. Beyond compilation, we have taken a detailed approach in ensuring that the absolute protein abundances reported are directly comparable across growth rates *and* data sets, allowing us to make assertions about the physiology of *E. coli* rather than chalking up discrepancies with our simple estimates to experimental noise and systematic errors. We have made this data accessible through a GitHub repository, and an interactive figure that allows exploration of specific protein and protein complex copy numbers.

Through a series of order-of-magnitude estimates that traverse key steps in the bacterial cell cycle, this proteomic data has been a resource to guide our understanding of two key questions: what biological processes limit the absolute speed limit of bacterial growth, and how do cells alter their molecular constituents as a function of changes in growth rate or nutrient availability? While not exhaustive, our series of estimates provide insight on the scales of macromolecular complex abundance across four classes of cellular processes – the transport of nutrients, the production of energy, the synthesis of the membrane and cell wall, and the numerous steps of the central dogma.

In general, the copy numbers of the complexes involved in these processes were in reasonable agreement with our order-of-magnitude estimates. Since many of these estimates represent soft lower-bound quantities, this suggests that cells do not express proteins grossly in excess of what is needed for a particular growth rate. Several exceptions, however, also highlight the dichotomy between a proteome that appears to “optimize” expression according to growth rate and one that must be able to quickly adapt to environments of different nutritional quality. Take, for example, the expression of carbon transporters. Shown in ***Figure 2***(B), we find that cells always express a similar number of glucose transporters irrespective of growth condition. At the same time, it is interesting to note that many of the alternative carbon transporters are still expressed in low but non-zero numbers (≈ 10-100 copies per cell) across growth conditions. This may relate to the regulatory configuration for many of these operons, which require the presence of a metabolite signal in order for alternative carbon utilization operons to be induced (***Monod, 1949***; ***Laxhuber et al., 2020***). Furthermore, upon induction, these transporters are expressed and present in abundances in close agreement with a simple estimate.

Of the processes illustrated in ***Figure 1***, we arrive at a ribosome-centric view of cellular growth rate control. This is in some sense unsurprising given the long-held observation that *E. coli* and many other organisms vary their ribosomal abundance as a function of growth conditions and growth rate (***Scott et al., 2010***; ***Metzl-Raz et al., 2017***). However, through our dialogue with the proteomic data, two additional key points emerge. The first relates to our question of what process sets the absolute speed limit of bacterial growth. While a cell can parallelize many of its processes simply by increasing the abundance of specific proteins or firing multiple rounds of DNA replication, this is not so for synthesis of ribosomes [***Figure 10***(A)]. The translation time for each ribosome [≈ 7 min, ***Dill et al.*** (***2011***)] places an inherent limit on the growth rate that can only be surpassed if the cell were to increase their polypeptide elongation rate, or if they could reduce the total protein and rRNA mass of the ribosome. The second point relates to the long-observed correlations between growth rate and cell size (***Schaechter et al., 1958***; ***Si et al., 2017***), and between growth rate and ribosomal mass fraction. While both trends have sparked tremendous curiosity and driven substantial amounts of research in their own regards, these relationships are themselves intertwined. In particular, it is the need for cells to increase their absolute number of ribosomes under conditions of rapid growth that require cells to also grow in size. Further experiments are needed to test the validity of this hypothesis. In particular, we believe that the change in growth rate in response to translation-inhibitory drugs (such as chloramphenicol) could be quantitatively predicted, given one had precision measurement of the relevant parameters, including the fraction of actively translating ribosomes *f*_*a*_ and changes in the metabolic capacity of the cell (i.e. the rate that amino acids can be made available) for a particular growth condition.

While the generation of new ribosomes plays a dominant role in growth rate control, there exist other physical limits to the function of cellular processes. One of the key motivations for considering energy production was the physical constraints on total volume and surface area as cells vary their size (***Harris and Theriot, 2018***; ***Ojkic et al., 2019***). As *E. coli* get larger at faster growth rates, an additional constraint begins to arise in energy production and nutrient uptake due to the relative decrease in total surface area, where ATP is predominantly produced (***Szenk et al., 2017***). Specifically, the cell interior requires an amount of energy that scales cubically with cell size, but the available surface area only grows quadratically [***Figure 5***(A)]. While this threshold does not appear to be met for *E. coli* cells growing at 2 hr^−1^ or less, it highlights an additional constraint on growth given the apparent need to increase cell size in order to grow faster. This limit is relevant even to eukaryotic organisms, whose mitochondria exhibit convoluted membrane structures that nevertheless remain bacteria-sized organelles (***Guo et al., 2018***). In the context of bacterial growth and energy production more generally, we have mainly limited our analysis to the aerobic growth conditions associated with the proteomic data and further consideration will be needed for anaerobic growth.

This work is by no means meant to be a complete dissection of bacterial growth rate control, and there are many aspects of the bacterial proteome and growth that we neglected to consider. For example, other recent work (***Liebermeister et al., 2014***; ***Hui et al., 2015***; ***Schmidt et al., 2016***) has explored how the proteome is structured and how that structure depends on growth rate. In the work of ***Hui et al.*** (***2015***), the authors coarse-grained the proteome into six discrete categories being related to either translation, catabolism, anabolism, and others related to signaling and core metabolism. The relative mass fraction of the proteome occupied by each sector could be modulated by external application of drugs or simply by changing the nutritional content of the medium. While we have explored how the quantities of individual complexes are related to cell growth, we acknowledge that higher-order interactions between groups of complexes or metabolic networks at a systems-level may reveal additional insights into how these growth-rate dependences are mechanistically achieved. Furthermore, while we anticipate the conclusions summarized here are applicable to a wide collection of bacteria with similar lifestyles as *E. coli*, other bacteria and archaea may have evolved other strategies that were not considered. Further experiments with the level of rigor now possible in *E. coli* will need to be performed in a variety of microbial organisms to learn more about how regulation of proteomic composition and growth rate control has evolved over the past 3.5 billion years.

## Methods

### Data Analysis and Availability

All proteomic measurements come from the experimental work of ***Schmidt et al.*** (***2016***); ***Peebo et al.*** (***2015***); ***Valgepea et al.*** (***2013***) (mass spectrometry) and ***Li et al.*** (***2014***) (ribosomal profiling). Data curation and analysis was done programmatically in Python, and compiled data and analysis files are accessible through a GitHub repository (DOI:10.5281/zenodo.4091457) associated with this paper as well as on the associated paper website. Additionally, we provide two interactive figures that allow for rapid exploration of the compiled data sets as well as exploration of the parameter space of the minimal model.

## Acknowledgements

We thank Matthias Heinemann, Alexander Schmidt, and Gene-Wei Li for additional input regarding their data. We also thank all members of the Phillips, Theriot, Kondev, Garcia labs, as well as Ron Milo and Terry Hwa for useful discussions. We thank Suzannah M. Beeler, Jonas Cremer, Avi Flamholz, Soichi Hirokawa, and Manuel Razo-Mejia for reading and providing comments on drafts of this manuscript. R.P. is supported by La Fondation Pierre-Gilles de Gennes, the Rosen Center at Caltech, and the NIH 1R35 GM118043 (MIRA). J.A.T. is supported by the Howard Hughes Medical Institute, and NIH Grant R37-AI036929. N.M.B is a HHMI Fellow of The Jane Coffin Childs Memorial Fund. H.G.G. is supported by the Burroughs Wellcome Fund Career Award at the Scientific Interface, the Sloan Research Foundation, the Human Frontiers Science Program, the Searle Scholars Program, the Shurl & Kay Curci Foundation, the Hellman Foundation, the NIH Director’s New Innovator Award (DP2 OD024541-01), and an NSF CAREER Award (1652236). D.S.F. is supported by an NSF award (PHY-1607606) and the NIH (NIH R01-AI13699201).

---

**Figure 2–Figure supplement 1.**
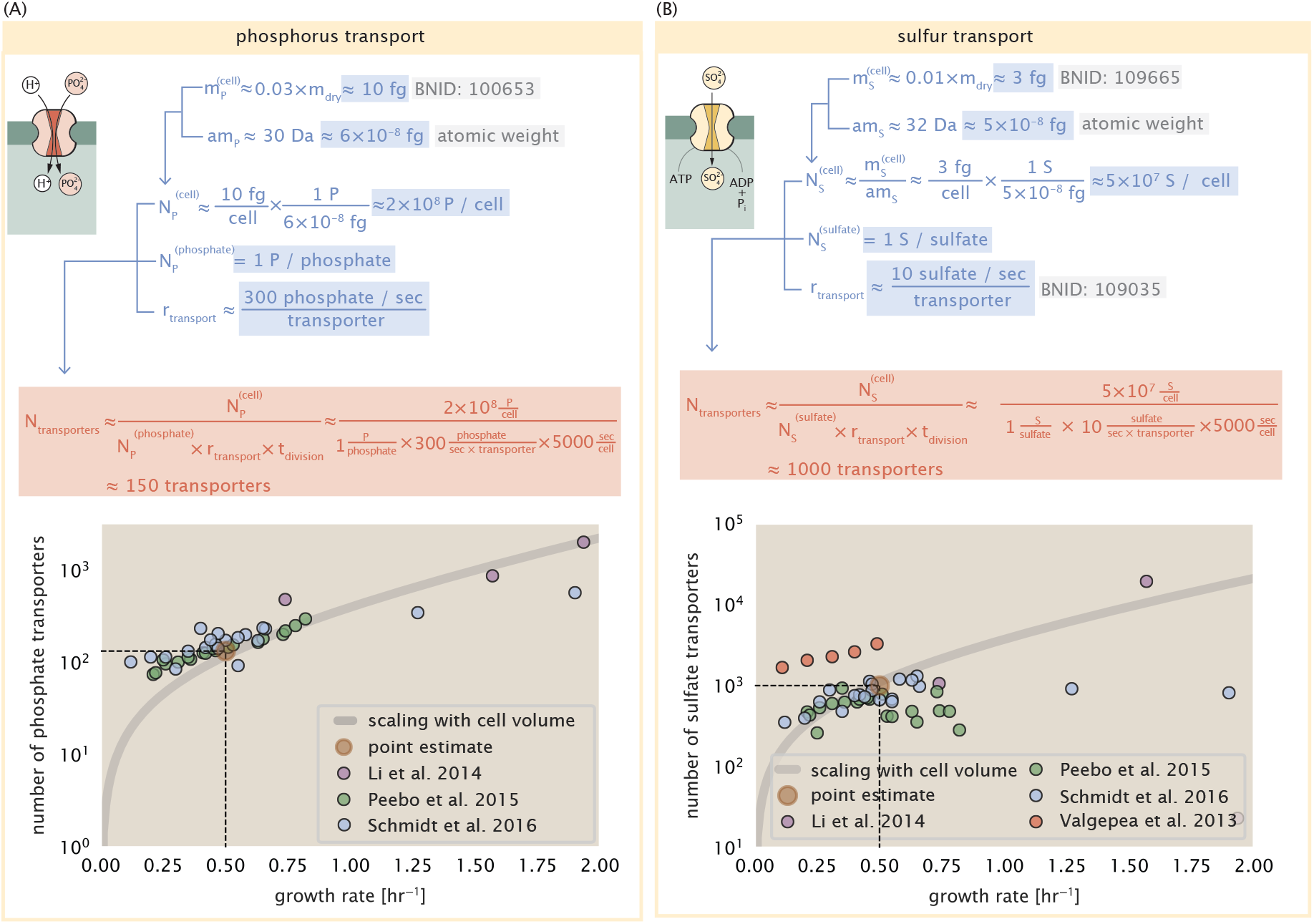
(A) Estimate for the number of PitA phosphate transport systems needed to maintain a 3% phosphorus *E. coli* dry mass. Points in plot correspond to the total number of PitA transporters per cell. (B) Estimate of the number of CysUWA complexes necessary to maintain a 1% sulfur *E. coli* dry mass. Points in plot correspond to average number of CysUWA transporter complexes that can be formed given the transporter stoichiometry [CysA]_2_[CysU][CysW][Sbp/CysP]. Grey line in (A) and represents the estimated number of transporters per cell at a continuum of growth rates.

**Figure 7–Figure supplement 1.**
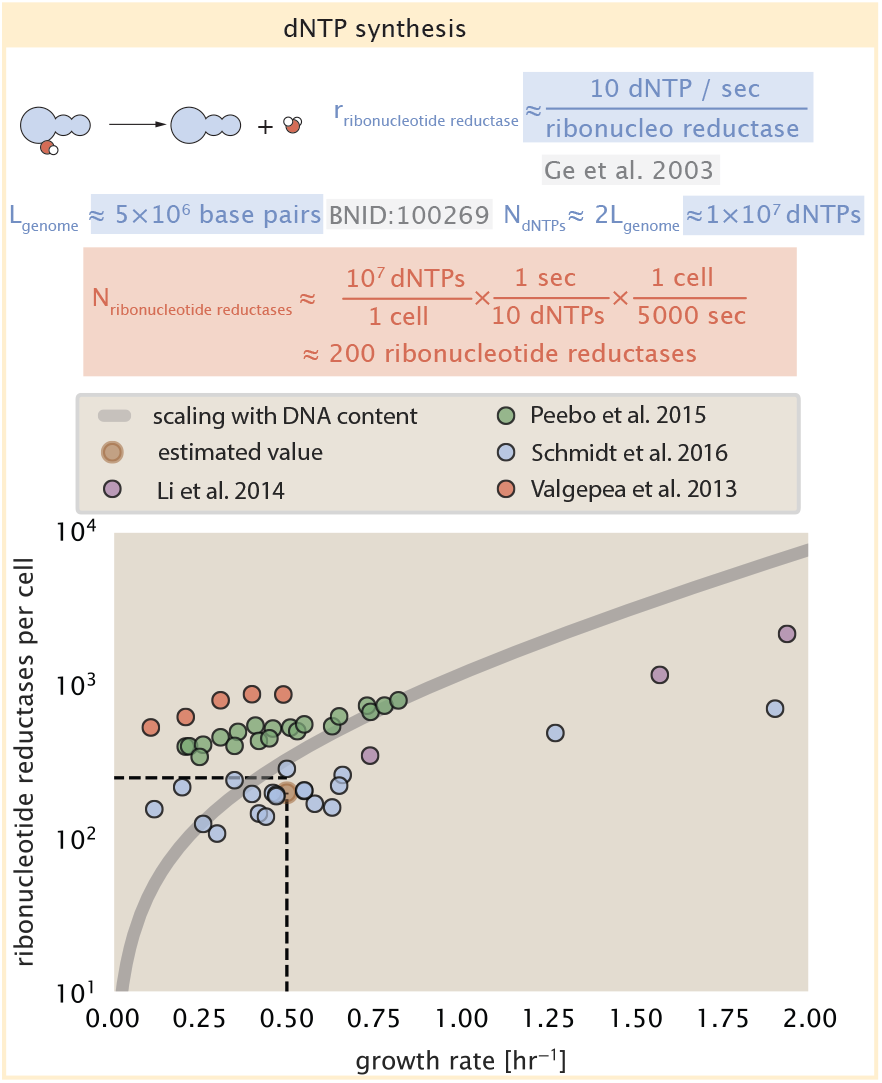
Estimate of the number of ribonucleotide reductase enzymes needed to facilitate the synthesis of ≈ 10^7^ dNTPs over the course of a 5000 second generation time. Points in the plot correspond to the total number of ribonucleotide reductase I ([NrdA]_2_[NrdB]_2_) and ribonucleotide reductase II ([NrdE]_2_[NrdF]_2_) complexes. Grey lines in top panel show the estimated number of complexes needed as a function of growth, the details of which are described in the Appendix.

**Figure 8–Figure supplement 1.**
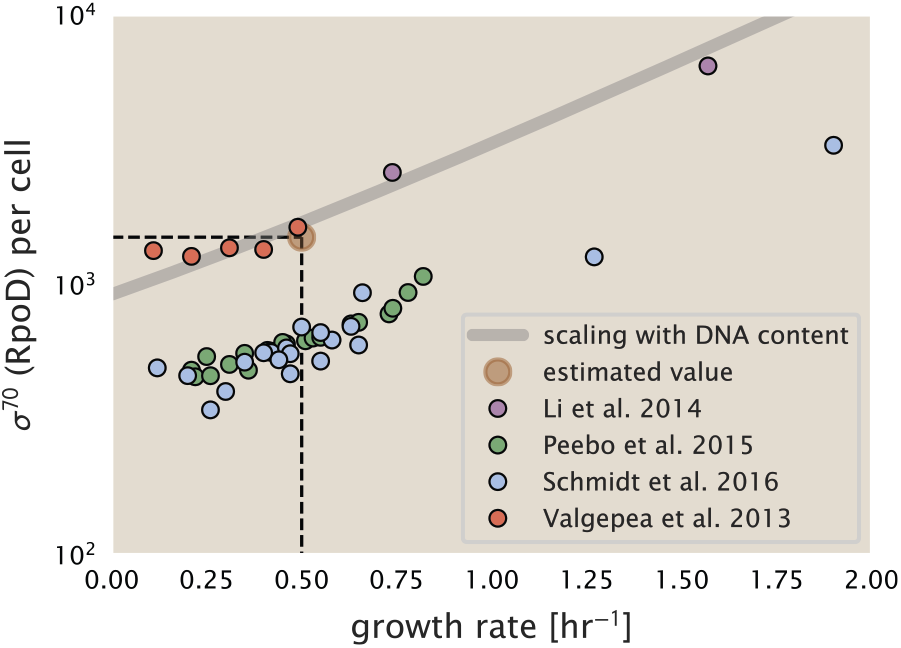
The abundance of σ^70^ as a function of growth rate. Estimated value for the number of RNAP is shown as a translucent brown point and grey line.

**Figure 9–Figure supplement 1.**
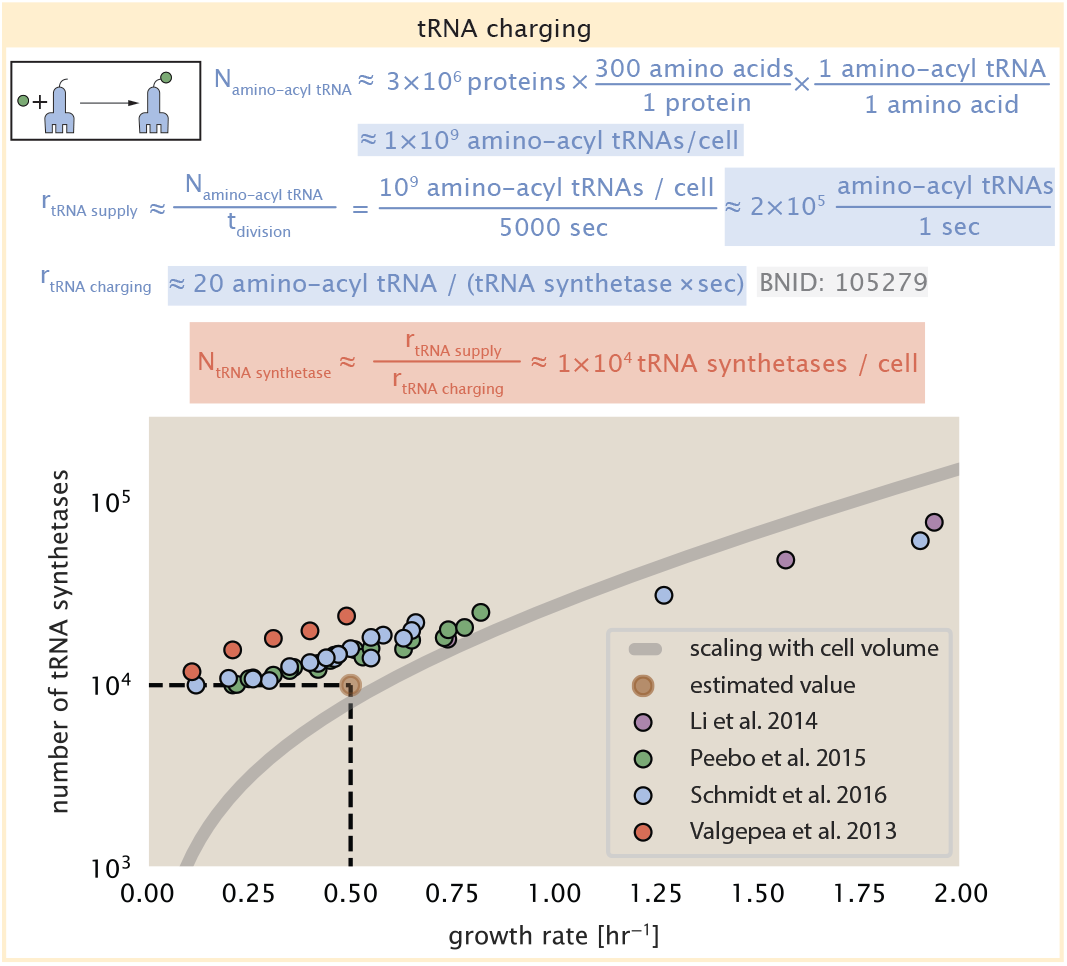
Estimation for the number of tRNA synthetases that will supply the required amino acid demand. The sum of all tRNA synthetases copy numbers are plotted as a function of growth rate ([ArgS], [CysS], [GlnS], [GltX], [IleS], [LeuS], [ValS], [AlaS]_2_, [AsnS]_2_, [AspS]_2_, [TyrS]_2_, [TrpS]_2_, [ThrS]_2_, [SerS]_2_, [ProS]_2_, [PheS]_2_[PheT]_2_, [MetG]_2_, [lysS]_2_, [HisS]_2_, [GlyS]_2_[GlyQ]_2_).

**Figure 10–Figure supplement 1.**
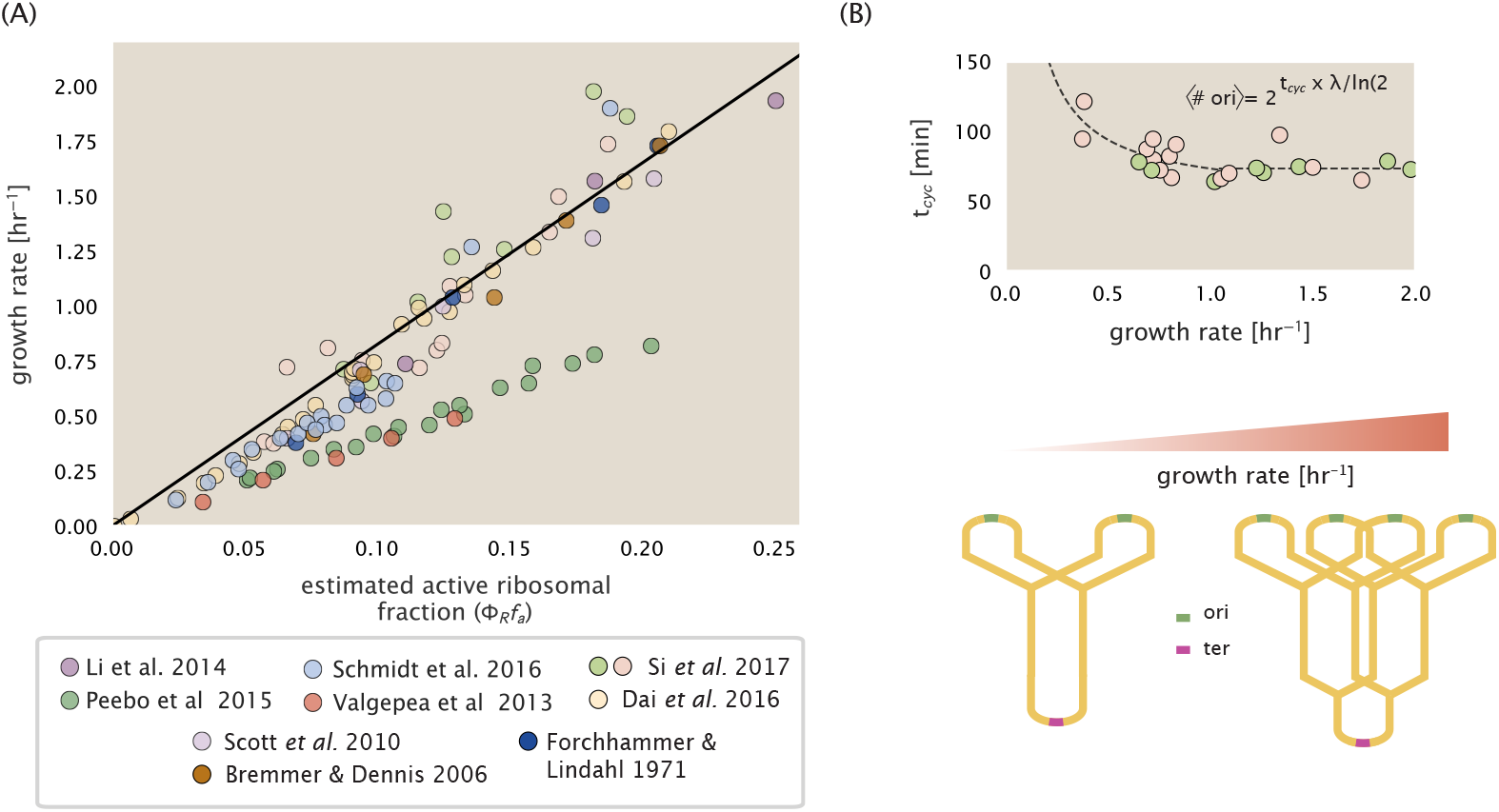
(A) Actively translating ribosomal fraction versus growth rate. The actively translating ribosomal fraction is calculated using the estimated values of *f*_*a*_ from ***Dai et al.*** (***2016***) (shown in inset; see the Appendix Section “Calculation of active ribosomal fraction for additional detail). Additional measurements in addition to the proteomic measurements are based on measurements of cellular RNA to protein ratio, with Φ_*R*_ ≈ the cellular RNA to protein ratio divided by 2.1 (***Dai et al., 2016***). (B) Experimental measurements of the cell doubling time τ and cell cycle time *t*_*cyc*_ from Si *et al.* (2017). Dashed line shows fit to the data, which were used to estimate 〈# ori〉. *t*_*cyc*_ was assumed to vary in proportion to τ for doubling times greater than 40 minutes, and reach a minimum value of 73 minutes. See Appendix Section “Estimation of 〈# ori〉/ 〈# ter〉 and 〈# ori〉 “ for additional details exact estimation of rRNA copy number. Red data points correspond to measurements in strain MG1655, while light green points are for strain NCM3722. Schematic shows the expected increase in replication forks (or number of ori regions) as *E. coli* cells grow faster.

**Figure 12–Figure supplement 1.**
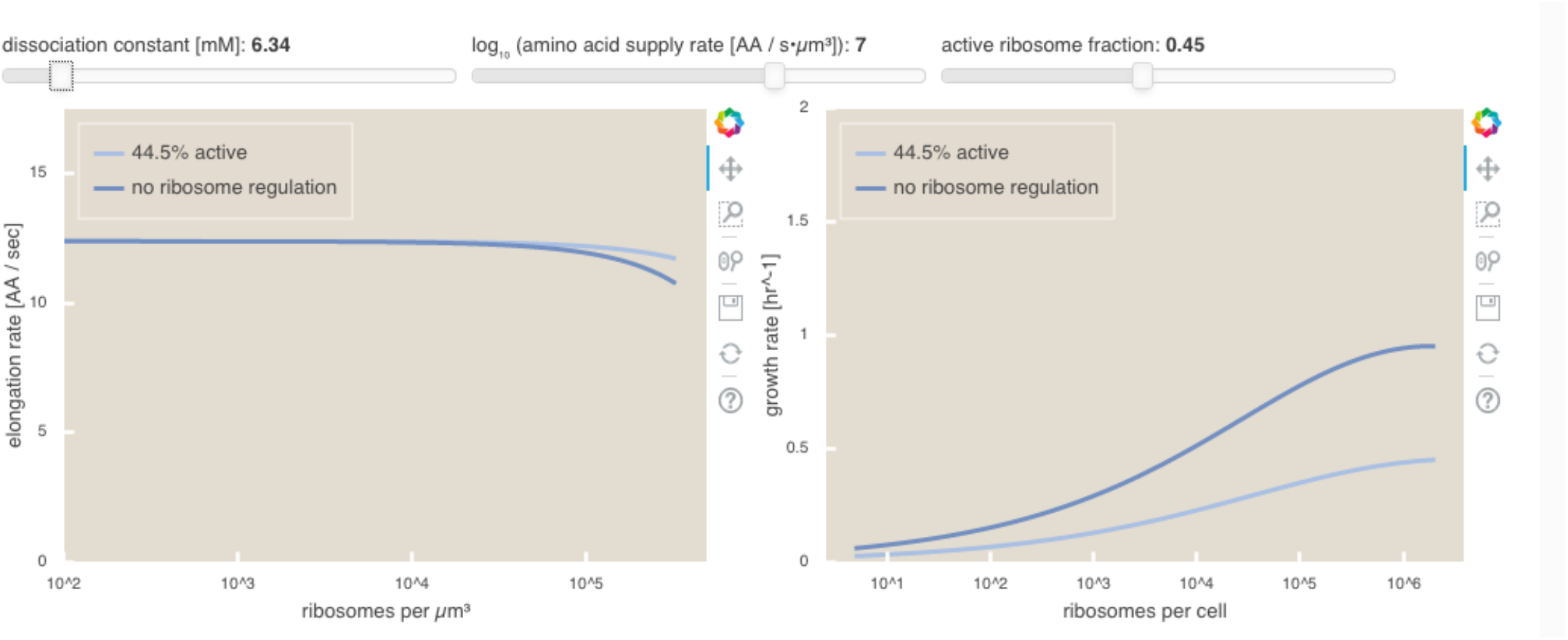
An interactive version of parts (B) and (C) of **Figure 12** which permit the user to modulate the rate of amino acid supply, the dissociation constant of amino acids to the ribosome, and the fraction of the ribosome pool that is actively translating. This interactive figure, and the code used to generate it, is available on the paper website.

## Appendix for: Fundamental limits on the rate of bacterial growth

## Additional Estimates of Fundamental Biological Processes

In the main text of this work, we present estimates for a significant number of fundamental biological processes that are necessary for cell division. While we believe the estimates provided in the main text provide a succinct summary of the corresponding process, we left out additional estimates of related processes for brevity. In this section of the appendix, we present these additional estimates in full.

## Nutrient Transport

In the main text, we make passing mention that in typical laboratory conditions, transport carbon often comes in the form of carbohydrates and sugar alcohols while other critical elements – such as nitrogen, sulfur, and phosphorus – are transported as inorganic ions. Below, we present estimates for the transport requirements of these materials.

## Nitrogen

We must first address which elemental sources must require active transport, meaning that the cell cannot acquire appreciable amounts simply via diffusion across the membrane. The permeability of the lipid membrane to a large number of solutes has been extensively characterized over the past century. Large, polar molecular species (such as various sugar molecules, sulfate, and phosphate) have low permeabilities while small, non-polar compounds (such as oxygen, carbon dioxide, and ammonia) can readily diffuse across the membrane. Ammonia, a primary source of nitrogen in typical laboratory conditions, has a permeability on par with water (≈ 1 × 10^5^ nm/s, BNID:110824). In nitrogen-poor conditions, *E. coli* expresses a transporter (AmtB) which appears to aid in nitrogen assimilation, though the mechanism and kinetic details of transport are still a matter of debate (***van Heeswijk et al., 2013***; ***Khademi et al., 2004***). Beyond ammonia, another plentiful source of nitrogen come in the form of glutamate, which has its own complex metabolism and scavenging pathways. However, nitrogen is plentiful in the growth conditions examined in this work, permitting us to neglect nitrogen transport as a potential rate limiting process in cell division in typical experimental conditions.

## Phosphorus

Phosphorus is critical to the cellular energy economy in the form of high-energy phosphodiester bonds making up DNA, RNA, and the NTP energy pool as well as playing a critical role in the post-translational modification of proteins and defining the polar-heads of lipids. In total, phosphorus makes up ≈3% of the cellular dry mass which in typical experimental conditions is in the form of inorganic phosphate. The cell membrane has remarkably low permeability to this highly-charged and critical molecule, therefore requiring the expression of active transport systems. In *E. coli*, the proton electrochemical gradient across the inner membrane is leveraged to transport inorganic phosphate into the cell (***Rosenberg et al., 1977***). Proton-solute symporters are widespread in *E. coli* (***Ramos and Kaback, 1977***; ***Booth et al., 1979***) and can have rapid transport rates of 50 to 100 molecules per second for sugars and other solutes (BNID: 103159; 111777). As a more extreme example, the proton transporters in the F_1_-F_0_ ATP synthase, which use the proton electrochemical gradient for rotational motion, can shuttle protons across the membrane at a rate of ≈ 1000 per second (BNID: 104890; 103390). In *E. coli* the PitA phosphate transport system has been shown to be very tightly coupled with the proton electrochemical gradient with a 1:1 proton:phosphate stoichiometric ratio (***Harris et al., 2001***; ***Feist et al., 2007***). Taking the geometric mean of the aforementioned estimates gives a plausible rate of phosphate transport on the order of 300 per second. Illustrated in ***Figure 2–Figure Supplement 1***(A), we can estimate that ≈ 200 phosphate transporters are necessary to maintain an ≈ 3% dry mass with a 5000 s division time. This estimate is consistent with observation when we examine the observed copy numbers of PitA in proteomic data sets (plot in ***Figure 2–Figure Supplement 1***(A)). While our estimate is very much in line with the observed numbers, we emphasize that this is likely a slight overestimate of the number of transporters needed as there are other phosphorous scavenging systems, such as the ATP-dependent phosphate transporter Pst system which we have neglected.

## Sulfur

Similar to phosphate, sulfate is highly-charged and not particularly membrane permeable, requiring active transport. While there exists a H+/sulfate symporter in *E. coli*, it is in relatively low abundance and is not well characterized (***Zhang et al., 2014***). Sulfate is predominantly acquired via the ATP-dependent ABC transporter CysUWA system which also plays an important role in selenium transport (***Sekowska et al., 2000***; ***Sirko et al., 1995***). While specific kinetic details of this transport system are not readily available, generic ATP transport systems in prokaryotes transport on the order of 1 to 10 molecules per second (BNID: 109035). Combining this generic transport rate, measurement of sulfur comprising 1% of dry mass, and a 5000 second division time yields an estimate of ≈ 1000 CysUWA complexes per cell (***Figure 2–Figure Supplement 1***(B)). Once again, this estimate is in notable agreement with proteomic data sets, suggesting that there are sufficient transporters present to acquire the necessary sulfur. In a similar spirit of our estimate of phosphorus transport, we emphasize that this is likely an overestimate of the number of necessary transporters as we have neglected other sulfur scavenging systems that are in lower abundance.

## Additional Process of the Central Dogma

In the main text, we consider the processes underlying the backbone of the central dogma, namely DNA replication, RNA transcription, and protein translation. In this section we turn our attention to additional processes related to the central dogma, primarily dNTP synthesis for DNA replication and amino-acyl tRNA synthesis for translation. Additionally, we explore in more detail the estimates shown in ***Figure 8***(A) for the RNA polymerase requirements of mRNA and tRNA synthesis.

## dNTP synthesis

The four major dNTPs (dATP, dTTP, dCTP, and dGTP) serve as the fundamental units of the genetic code. Thus, to faithfully replicate the chromosome, the cell must be able to synthesize enough of these bases in the first place. All dNTPs are synthesized *de novo* in separate pathways, requiring different building blocks. However, a critical step present in all dNTP synthesis pathways is the conversion from ribonucleotide to deoxyribonucleotide via the removal of the 3’ hydroxyl group of the ribose ring (***Rudd et al., 2016***). This reaction is mediated by a class of enzymes termed ribonucleotide reductases, of which *E. coli* possesses two aerobically active complexes (termed I and II) and a single anaerobically active enzyme. Due to their peculiar formation of a radical intermediate, these enzymes have received much biochemical, kinetic, and structural characterization. One such work (***Ge et al., 2003***) performed a detailed *in vitro* measurement of the steady-state kinetic rates of these complexes, revealing a turnover rate of ≈ 10 dNTP per second.

Since this reaction is central to the synthesis of all dNTPs, it is reasonable to consider the abundance of these complexes is a measure of the total dNTP production in *E. coli*. Illustrated schematically in ***Figure 7***(A), we consider the fact that to replicate the cell’s genome, on the order of ≈ 1 × 10^7^ dNTPs must be synthesized. Assuming a production rate of 10 per second per ribonucleotide reductase complex and a cell division time of 5000 seconds, we arrive at an estimate of ≈ 200 complexes needed per cell. As shown in ***Figure 7–Figure Supplement 1***, this estimate agrees with the experimental measurements of these complexes abundances within ≈ 1/2 an order of magnitude. Extension of this estimate across a continuum of growth rate, including the fact that multiple chromosomes can be replicated at a given time, is shown as a grey transparent line in ***Figure 7–Figure Supplement 1***. Similarly to our point estimate, this refinement agrees well with the data, accurately describing both the magnitude of the complex abundance and the dependence on growth rate.

Recent work has revealed that during replication, the ribonucleotide reductase complexes coalesce to form discrete foci colocalized with the DNA replisome complex (***Sánchez-Romero et al., 2011***). This is particularly pronounced in conditions where growth is slow, indicating that spatial organization and regulation of the activity of the complexes plays an important role.

## mRNA and tRNA Synthesis

In ***Figure 8*** of the main text, we presented estimates for the number of RNA polymerases needed to synthesize enough mRNA and tRNA molecules. Here, we present a more detailed rationalization of these estimates.

To form a functional protein, all protein coding genes must first be transcribed from DNA to form an mRNA molecule. While each protein requires an mRNA blueprint, many copies of the protein can be synthesized from a single mRNA. Factors such as strength of the ribosomal binding site, mRNA stability, and rare codon usage frequency dictate the number of proteins that can be made from a single mRNA, with yields ranging from 10^1^ to 10^4^ (BNID: 104186; 100196; 106254). Computing the geometric mean of this range yields ≈ 1000 proteins synthesized per mRNA, a value that agrees with experimental measurements of the number of proteins per cell (≈ 3 × 10^6^, BNID: 100088) and total number of mRNA per cell (≈ 3 × 10^3^, BNID:100064).

This estimation captures the *steady-state* mRNA copy number, meaning that at any given time, there will exist approximately 3000 unique mRNA molecules. To determine the *total* number of mRNA that need to be synthesized over the cell’s lifetime, we must consider degradation of the mRNA. In most bacteria, mRNAs are rather unstable with life times on the order of several minutes (BNID: 104324; 106253; 111927; 111998). For convenience, we assume that the typical mRNA in our cell of interest has a typical lifetime of ≈ 300 seconds. Using this value, we can determine the total mRNA production rate to maintain a steady-state copy number of 3000 mRNA per cell. While we direct the reader to the appendix for a more detailed discussion of mRNA transcriptional dynamics, we state here that the total mRNA production rate must be on the order of ≈ 15 mRNA per second. In *E. coli*, the average protein is ≈ 300 amino acids in length (BNID: 108986), meaning that the corresponding mRNA is ≈ 900 nucleotides which we will further approximate as ≈ 1000 nucleotides to account for the non-protein coding regions on the 5’ and 3’ ends. This means that the cell must have enough RNA polymerase molecules around to sustain a transcription rate of ≈ 1.5 × 10^4^ nucleotides per second. Knowing that a single RNA polymerase polymerizes RNA at a clip of 40 nucleotides per second, we arrive at a comfortable estimate of ≈ 250 RNA polymerase complexes needed to satisfy the mRNA demands of the cell. It is worth noting that this number is approximately half of that required to synthesize enough rRNA, as we saw in the previous section. We find this to be a striking result as these 250 RNA polymerase molecules are responsible for the transcription of the ≈ 4000 protein coding genes that are not ribosome associated.

We now turn our attention to the synthesis of tRNA. Unlike mRNA or rRNA, each individual tRNA is remarkably short, ranging from 70 to 95 nucleotides each (BNID: 109645; 102340). What they lack in length, they make up for in abundance, with reported values ranging from ≈ 5 ×10^4^ (BNID: 105280) to ≈ 5 ×10^5^ (BNID: 108611). To test tRNA synthesis as a possible growth-rate limiting stage, we will err towards a higher abundance of ≈ 5×10^5^ per cell. Combining the abundance and tRNA length measurements, we make the estimate that ≈ 5 × 10^7^ nucleotides are sequestered in tRNA per cell. Unlike mRNA, tRNA is remarkably stable with typical lifetimes *in vivo* on the order of ≈ 48 hours (***Abelson et al., 1974***; ***Svenningsen et al., 2017***) – well beyond the timescale of division. Once again using our rule-of-thumb for the rate of transcription to be 40 nucleotides per second and assuming a division time of ≈ 5000 seconds, we arrive at an estimate of ≈ 200 RNA polymerases to synthesize enough tRNA. This requirement pales in comparison to the number of polymerases needed to generate the rRNA and mRNA pools and can be neglected as a significant transcriptional burden.

## tRNA Charging

In the previous subsection, we focused solely on estimating the number of RNA polymerases needed for the generation of the tRNA molecule itself. We now explore the protein complex requirements for ligation of the appropriate amino acid to each tRNA. We begin by again using an estimate of ≈ ≈ 3 × 10^6^ proteins per cell at a 5000 s division time (BNID: 115702) and a typical protein length of ≈ 300 amino acids (BNID: 100017), we can estimate that a total of ≈ 1 × 10^9^ amino acids are stitched together by peptide bonds.

How many tRNAs are needed to facilitate this remarkable number of amino acid delivery events to the translating ribosomes? It is important to note that tRNAs are recycled after they’ve passed through the ribosome and can be recharged with a new amino acid, ready for another round of peptide bond formation. While some *in vitro* data exists on the turnover of tRNA in *E. coli* for different amino acids, we can make a reasonable estimate by comparing the number of amino acids to be polymerized to cell division time. Using our stopwatch of 5000 s and ≈ 1 × 10^9^ amino acids, we arrive at a requirement of ≈ 2 × 10^5^ tRNA molecules to be consumed by the ribosome per second.

There are many processes which go into synthesizing a tRNA and ligating it with the appropriate amino acids. As we discussed previously, there appear to be more than enough RNA polymerases per cell to synthesize the needed pool of tRNAs. Without considering the many ways in which amino acids can be scavenged or synthesized *de novo*, we can explore ligation the as a potential rate limiting step. The enzymes which link the correct amino acid to the tRNA, known as tRNA synthetases or tRNA ligases, are incredible in their proofreading of substrates with the incorrect amino acid being ligated once out of every 10^4^ to 10^5^ events (BNID: 103469). This is due in part to the consumption of energy as well as a multi-step pathway to ligation. While the rate at which tRNA is ligated is highly dependent on the identity of the amino acid, it is reasonable to state that the typical tRNA synthetase has charging rate of ≈ 20 AA per tRNA synthetase per second (BNID: 105279).

We can make an assumption that amino-acyl tRNAs are in steady-state where they are produced at the same rate they are consumed, meaning that 2 × 10^5^ tRNAs must be charged per second. Combining these estimates together, as shown schematically in ***Figure 9–Figure Supplement 1***, yields an estimate of ≈ 1 × 10^4^ tRNA synthetases per cell with a division time of 5000 s. This point estimate is in very close agreement with the observed number of synthetases (the sum of all 20 tRNA synthetases in *E. coli*). This estimation strategy seems to adequately describe the observed growth rate dependence of the tRNA synthetase copy number (shown as the grey line in ***Figure 9–Figure Supplement 1***, suggesting that the copy number scales with the cell volume.

In total, the estimated and observed ≈ 1 × 10^4^ tRNA synthetases occupy only a meager fraction of the total cell proteome, around 0.5% by abundance. It is reasonable to assume that if tRNA charging was a rate limiting process, cells would be able to increase their growth rate by devoting more cellular resources to making more tRNA synthases. As the synthesis of tRNAs and the corresponding charging can be highly parallelized, we can argue that tRNA charging is not a rate limiting step in cell division, at least for the growth conditions explored in this work.

## Experimental Details Behind Proteomic Data

Here we provide a brief summary of the experiments behind each proteomic data set considered. The purpose of this section is to identify how the authors arrived at absolute protein abundances. In the following section (see section on Summary of Proteomic Data) we will then provide a summary of the protein abundance measurements. Table A1 provides an overview of the publications we considered. These are predominately mass spectrometry-based, with the exception of the work from ***Li et al.*** (***2014***) which used ribosomal profiling, and the fluorescence-based counting done in ***Taniguchi et al.*** (***2010***). After having compiled and comparing these measurements, we noted substantial deviations in the measurements from ***Taniguchi et al.*** (***2010***) and ***Soui et al.*** (***2015***) (shown in the following section), and decided to only use the data from ***Schmidt et al.*** (***2016***); ***Li et al.*** (***2014***); ***Valgepea et al.*** (***2013***); ***Peebo et al.*** (***2015***) in the main text. For completeness, we include these additional datasets in our discussion of the experimental data.

**Table A1.** Overview of proteomic data sets.

| Author | Method | Reported Quantity |
| --- | --- | --- |
| Taniguchi *et al.* (2010) | YFP-fusion, cell fluorescence | fg/copies per cell |
| Valgepea *et al.* (2012) | mass spectrometry | fg/copies per cell |
| Peebo *et al.* (2014) | mass spectrometry | fg/copies per fL |
| Li *et al.* (2014) | ribosomal profiling | fg/copies per cell <sup>a</sup> |
| Soufi *et al.* (2015) | mass spectrometry | fg/copies per cell |
| Schmidt *et al.* (2016) | mass spectrometry | fg/copies per cell <sup>b</sup> |

a. The reported values assume that the proteins are long-lived compared to the generation time but are unable to account for post-translational modifications that may alter absolute protein abundances.
b. This mass spectrometry approach differs substantially from the others since in addition to the relative proteome-wide abundance measurements, the authors performed absolute quantification of 41 proteins across all growth conditions (see section on Additional Considerations of Schmidt *et al.* Data Set for more details on this).

## Fluorescence based measurements

In the work of ***Taniguchi et al.*** (***2010***), the authors used a chromosomal YFP fusion library where individual strains have a specific gene tagged with a YFP-coding sequence. 1018 of their 1400 attempted strains were used in the work. A fluorescence microscope was used to collect cellular YFP intensities across all these strains. Through automated image analysis, the authors normalized intensity measurements by cell size to account for the change in size and expression variability across the cell cycle. Following correction of YFP intensities for cellular autofluorescence, final absolute protein levels were determined by a calibration curve with single-molecule fluorescence intensities. This calibration experiment was performed separately using a purified YFP solution.

## Ribosomal profiling measurements

The work of ***Li et al.*** (***2014***) takes a sequencing based approach to estimate protein abundance. Ribosomal profiling, which refers to the deep sequencing of ribosome-protected mRNA fragments, can provide a quantitative measurement of the protein synthesis rate. As long as the protein life-time is long relative to the cell doubling time, it is possible to estimate absolute protein copy numbers. The absolute protein synthesis rate has units of proteins per generation, and for stable proteins will also correspond to the protein copy number per cell.

In the experiments, ribosome-protected mRNA is extracted from cell lysate and selected on a denaturing polyacrylamide gel for deep sequencing (15–45 nt long fragments collected and sequenced by using an Illumina HiSeq 2000 in ***Li et al.*** (***2014***)). Counts of ribosome footprints from the sequencing data were then corrected empirically for position-dependent biases in ribosomal density across each gene, as well as dependencies on specific sequences including the Shine-Dalgarno sequence. These data-corrected ribosome densities represent relative protein synthesis rates. Absolute protein synthesis rates are obtained by multiplying the relative rates by the total cellular protein per cell. The total protein per unit volume was determined with the Lowry method to quantify total protein, calibrated against bovine serum albumin (BSA). By counting colony-forming units following serial dilution of their cell cultures, they then calculated the total protein per cell.

## Mass spectrometry measurements

Perhaps not surprisingly, the data is predominantly mass spectrometry based. This is largely due to tremendous improvements in the sensitivity of mass spectrometers, as well as improvements in sample preparation and data analysis pipelines. It is now a relatively routine task to extract protein from a cell and quantify the majority of proteins present by shotgun proteomics. In general, this involves lysing cells, enzymatically digesting the proteins into short peptide fragments, and then introducing them into the mass spectrometer (e.g. with liquid chromatography and electrospray ionization), which itself can have multiple rounds of detection and further fragmentation of the peptides.

Most quantitative experiments rely on labeling protein with stable isotopes, which allow multiple samples to be measured together by the mass spectrometer. By measuring samples of known total protein abundance simultaneously (i.e. one sample of interest, and one reference), it is possible to determine relative protein abundances. Absolute protein abundances can be estimated following the same approach used above for ribosomal profiling, which is to multiply each relative abundance measurement by the total cellular protein per cell. This is the approach taken by ***Valgepea et al.*** (***2013***); ***Peebo et al.*** (***2015***) and ***Soui et al.*** (***2015***), with relative protein abundances determined based on the relative peptide intensities (label free quantification ‘LFQ’ intensities). For the data of ***Valgepea et al.*** (***2013***), total protein per cell was determined by measuring total protein by the Lowry method, and counting colony-forming units following serial dilution. For the data from ***Peebo et al.*** (***2015***), the authors did not determine cell quantities and instead report the cellular protein abundances in protein per unit volume by assuming a mass density of 1.1 g/ml, with a 30% dry mass fraction.

An alternative way to arrive at absolute protein abundances is to dope in synthetic peptide fragments of known abundance. These can serve as a direct way to calibrate mass spectrometry signal intensities to absolute mass. This is the approach taken by ***Schmidt et al.*** (***2016***). In addition to a set of shotgun proteomic measurements to determine proteome-wide relative abundances, the authors also performed absolute quantification of 41 proteins covering over four orders of magnitude in cellular abundance. Here, a synthetic peptide was generated for each of the proteins, doped into each protein sample, and used these to determine absolute protein abundances of the 41 proteins. These absolute measurements, determined for every growth condition, were then used as a calibration curve to convert proteomic-wide relative abundances into absolute protein abundance per cell. A more extensive discussion of the ***Schmidt et al.*** (***2016***) data set can be found in Section Additional Considerations of Schmidt *et al.* Data Set.

## Summary of Proteomic Data

In the work of the main text we only used the data from ***Valgepea et al.*** (***2013***); ***Li et al.*** (***2014***); ***Peebo et al.*** (***2015***); ***Schmidt et al.*** (***2016***). As shown in ***Figure A1***(A), the reported total protein abundances in the work of ***Taniguchi et al.*** (***2010***) and ***Soui et al.*** (***2015***) differed quite substantially from the other work. For the work of ***Taniguchi et al.*** (***2010***) this is in part due to a lower coverage in total proteomic mass quantified, though we also noticed that most proteins appear undercounted when compared to the other data.

***Figure A1***(B) summarizes the total protein mass for each data set used in our final compiled data set. Our inclination initally was to leave reported copy numbers untouched, but a notable descrepency between the scaling of the total protein per cell between ***Schmidt et al.*** (***2016***) and the other data sets forced us to dig deeper into those measurements (compare ***Schmidt et al.*** (***2016***) and ***Li et al.*** (***2014***) data in ***Figure A1***(A)). The particular trend in ***Schmidt et al.*** (***2016***) appears to be due to assumptions made about cell size and we provide a more extensive discussion and analysis of their data in Additional Considerations of Schmidt *et al.* Data Set. As a compromise, and in an effort to treat all data equally, we instead applied an correction factor to all protein abundance values based on a data-driven estimate of total protein per cell. Here we used cell size measurements from ***Si et al.*** (***2017***, 2019), and an estimate of total protein content through expected dry mass. Total protein per cell was then determined using available data on total DNA, RNA, and protein from ***Basan et al.*** (***2015***); ***Dai et al.*** (***2016***), which account for the majority of dry mass in the cell. We describe these details further in sections on Estimation of Cell Size and Surface Area and Estimation of Total Protein Content per Cell that follows.

Lastly, in ***Figure A2*** we show the total proteomic coverage and overlap of proteins quantified across each data set. In the horizontal bar plot (***Figure A2***, bottom left) we plot the total number of unique proteins from each data set, while in the main plot we show the intersections across each data set. Overall, the overlap in quantified proteins is quite high, with 1157 proteins quantified across all data sets. The sequencing based approach of ***Li et al.*** (***2014***) has substantially higher coverage compared to the mass spectrometry data sets (3394 genes versus the 2041 genes quantified in the work of ***Schmidt et al.*** (***2016***)). However, in terms of total protein mass, the data from ***Li et al.*** (***2014***); ***Schmidt et al.*** (***2016***); ***Peebo et al.*** (***2015***) each quantify roughly equivalent total protein mass. An exception to this is in the data from ***Valgepea et al.*** (***2013***), where we find that the total protein quantified in ***Valgepea et al.*** (***2013***) is 90-95 % of the total protein mass (when using the data from ***Schmidt et al.*** (***2016***) as a reference).

**Figure A1.**
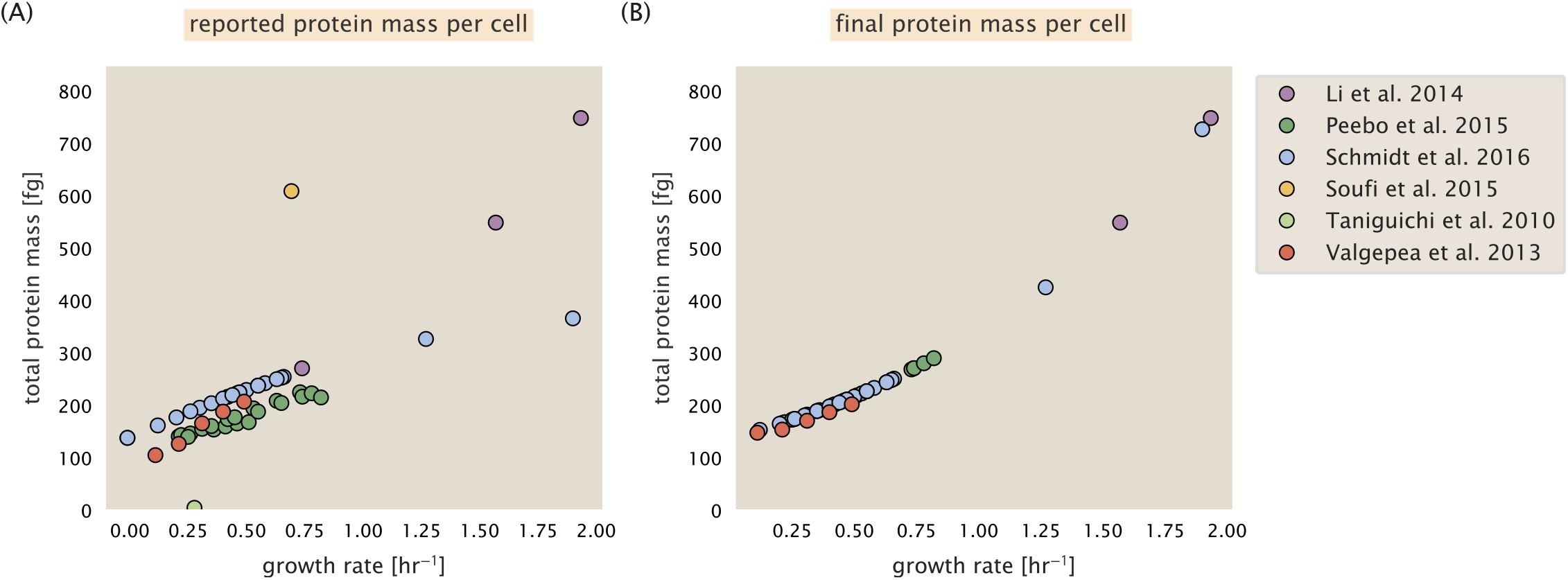
Summary of the growth-rate dependent total protein abundance for each data set. (A) Total protein abundance per cell as originally reported in the data sets of ***Taniguchi et al.*** (***2010***); ***Valgepea et al.*** (***2013***); ***Li et al.*** (***2014***); ***Soui et al.*** (***2015***); ***Peebo et al.*** (***2015***); ***Schmidt et al.*** (***2016***). Note that the data from ***Peebo et al.*** (***2015***) only reported protein abundances per unit volume and total protein per cell was found by multiplying these by the growth-rate dependent cell size as determined by ***Si et al.*** (***2017***). (B) Adjusted total protein abundances across the proteomic data sets are summarized. Protein abundances were adjusted so that all data shared a common set of growth-rate dependent total protein per cell and cellular protein concentration following the cell size expectations of ***Si et al.*** (***2017***) (see section on Estimation of Cell Size and Surface Area for further details).

**Figure A2.**
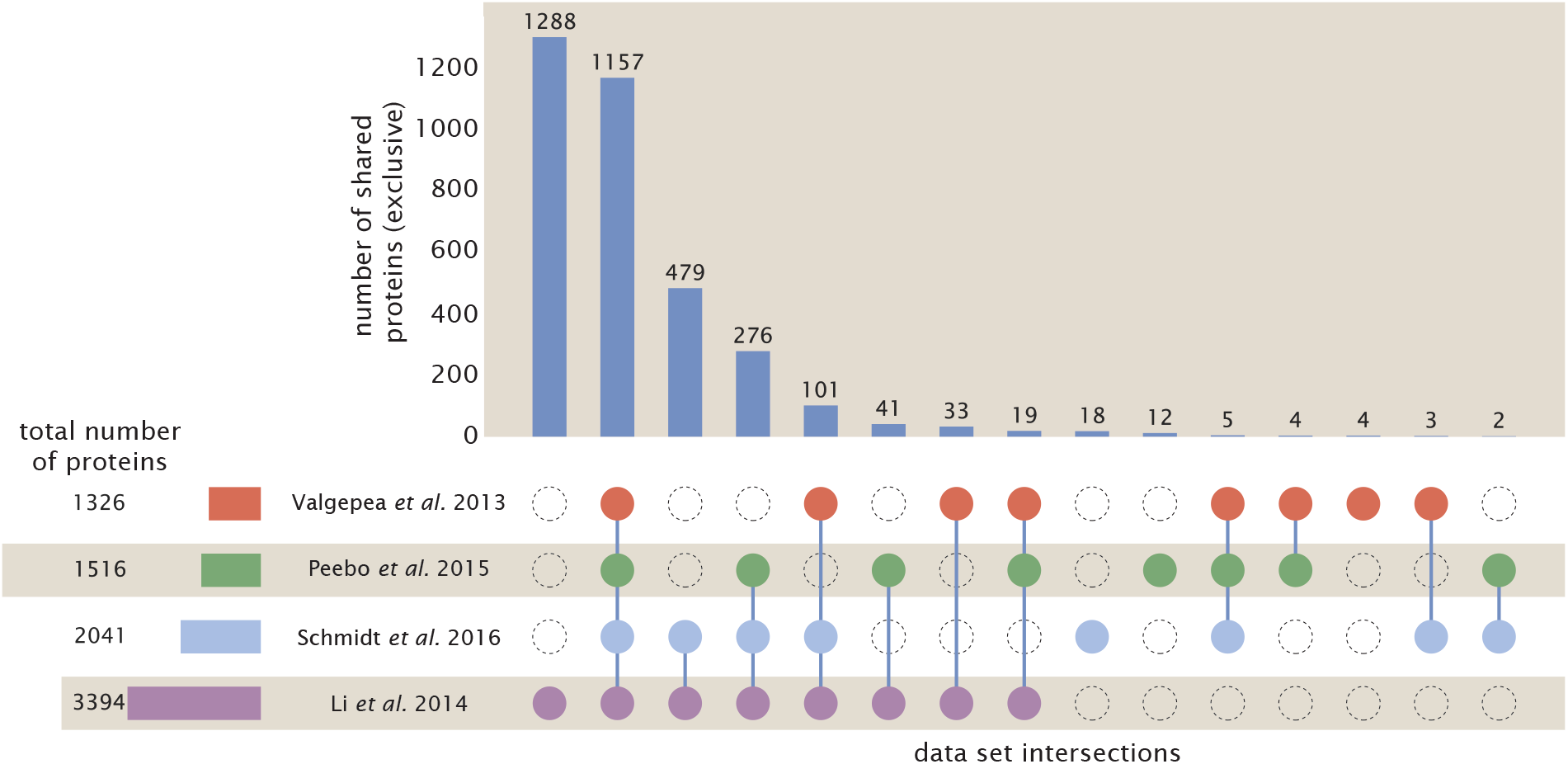
Comparison of proteomic coverage across diferent data sets. An UpSet diagram (***Lex et al., 2014***) summarizes the total number of protein coding genes whose protein abundance was reported in the data sets of ***Valgepea et al.*** (***2013***); ***Li et al.*** (***2014***); ***Schmidt et al.*** (***2016***); ***Peebo et al.*** (***2015***). Bar plot on bottom left indicates the total number of genes reported in each individual data. The main bar plot summarizes the number of unique proteins identified across overlapping subsets of the data. For example, in the first column only the data from Li et al. (2014) is considered (indicated by solid purple circle) and 1288 proteins are identified as exclusive to the data set. In the second column, the intersection of all four data sets is considered, with 1157 proteins quantified across them. This follows for each additional column in the plot, with the subset under consideration denoted by the solid colored circles.

**Figure A3.**
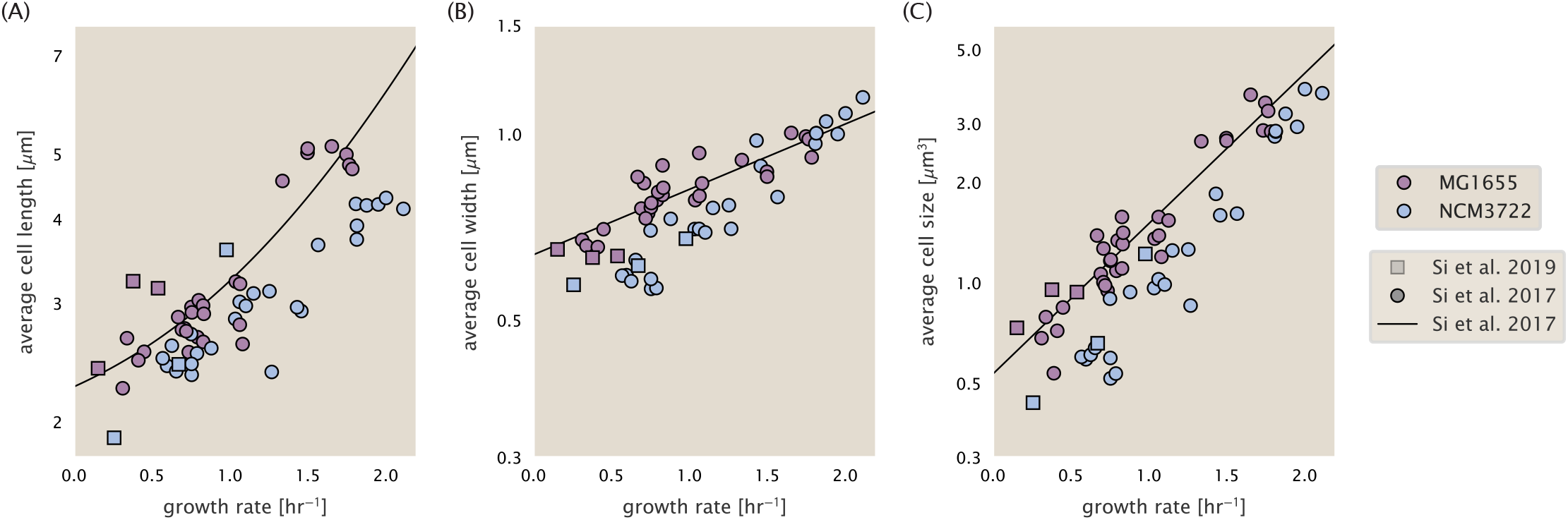
Summary of size measurements from Si *et al.* 2017, 2019. Cell lengths and widths were measured from cell contours obtained from phase contrast images, and refer to the long and short axis respectively. (A) Cell lengths and (B) cell widths show the mean measurements reported (they report 140-300 images and 5,000-30,000 for each set of samples; which likely means about 1,000-5,000 measurements per mean value reported here since they considered about 6 conditions at a time). Fits were made to the MG1655 strain data; length: 0.5 *e*^1.09.*λ*^ + 1.76 μm, width: 0.64 *e*^0.24.*λ*^ μm. (C) Cell size was calculated as cylinders with two hemispherical ends (Equation 1). The MG1655 strain data gave a best fit of 0.533 *e*^1.037.*λ*^ μm^3^.

## Estimation of Cell Size and Surface Area

Since most of the proteomic data sets lack cell size measurements, we chose instead to use a common estimate of size for any analysis requiring cell size or surface area. Since each of the data sets used either K-12 MG1655 or its derivative, BW25113 (from the lab of Barry L. Wanner; the parent strain of the Keio collection (***Datsenko and Wanner, 2000***; ***Baba et al., 2006***)), below we fit the MG1655 cell size data from the supplemental material of ***Si et al.*** (***2017***, 2019) using non-linear least squares regression as implemented by the `optimize.curve_fit` function from the Scipy Python package (***Virtanen et al., 2020***). Throughout the text, we usually refer to cell size, in units of μm^3^; however, on occasion we will mention size as a volume in units of fL.

The average size measurements from each of their experiments are shown in ***Figure A3***, with cell length and width shown in (A) and (B), respectively. The length data was well described by the exponential function 0.5 e^1.09·*λ*^ + 1.76 μm, while the width data was well described by 0.64 *e*^0.24.*λ*^ μm. In order to estimate cell size we take the cell as a cylinder with two hemispherical ends (***Si et al., 2017***; ***Basan et al., 2015***). Specifically, cell size is estimated from,

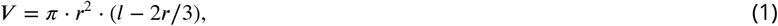

where *r* is half the cell width. A best fit to the data is described by 0.533 *e*^1.037.*λ*^ μm^3^. Calculation of the cell surface area is given by,

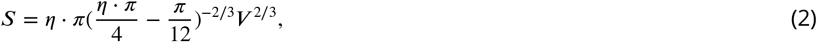

where η is the aspect ratio (*η* = *l/w*) (***Ojkic et al., 2019***).

## Estimation of Total Protein Content per Cell

In order to estimate total protein per cell for a particular growth rate, we begin by estimating the cell size from the fit shown in ***Figure A3***(C) (cell size = 0.533 *e*^1.037.*λ*^ μm^3^, as noted in the previous section). We then estimate the total protein content from the total dry mass of the cell. Here we begin by noting that for almost the entire range of growth rates considered here, protein, DNA, and RNA were reported to account for at least 90 % of the dry mass (***Basan et al.*** (***2015***)). The authors also found that the total dry mass concentration was roughly constant across growth conditions. Under such a scenario, we can calculate the total dry mass concentration for protein, DNA, and RNA, which is given by 1.1 g/ml × 30 % × 90 % or about [*M_P_*] = 300 fg per fL. Multiplying this by our prediction of cell size gives the total dry mass per cell.

However, even if dry mass concentration is relatively constant across growth conditions, it is not obvious how protein concentration might vary due to the substantial increase in rRNA at faster growth rates (***Dai et al.*** (***2016***)). The increase in rRNA increases from the linear increase in ribosomal content with faster growth rate (***Scott et al.*** (***2010***)), since it makes up about about 2/3 or the ribosomal mass. To proceed we therefore relied on experimental measurements of total DNA content per cell from ***Basan et al.*** (***2015***), and RNA to protein ratios that were measured in Dai *et al.* (and cover the entire range of growth conditions considered here). These are reproduced in ***Figure A4***(A) and (B), respectively.

Assuming that the protein, DNA, and RNA account for 90 % of the total dry mass, the protein mass can then determined by first subtracting the experimentally measured DNA mass, and then using the experimental estimate of the RNA to protein ratio. The total protein per cell is will be related to the summed RNA and protein mass by,

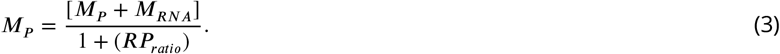

(*RP_ratio_* refers to the RNA to protein ratio as measured by Dai *et al.*. In ***Figure A4***(C) we plot the estimated cellular concentrations for protein, DNA, and RNA from these calculations, and in ***Figure A4***(D) we plot their total expected mass per cell. This later quantity is the growth rate-dependent total protein mass that was used to estimate total protein abundance across all data sets (and summarized in ***Figure A1***(B)).

## Estimating Volume and Number of Amino Acids from Ribosome Copy Number

Towards the end of the main text, we examine a coarse-grained model of nutrient-limited growth. A key point in our analysis was to consider how elongation rate *r*_*t*_ and growth rate *λ* vary with respect to the experimentally observed changes in cell size, total number of peptide bonds per cell *N*_pep_, and ribosomal content. In order to restrict parameters to those observed experimentally, but otherwise allow us to explore the model, we performed a phenomenological fit of *N*_pep_ and *V* as a function of the measured ribosomal copy number *R*. As has been described in the preceding sections of this supplement, we estimate cell volume for each growth condition using the size measurements from ***Si et al.*** (***2017***, 2019), and *N*_pep_ is approximated by taking the total protein mass and dividing this number by the average mass of an amino acid, 110 Da (BNID: 104877).

Given the exponential scaling of *V* and *N*_pep_ with growth rate, we performed a linear regression of the log transform of these parameters as a function of the log transform of the ribosome copy number. Using optimization by minimization, we estimated the best-fit values of the intercept and slope for each regression. ***Figure A5*** shows the result of each regression as a dashed line.

## Additional Considerations of Schmidt *et al.* Data Set

While the data set from ***Schmidt et al.*** (***2016***) remains a heroic effort that our labs continue to return to as a resource, there were steps taken in their calculation of protein copy number that we argue needed further consideration. In particular, the authors made an assumption of constant cellular protein concentration across all growth conditions and used measurements of cell volume that appear inconsistent with an expected exponential scaling of cell size with growth rate that is well-documented in *E. coli* (***Schaechter et al.*** (***1958***); ***Taheri-Araghi et al.*** (***2015***); ***Si et al.*** (***2017***)).

We begin by looking at their cell volume measurements, which are shown in blue in Figure ***Figure A6***. As a comparison, we also plot cell sizes reported in three other recent papers: measurements from Taheri-Araghi *et al.* and Si *et al.* come from the lab of Suckjoon Jun, while those from Basan *et al.* come from the lab of Terence Hwa. Each set of measurements used microscopy and cell segmentation to determine the length and width, and then calculated cell size by treating the cell as a cylinder with two hemispherical ends, as we considered in the previous section. While there is notable discrepancy between the two research groups, which are both using strain NCM3722, Basan *et al.* found that this came specifically from uncertainty in determining the cell width. This is prone to inaccuracy given the small cell size and optical resolution limits (further described in their supplemental text). Perhaps the more concerning point is that while each of these alternative measurements show an exponential increase in cell size at faster growth rates, the measurements used by Schmidt *et al.* appear to plateau. This resulted in an analogous trend in their final reported total cellular protein per cell as shown in ***Figure A7*** (purple data points), and is in disagreement with other measurements of total protein at these growth rates (***Basan et al., 2015***).

Since it is not obvious how measurements of cell size influenced their reported protein abundances, in the following subsections we begin by considering how the authors determined total protein mass per cell. We then consider three different approaches to estimate the growth-rate dependent total protein mass and compare these estimates with those reported by ***Schmidt et al.*** (***2016***). Those results are summarized in ***Figure A6***(B), with the original values from both ***Schmidt et al.*** (***2016***) and ***Li et al.*** (***2014***) shown in ***Figure A6***(A) for reference. For most growth conditions, we find reasonable agreement between our estimates and the reported total protein per cell. However, for the fastest growth conditions, with glycerol + supplemented amino acids, and LB media, all estimates are substantially higher than those originally reported. This is the main reason why we chose to readjuste protein abundance as shown in ***Figure A1***(B) (with the calculation described in section Estimation of Total Protein Content per Cell).

**Figure A4.**
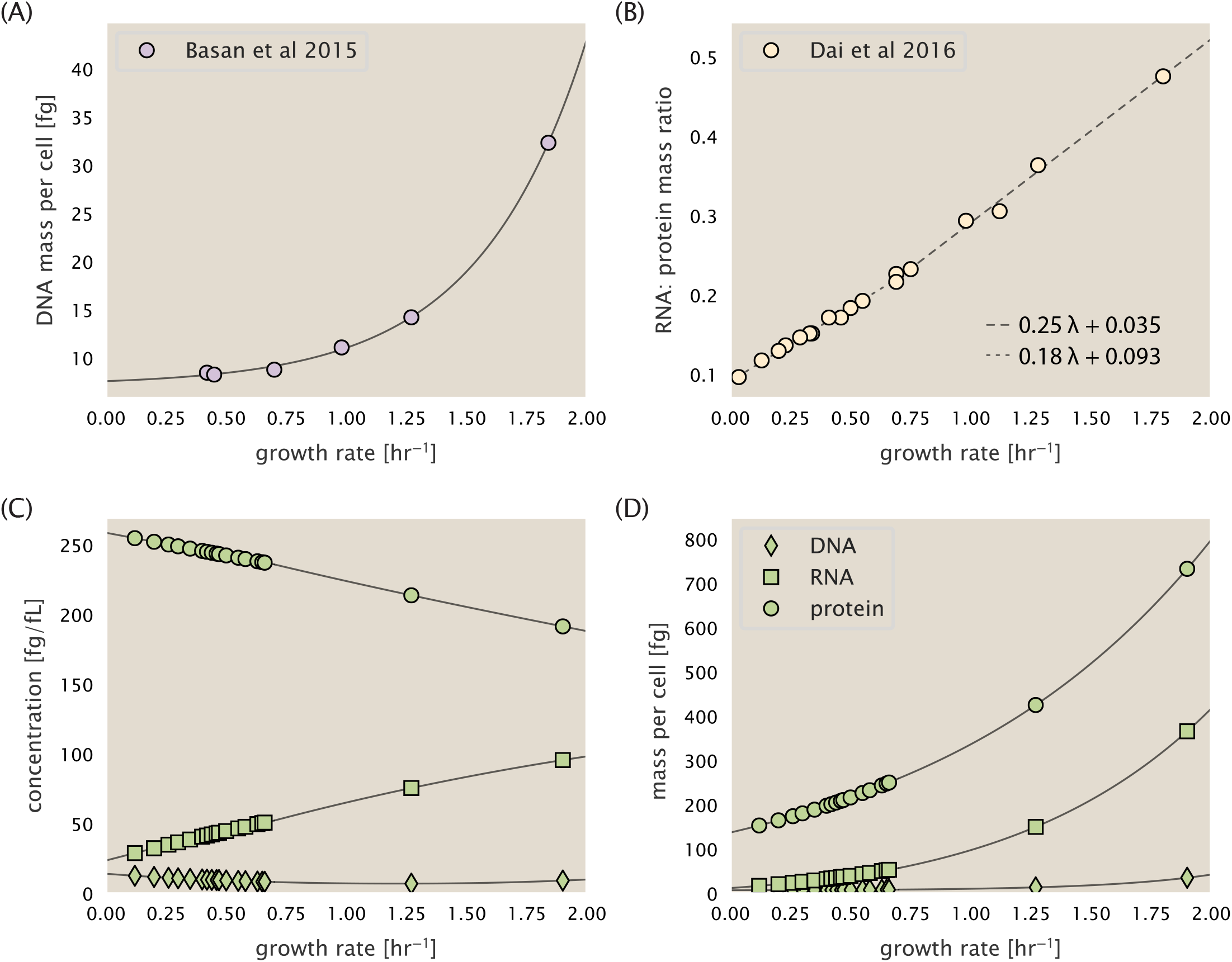
Empirical estimate of cellular protein, DNA, and RNA as a function of growth rate. (A) Measured DNA mass per cell as a function of growth rate, reproduced from Basan *et al.* 2015. The data was fit to an exponential curve (DNA mass in fg per cell is given by 0.42 *e*^2.23.*λ*^ + 7.2 fg per cell, where *λ* is the growth rate in hr^−1^). (B) RNA to protein measurements as a function of growth rate. The data was fit to two lines (shown in black) due to the change in slope at slower growth rates ***Neidhardt et al.*** (***1991***); ***Dai et al.*** (***2016***). For growth rates below 0.7 hr^−1^, the RNA/protein ratio is 0.18.*λ* + 0.093, while for growth rates faster than 0.7 hr^−1^ the RNA/protein ratio is given by 0.25.*λ* + 0.035. For (A) and (B) cells are grown under varying levels of nutrient limitation, with cells grown in minimal media with different carbon sources for the slowest growth conditions, and rich-defined media for fast growth rates. (C) Estimation of cellular protein, DNA, and RNA concentration. (D) Total cellular mass estimated for protein, DNA, and RNA using the cell size calculated in Estimation of Cell Size and Surface Area. Symbols (diamond: DNA, square: RNA, circle: protein) show estimated values of mass concentration and mass per cell for the specific growth rates in ***Schmidt et al.*** (***2016***).

**Figure A5.**
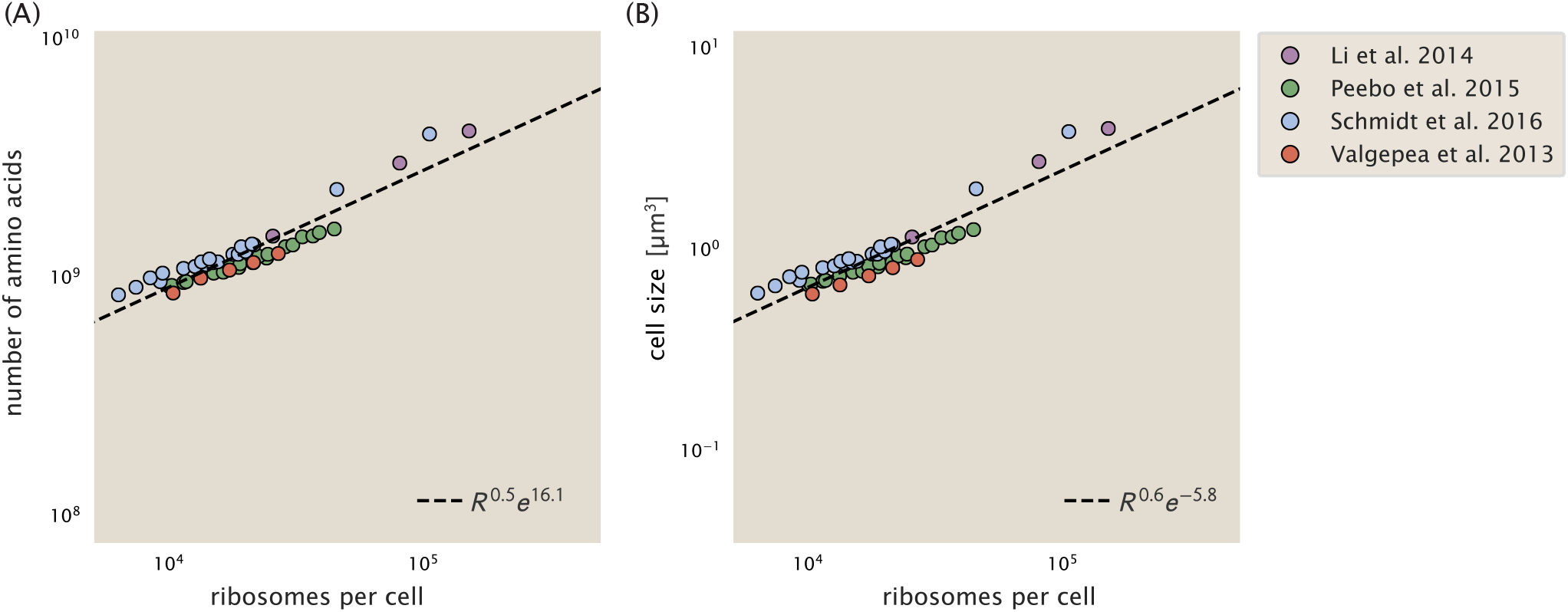
Phenomenological regression of cell volume and number of amino acids per cell as a function of the ribosome copy number. (A) Estimated total number of peptide bonds per cell *N*_pep_ as a function of number of ribosomes per cell. (B) Estimated cell size as described in Estimation of Cell Size and Surface Area, as a function of number of ribosomes per cell. Colored points correspond to the measured value (or calculated value in the case of the cell size) with colors denoting different data sets. The dashed black line shows the result of the fit, with the functional form of the equation given in the legend with R representing the ribosome copy number.

**Figure A6.**
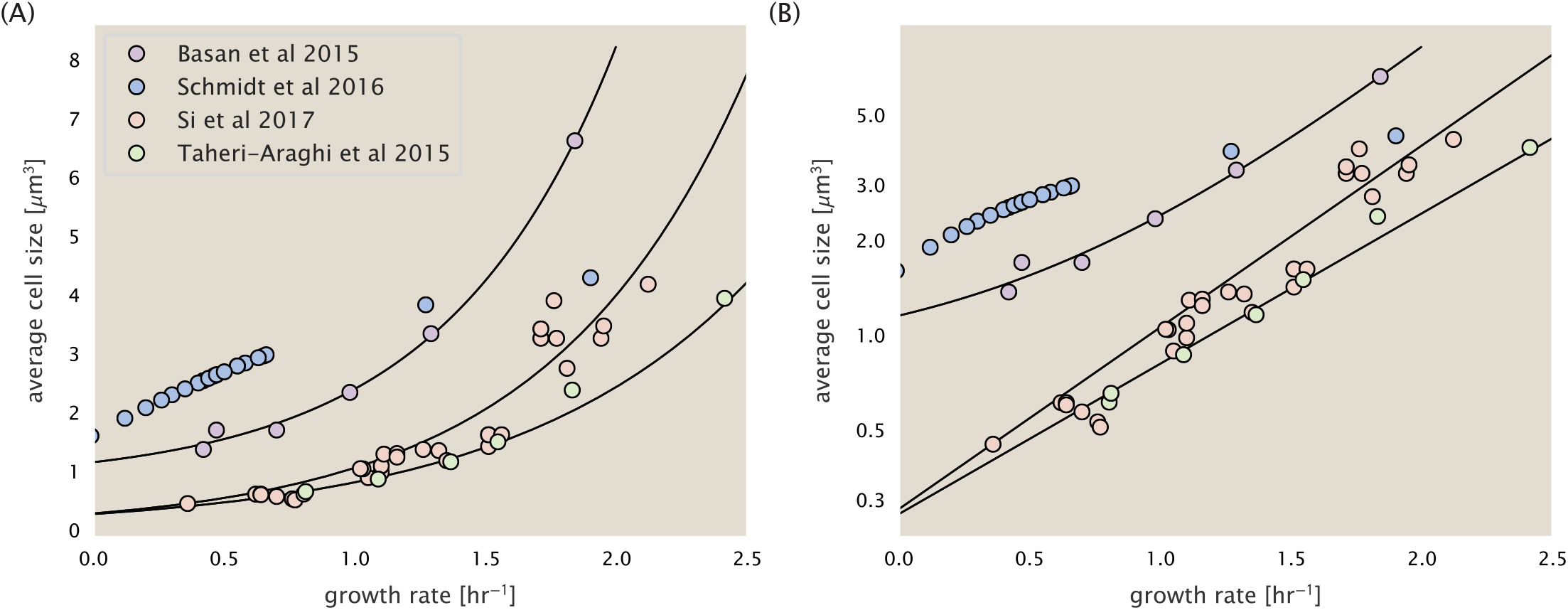
Measurements of cell size as a function of growth rate. (A) Plot of the reported cell sizes from several recent papers. The data in blue come from Volkmer and Heinemann, 2011 (***Volkmer and Heinemann*** (***2011***)) and were used in the work of Schmidt *et al.*. Data from the lab of Terence Hwa are shown in purple (***Basan et al.*** (***2015***)), while the two data sets shown in green and light red come from the lab of Suckjoon Jun (***Taheri-Araghi et al.*** (***2015***); ***Si et al.*** (***2017***)). (B) Same as in (A) but with the data plotted on a logarithmic y-axis to highlight the exponential scaling that is expected for *E. coli*.

**Figure A7.**
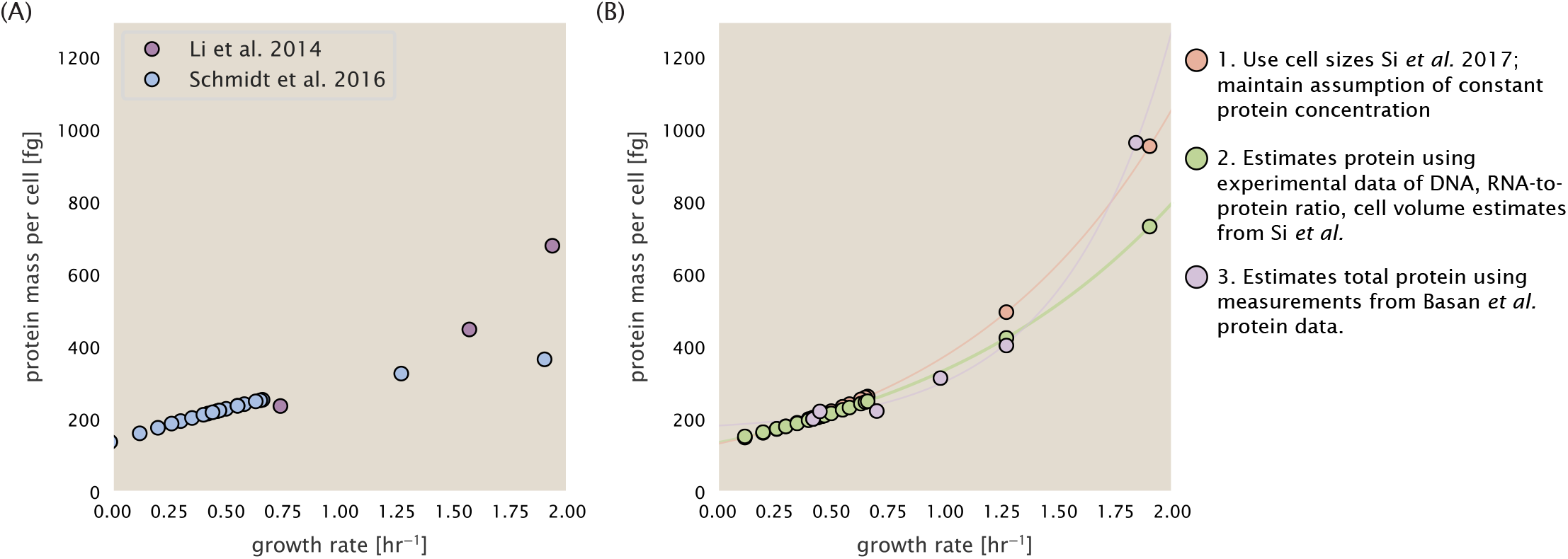
Alternative estimates of total cellular protein for the growth conditions considered in Schmidt ***et al.*** (A) The original protein mass from Schmidt *et al.* and Li *et al.* are shown in purple and blue, respectively. (B) Three alternative estimates of total protein per cell. 1. *light red*: Rescaling of total protein mass assuming a growth rate independent protein concentration and cell volumes estimated from Si *et al.* 2017. 2. *light green*: Rescaling of total protein mass using estimates of growth rate-dependent protein concentrations and cell volumes estimated from Si *et al.* 2017. Total protein per cell is calculated by assuming a 1.1 g/ml cellular mass density, 30% dry mass, with 90% of the dry mass corresponding to DNA, RNA, and protein (***Basan et al., 2015***). See Estimation of Total Protein Content per Cell for details on calculation. 3. *light purple*: Rescaling of total protein mass using the experimental measurements from Basan *et al.* 2015.

## Efect of cell volume on reported absolute protein abundances

As noted in Experimental Details Behind Proteomic Data, the authors from the work in ***Schmidt et al.*** (***2016***) calculated proteomewide protein abundances by first determining absolute abundances of 41 pre-selected proteins, which relied on adding synthetic heavy reference peptides into their protein samples at known abundance. This absolute quantitation was performed in replicate for each growth condition. Separately, the authors also performed a more conventional mass spectrometry measurement for samples from each growth condition, which attempted to maximize the number of quantified proteins but only provided relative abundances based on peptide intensities. Finally, using their 41 proteins with absolute abundances already determined, they then created calibration curves with which to relate their relative intensity to absolute protein abundance for each growth condition. This allowed them to estimate absolute protein abundance for all proteins detected in their proteome-wide data set. Combined with their flow cytometry cell counts, they were then able to determine absolute abundance of each protein detected on a per cell basis.

While this approach provided absolute abundances, another necessary step to arrive at total cellular protein was to account for any protein loss during their various protein extraction steps. Here the authors attempted to determine total protein separately using a BCA protein assay. In personal communications, it was noted that determining reasonable total protein abundances by BCA across their array of growth conditions was particularly troublesome. Instead, they noted confidence in their total protein measurements for cells grown in M9 minimal media + glucose and used this as a reference point with which to estimate the total protein for all other growth conditions.

For cells grown in M9 minimal media + glucose an average total mass of *M_P_* = 240 fg per cell was measured. Using their reported cell volume, reported as *V_orig_* = 2.84 fl, a cellular protein concentration of [*M_P_*]*_orig_* = *M_P_*/*V_orig_* = 85 fg/fl. Now, taking the assumption that cellular protein concentration is relatively independent of growth rate, they could then estimate the total protein mass for all other growth conditions from,

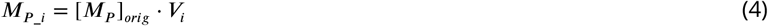

where 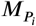 represents the total protein mass per cell and *V_i_* is the cell volume for each growth condition *i* as measured in Volkmer and Heinemann, 2011. Here the thinking is that the values of 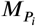 reflects the total cellular protein for growth condition i, where any discrepancy from their absolute protein abundance is assumed to be due to protein loss during sample preparation. The protein abundances from their absolute abundance measurements noted above were therefore scaled to their estimates and are shown in Figure ***Figure A7*** (purple data points).

If we instead consider the cell volumes predicted in the work of Si *et al.*, we again need to take growth in M9 minimal media + glucose as a reference with known total mass, but we can follow a similar approach to estimate total protein mass for all other growth conditions. Letting *V_Si_glu_* = 0.6 fl be the predicted cell volume, the cellular protein concentration becomes [*M_P_*]*_Si_* = *M_P_*/*V_Si_glu_* = 400 fg/fl. The new total protein mass per cell can then be calculated from,

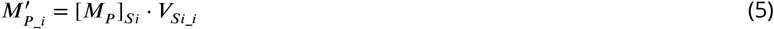

where 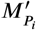 is the new protein mass prediction, and 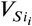 refers to the new volume prediction for each condition *i*, These are shown as red data points in Figure ***Figure A7***(B).

## Relaxing assumption of constant protein concentration across growth conditions

We next relax the assumption that cellular protein concentration is constant and instead, attempt to estimate it using experimental data. Here we use the estimation of total protein mass per cell detailed in Estimation of Total Protein Content per Cell for all data points in the ***Schmidt et al.*** (***2016***) data set. The green data points in ***Figure A7***(B) show this prediction, and this represents the approach used to estimate total protein per cell for all data sets.

## Comparison with total protein measurements from Basan *et al.* 2015.

One of the challenges in our estimates in the preceding sections is the need to estimate protein concentration and cell volumes. These are inherently difficult to measure accurately due to the small size of *E. coli*. Indeed, for all the additional measurements of cell volume included in Figure ***Figure A6***, no measurements were performed for cells growing at rates below 0.5 hr^−1^. It therefore remains to be determined whether our extrapolated cell volume estimates are appropriate, with the possibility that the logarithmic scaling of cell size might break down for slower growth.

In our last approach we therefore attempt to estimate total protein using experimental data that required no estimates of concentration or cell volume. Specifically, in the work of Basan *et al*, the authors measured total protein per cell for a broad range of growth rates (reproduced in Figure ***Figure A8***). These were determined by first measuring bulk protein from cell lysate, measured by the colorimetric Biuret method (***You et al.*** (***2013***)), and then abundance per cell was calculated from cell counts from either plating cells or a Coulter counter. While it is unclear why Schmidt *et al.* was unable to take a similar approach, the results from Basan *et al* appear more consistent with our expectation that cell mass will increase exponentially with faster growth rates. In addition, although they do not consider growth rates below about 0.5 hr^−1^, it is interesting to note that the protein mass per cell appears to plateau to a minimum value at slow growth. In contrast, our estimates using cell volume so far have predicted that total protein mass should continue to decrease slightly for slower growing cells. By fitting this data to an exponential function dependent on growth rate, we could then estimate the total protein per cell for each growth condition considered by ***Schmidt et al.*** (***2016***). These are plotted as red data points in ***Figure A7***(B).

## Calculation of Complex Abundance

All protein data quantified the abundance of individual proteins per cell. However, this work requires estimates on the abundance of individual protein *complexes*, rather than the copy number of individual proteins. In our analysis of the protein copy number data, it became clear that the reported copy numbers do not always align with those based on reported stiochometry. As one example of this, the F-O subunit of ATP synthase consists of three protein subunits with a stiochometry of [AtpB][AtpF]_2_[AtpE]_10_ (also referred to as subunits a, b, and c, respectively). In the experimental data of ***Schmidt et al.*** (***2016***), the values deviate from this quite substantially, with approximately 1000 AtpB, 9000 AtpF, and 300 AtpE reported per cell (minimal media + glucose growth condition). This highlights the technical challenges that still remain in our ability to quantify cellular composition, particularly for membrane-bound proteins like the ATP synthase complex considered here. In this section, we outline the approach we used to annotate proteins as part of each macromolecular complex and how we used averaging across the individual protein measurements to estimate an absolute complex abundances per cell.

Protein complexes, and proteins individually, often have a variety of names, both longform and shorthand. As individual proteins can have a variety of different synonyms, we sought to ensure that each protein annotated in the data sets used the same synonym. To do use, we relied heavily on the EcoCyc Genomic Database (***Keseler et al., 2017***). Each protein in available data sets included an annotation of one of the gene name synonyms as well as an accession ID – either a UniProt or Blattner “b-number". We programmatically matched up individual accession IDs between the proteins in different data sets. In cases where accession IDs matched but the gene names were different, we manually verified that the gene product was the same between the datasets and chose a single synonym. All code used in the data cleaning and unification procedures can be found on the associated [GitHub repository] (DOI:XXX) associated with this paper as well as on the associated paper website.

**Figure A8.**
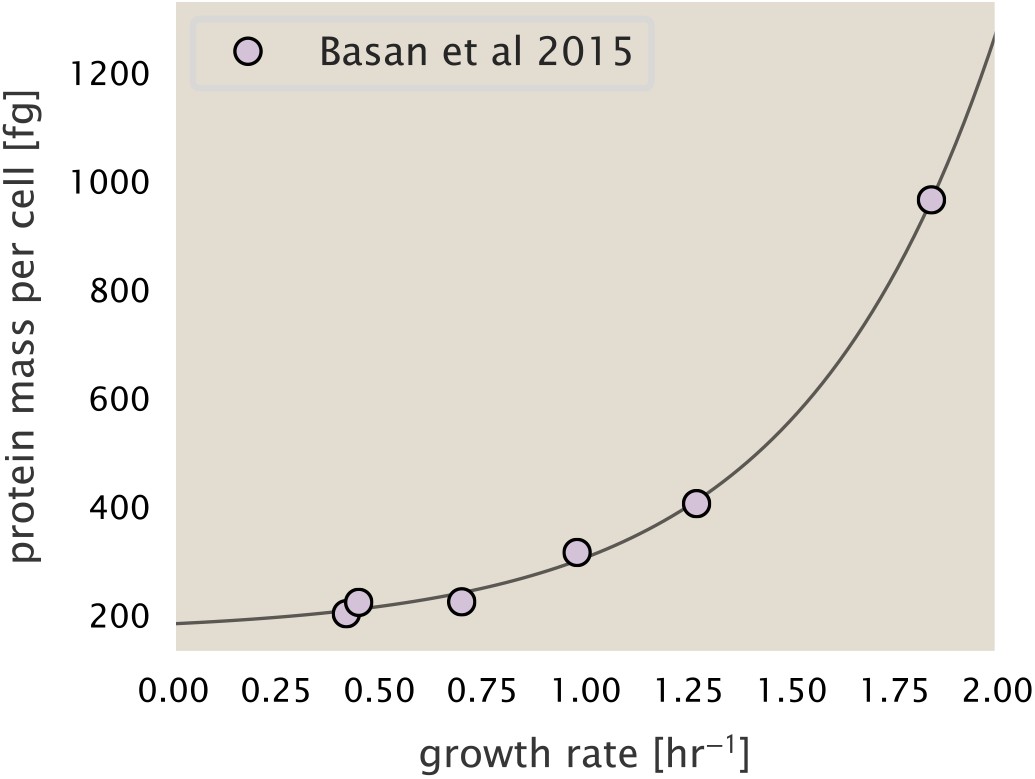
Total cellular protein reported in Basan *et al.* 2015. Measured protein mass as a function of growth rate as reproduced from Basan *et al.* 2015, with cells grown under different levels of nutrient limitation. The data was fit to an exponential curve where protein mass in fg per cell is given by 14.65 *e*^2,180.*λ*^ + 172 fg per cell, where *λ* is the growth rate in hr^−1^).

With each protein conforming to a single identification scheme, we then needed to identify the molecular complexes each protein was a member of. Additionally, we needed to identify how many copies of each protein were present in each complex (i.e. the subunit copy number) and compute the estimated abundance complex that accounted for fluctuations in subunit stoichiometry. To map proteins to complexes, we accessed the EcoCyc *E. coli* database ***Keseler et al.*** (***2017***) using PathwayTools version 23.0 ***Karp et al.*** (***2019***). With a license for PathWay Tools, we mapped each unique protein to its annotated complexes via the BioCyc Python package. As we mapped each protein with *all* of its complex annotations, there was redundancy in the dataset. For example, ribosomal protein L20 (RplT) is annotated to be a component of the 50S ribosome (EcoCyc complex CPLX-03962) as well as a component of the mature 70S ribosome (EcoCyc complex `CPLX-03964`).

In addition to the annotated complex, we collected information on the stoichiometry of each macromolecular complex. For a complex with *N_subunits_* protein species, for each protein subunit *i* we first calculate the number of complexes that *could* be formed given the measured protein copy numbers per cell,

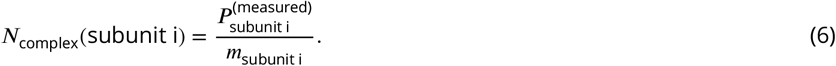

Here, 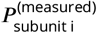 refers to the measured protein copy number of species *i*, and *m* refers to the number of monomers present for that protein in the complex. For example, the 70S mature ribosome complex has 55 protein components, all of which are present in a single copy except L4 (RplL), which is present in 4 copies (*m* = 4). For each ribosomal protein, we then calculate the maximum number of complexes that could be formed using ***Equation 6***. This example, along with example from 5 other macromolecular complexes, can be seen in ***Figure A9***.

It is important to note that measurement noise, efficiency of protein extraction, and differences in protein stability will mean that the precise value of each calculation will be different for each component of a given complex. Thus, to report the total complex abundance, we use the arithmetic mean of across all subunits in the complex, in ***Figure A9***, we show this mean value as a grey line for a variety of different complexes. Additionally, we have built an interactive figure accessible on the paper website where the validity of this approach can be examined for any complex with more than two subunits (thus, excluding monomers and dimers).

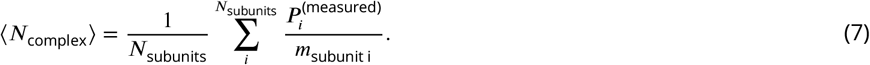

**Figure A9.**
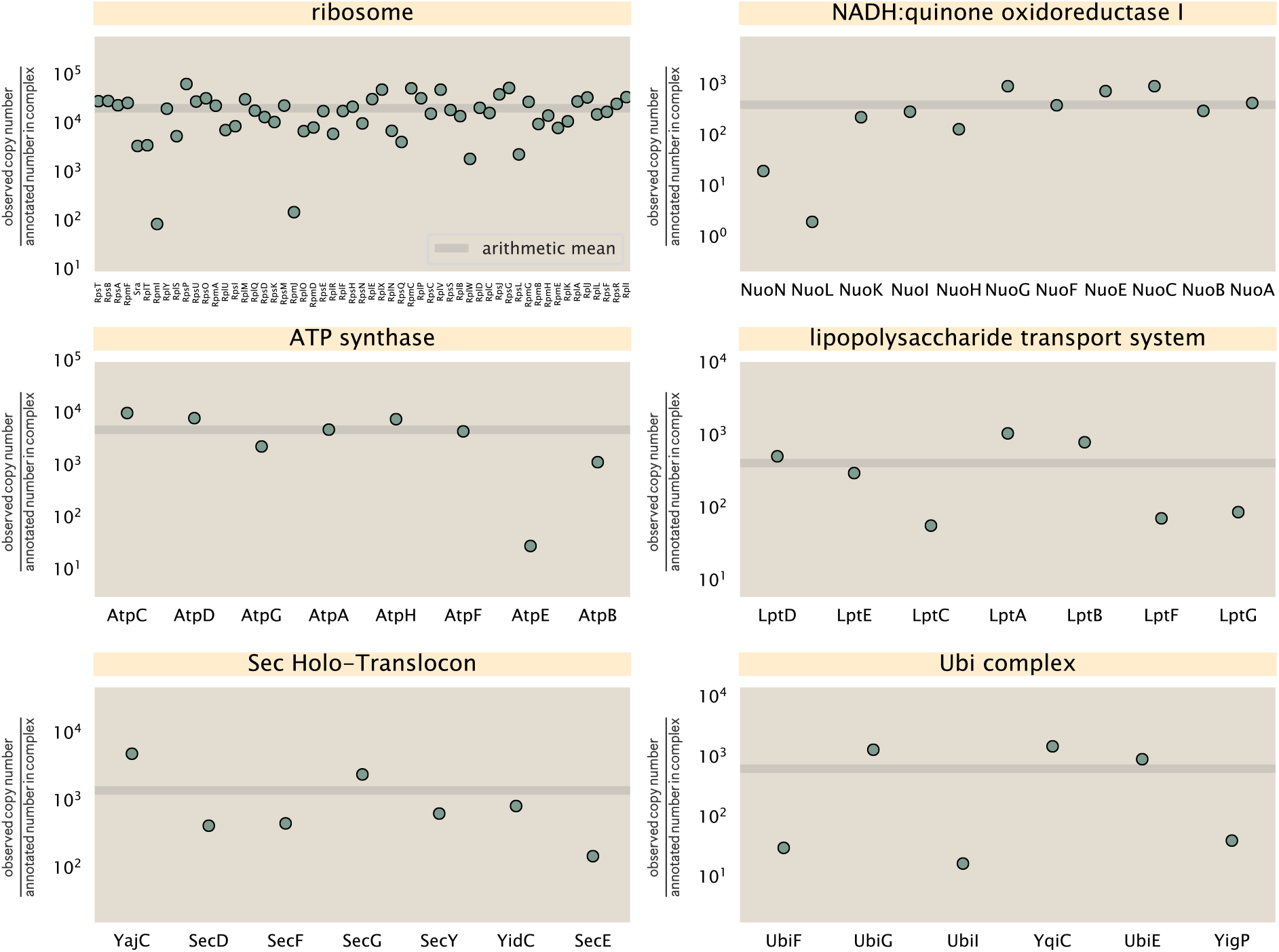
Calculation of the mean complex abundance from measurements of single subunits. Six of the largest complexes (by number of subunits) in *E. coli*. Points correspond to the maximum number of complexes that can be formed given measurement of that individual protein. Solid grey line corresponds to the arithmetic mean across all subunits. These data correspond to measurements from ***Schmidt et al.*** (***2016***) ina glucose-supplemented minimal growth medium.

## Extending Estimates to a Continuum of Growth Rates

In the main text, we considered a standard stopwatch of 5000 s to estimate the abundance of the various protein complexes considered. In addition to point estimates, we also showed the estimate as a function of growth rate as transparent grey curves. In this section, we elaborate on this continuum estimate, giving examples of estimates that scale with either cell volume, cell surface area, or number of origins of replication.

## Estimation of the total cell mass

For many of the processes estimated in the main text we relied on a cellular dry mass of ≈ 300 fg from which we computed elemental and protein fractions using knowledge of fractional composition of the dry mass. At modest growth rates, such as the 5000 s doubling time used in the main text, this is a reasonable number to use as the typical cell mass is ≈ 1 pg and *E. coli* cells can approximated as 70% water by volume. However, as we have shown in the preceding sections, the cell size is highly dependent on the growth rate. This means that a dry mass of 300 fg cannot be used reliably across all growth rates.

Rather, using the phenomenological description of cell volume scaling exponentially with growth rate, and using a rule-of-thumb of a cell buoyant density of ≈ 1.1 pg / fL (BNID: 103875), we can calculate the cell dry mass across a range of physiological growth rates as

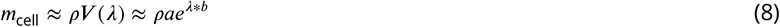

where *a* and *b* are constants with units of μm^3^ and hr, respectively. The value of these constants can be estimated from the careful volume measurements performed by ***Si et al.*** (***2017***), as considered in Appendix Estimation of Cell Size and Surface Area earlier.

## Complex Abundance Scaling With Cell Volume

Several of the estimates performed in the main text are implicitly dependent on the cell volume. This includes processes such as ATP utilization and, most prominently, the transport of nutrients, whose demand will be proportional to the volume of the cell. Of the latter, we estimated the number of transporters that would be needed to shuttle enough carbon, phosphorus, and sulfur across the membrane to build new cell mass. To do so, we used elemental composition measurements combined with a 300 fg cell dry mass to make the point estimate. As we now have a means to estimate the total cell mass as a function of volume, we can generalize these estimates across growth rates.

Rather than discussing the particular details of each transport system, we will derive this scaling expression in very general terms. Consider that we wish to estimate the number of transporters for some substance *X*, which has been measured to be made up some fraction of the dry mass, *θ_X_*. If we assume that, irrespective of growth rate, the cell dry mass is relatively constant (***Basan et al., 2015***) and ≈ 30% of the total cell mass, we can state that the total mass of substance *X* as a function of growth rate is

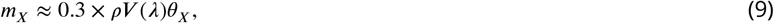

where we have used *ρV (λ)* as an estimate of the total cell mass, defined in ***Equation 8***. To convert this to the number of units *N_X_* of substance X in the cell, we can use the formula weight *w_X_* of a single unit of *X* in conjunction with ***Equation 9***,

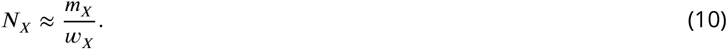

To estimate the number of transporters needed, we make the approximation that loss of units of *X* via diffusion through porins or due to the permeability of the membrane is negligible and that a single transporter complex can transport substance *X* at a rate *r_X_*. As this rate r_X_ is in units of *X* per time per transporter, we must provide a time window over which the transport process can occur. This is related to the cell doubling time τ, which can be calculated from the the growth rate *λ* as τ = log(2)/*λ*. Putting everything together, we arrive at a generalized transport scaling relation of

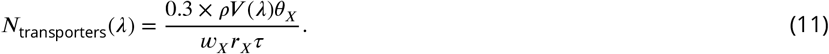

This function is used to draw the continuum estimates for the number of transporters seen in Figures 2 and 3 as transparent grey curves. Occasionally, this continuum scaling relationship will not precisely agree with the point estimate outlined in the main text. This is due to the choice of ≈ 300 fg total dry mass per cell for the point estimate, whereas we considered more precise values of cell mass in the continuum estimate. We note, however, that both this scaling relation and the point estimates are meant to describe the order-of-magnitude observed, and not the predict the exact values of the abundances.

***Equation 11*** is a very general relation for processes where the cell volume is the “natural variable” of the problem. This means that, as the cell increases in volume, the requirements for substance X also scale with volume rather than scaling with surface area, for example. So long as the rate of the process, the fraction of the dry mass attributable to the substance, and the formula mass of the substance is known, ***Equation 11*** can be used to compute the number of complexes needed. For example, to compute the number of ATP synthases per cell, ***Equation 11*** can be slightly modified to the form

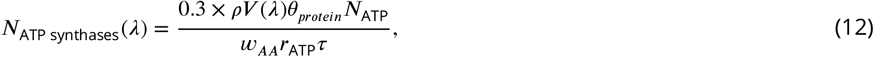

where we have included the term *N_ATP_* to account for the number of ATP equivalents needed per amino acid for translation (≈ 4, BNID: 114971), and *w_AA_* is the average mass of an amino acid. The grey curves in Figure 4 of the main text were made using this type of expression.

## A Relation for Complex Abundance Scaling With Surface Area

In our estimation for the number of complexes needed for lipid synthesis and peptidoglycan maturation, we used a particular estimate for the cell surface area (≈ 5 μ*m*, BNID: 101792) and the fraction of dry mass attributable to peptidoglycan (≈ 3%, BNID: 101936). Both of these values come from glucose-fed *E. coli* in balanced growth. As we are interested in describing the scaling as a function of the growth rate, we must also consider how these values scale with cell surface area, which is the natural variable for these types of processes. In the coming paragraphs, we highlight how we incorporate a condition-dependent surface area into our calculation of the number of lipids and murein monomers that need to be synthesized and crosslinked, respectively.

## Number of Lipids

To compute the number of lipids as a function of growth rate, we make the assumption that some features, such as the surface area of a single lipid (*A*_lipid_ ≈ 0.5 nm^2^, BNID: 106993) and the total fraction of the membrane composed of lipids (≈ 40%, BNID: 100078) are independent of the growth rate. Using these approximations combined with ***Equation 2***, and recognizing that each membrane is composed of two leaflets, we can compute the number of lipids as a function of growth rate as

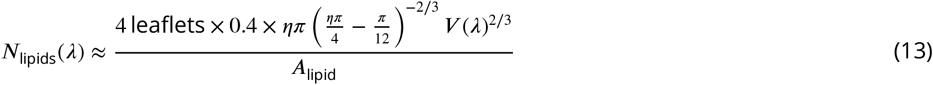

where *η* is the length-to-width aspect ratio and *V* is the cell volume.

## Number of Murein Monomers

In calculation of the number of transpeptidases needed for maturation of the peptidoglycan, we used an empirical measurement that ≈ 3% of the dry mass is attributable to peptidoglycan and that a single murien monomer is *m_murein_* ≈ 1000 Da. While the latter is independent of growth rate, the former is not. As the peptidoglycan exists as a thin shell with a width of *w* ≈ 10 nm encapsulating the cell, one would expect the number of murein monomers scales with the surface area of this shell. In a similar spirit to our calculation of the number of lipids, the total number of murein monomers as a function of growth rate can be calculated as

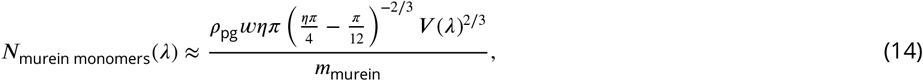

where *ρ*_pg_ is the density of peptidoglycan.

## Complex Abundance Scaling With Number of Origins, and rRNA Synthesis

While the majority of our estimates hinge on the total cell volume or surface area, processes related to the central dogma, namely DNA replication and synthesis of rRNA, depend on the number of chromosomes present in the cell. As discussed in the main text, the ability of *E. coli* to parallelize the replication of its chromosome by having multiple active origins of replication is critical to synthesize enough rRNA, especially at fast growth rates. Derived in ***Si et al.*** (***2017***) and reproduced in the main text and Appendix Estimation of 〈#ori〉 / 〈#ter〉 and 〈#ori〉 as below, the average number of origins of replication at a given growth rate can be calculated

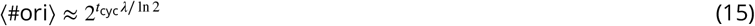

where *t_cyc_* is the total time of replication and division. We can make the approximation that *t_cyc_* ≈ 70 min, which is the time from the initiation of chromosomal replication until division. This time corresponds to the sum of the so-called C and D periods of the cell cycle, which correspond to the time it takes to replicate the entire chromosome (C period) and the time from completion to eventual division (D period) ***Helmstetter and Cooper*** (***1968***).

In the case of rRNA synthesis, the majority of the rRNA operons are surrounding the origin of replication. Thus, at a given growth rate *λ*, the average dosage of rRNA operons per cell *D_rRNA_* is

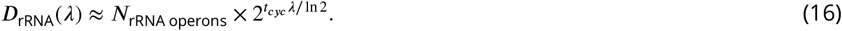

This makes the approximation that *all* rRNA operons are localized around the origin. In reality, the operons are some distance away from the origin, making ***Equation 16*** an approximation (***Dennis et al., 2004***).

In the main text, we stated that at a growth rate of 0.5 hr^−1^, there is ≈ 1 chromosome per cell. While a fair approximation, ***Equation 15*** illustrates that is not precisely true, even at slow growth rates. In estimating the number of RNA polymerases as a function of growth rate, we consider that regardless of the number of rRNA operons, they are all sufficiently loaded with RNA polymerase such that each operon produces one rRNA per second. Thus, the total number of RNA polymerase as a function of the growth rate can be calculated as

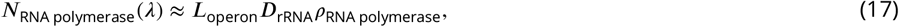

where *L_operon_* is the total length of an rRNA operon (≈ 4500 bp) and *ρ*_RNA polymerase_ is packing density of RNA polymerase on a given operon, taken to be 1 RNA polymerase per 80 nucleotides.

## Calculation of active ribosomal fraction

In the main text we used the active ribosomal fraction *f*_*a*_ that was reported in the work of ***Dai et al.*** (***2016***) to estimate the active ribosomal mass fraction Φ_*R*_ × *f*_*a*_ across growth conditions. We lacked any specific model to consider how *f*_*a*_ should vary with growth rate, and instead find that the data is well-approximated by fitting to an exponential curve (*f*_*a*_ = −0.889 *e*^4.6.*λ*^ + 0.922; dashed line in inset of ***Figure 10***(C)). We use this function to estimate *f*_*a*_ for each of the data points shown in ***Figure 10***(C).

## Estimation of 〈#ori〉/〈#ter〉 and 〈#ori〉

*E. coli* shows robust scaling of cell size with the average number of origins per cell, 〈#ori〉 (***Si et al., 2017***). Since protein makes up a majority of the cell’s dry mass, the change in cell size is also a reflection of the changes in proteomic composition and total abundance across growth conditions. Given the potential constraints on rRNA synthesis and changes in ribosomal copy number with 〈ori〉, it becomes important to also consider how protein copy numbers vary with the state of chromosomal replication. This is particularly true when trying to make sense of the changes in ribosomal fraction and growth-rate dependent changes in proteomic composition at a mechanistic level. As considered in the main text, it is becoming increasingly apparent that regulation through the secondary messengers (p)ppGpp may act to limit DNA replication and also reduce ribosomal activity in poorer nutrient conditions. In this context, both 〈ori〉, as well as the 〈#ori〉/〈#ter〉 ratio become important parameters to consider and keep track of. An increase in 〈#ori〉 / 〈#ter〉 ratio in particular, causes a relatively higher gene dosage in rRNA and r-protein genes due to skew in genes near the origin, where the majority of these are located

In the main text we estimated the change in 〈#ori〉 with growth rate using the nutrient-limited wild-type cell data from ***Si et al.***(***2017***). We consider their measurements of DNA replication time (*t_C′_*, ‘C’ period of cell division), total cell cycle time (*t_cyc′_*, ‘C’ + ‘D’ period of cell division), and doubling time τ from wild-type *E. coli* growing across a range of growth conditions. Here we show how we esimate this parameter, as well as the 〈#ori〉/ 〈# ter〉 ratio from their data. We begin by considering 〈#ori〉. If the cell cycle time takes longer than the time of cell division, the cell will need to initiate DNA replication more often than its rate of division, 2^*λt*^ = 2^*ln*(2)·t/τ^ to maintain steady state growth. Cells will need to do this in proportion to the ratio *λ_cyc_*/*λ* = *t_cyc_*/τ, and the number of origins per cell (on average) is then given by 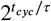. The average number of termini will in contrast depend on the lag time between DNA replication and cell division, *t_D′_*, with 〈#ori〉 / 〈#ter〉 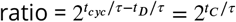.

In Figure A10(A) and (B) we plot the measured *t_C_* and *t_cyc_* values versus the doubling time from ***Si et al.*** (***2017***). The authors estimated *t*_*C*_ by marker frequency analysis using qPCR, while *t_cyc_* = *t_C_* + *t_D_* were inferred from *t_C_* and τ. In the plots we see that both *t_C_* and *t_cyc_* reach a minimum at around 40 and 75 minutes, respectively. For a C period of 40 minutes, this would correspond to a maximum rate of elongation of about 1,000 bp/sec. Since we lacked a specific model to describe how each of these parameters vary with growth condition, we assumed that they were linearly dependent on the doubling time. For each parameter, *t_C_* and *t_cyc_*, we split them up into two domains corresponding to poorer nutrient conditions and rich nutrient conditions (cut off at τ ≈ 40 minutes where chromosomal replication becomes nearly constant). The fit lines are shown as solid black lines. In Figure A10(C) and (D) we also show *t_C_* and *t_cyc_* as a function of growth rate *λ* along with our piecewise linear fits, which match the plots in the main text.

## Derivation of Minimal Model for Nutrient-Mediated Growth Rate Control

Here we provide a derivation of the minimal model for growth rate control under nutrient-limited growth. By growth rate control, we are specifically referring to the ability of bacteria to modulate their proteome (*N*_pep_, *R*, Φ_*R*_) and cell size as nutrient conditions change, with slower growing cells generally being smaller in size (***Ojkic et al., 2019***). This capability provides bacteria with a particular benefit when nutrients are more scarce since it will mean there is a smaller net demand on carbon, phosphorus, sulfur, and nitrogen. The specific goal of developing this model is to help us better explore the overall constraints on growth that follow from 1) our observation that many of the cellular processes we’ve considered require increased protein abundance at faster growth rates, and 2) a strict limit on growth rate that is governed by the ribosomal synthesis rate and ribosomal mass fraction Φ_*R*_.

In ***Figure 12***(A) of the main text we provide a schematic of the model, where we consider growth as simply governed by the rate of protein synthesis (*r*_*t*_ × *R* × *f*_*a*_). In order to grow rapidly, at least to the extent possible, these three parameters need to be maximized (with *r*_*t*_ ≤ 17 amino acids per second, and *f*_*a*_ ≤ 1 reported in the work of ***Dai et al.*** (***2016***)). The elongation rate *r*_*t*_ will depend on how quickly ribosomes can match codons with their correct amino-acyl tRNA, along with the subsequent steps of peptide bond formation and translocation. This ultimately depends on the cellular concentration amino acids, which we treat as a single effective species, [*AA*]_eff_.

**Figure A10.**
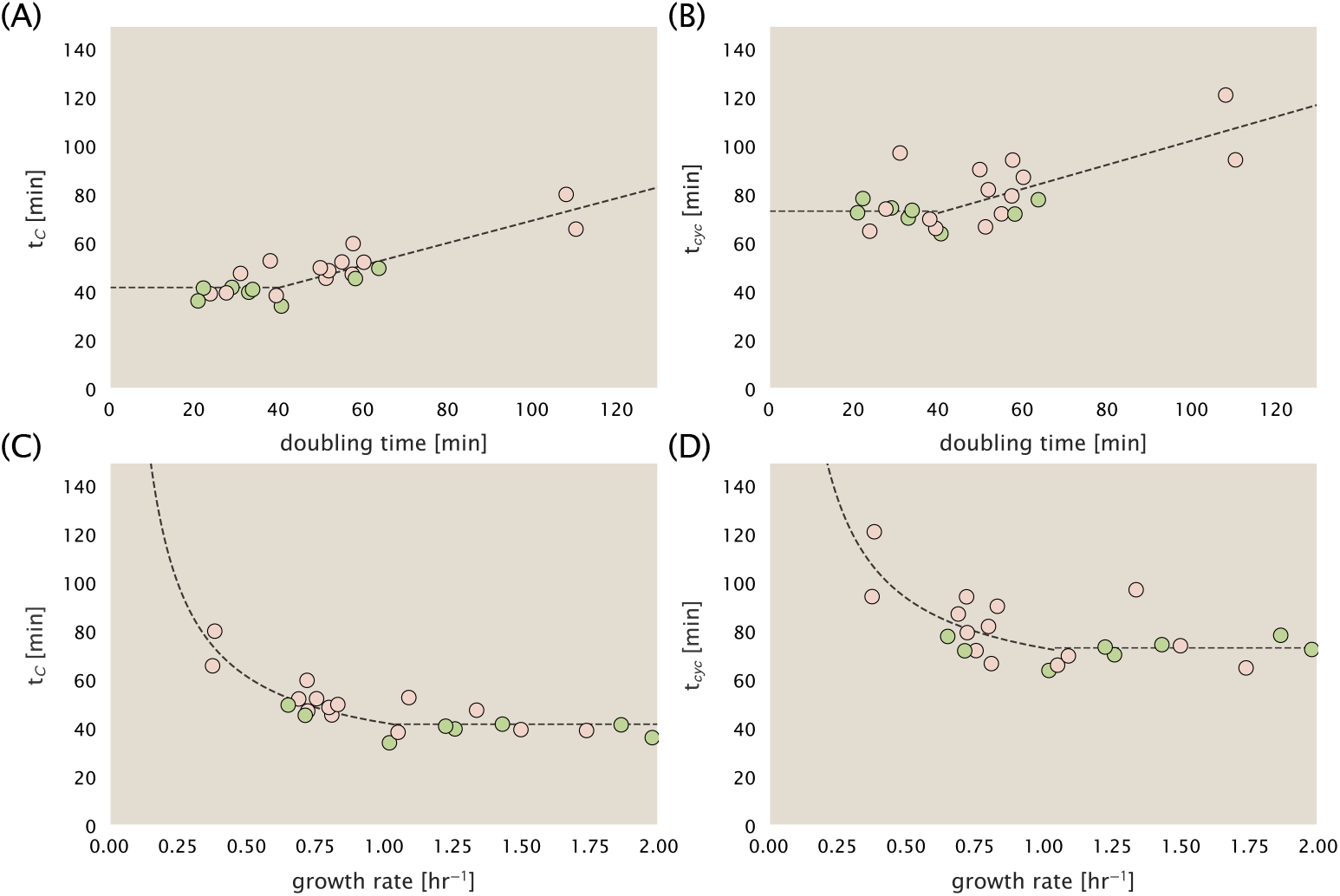
Estimation of 〈#ori〉 / 〈#ter〉 and 〈#ori〉 using data from Si *et al.* (2017). (A) and (B) plot the reported *t_C_* and *t_cyc_* as a function of cell doubling time τ, respectively. The dashed lines show a piecewise fit to the data. For short doubling times (rich media), *t_C_* and *t_cyc_* constant (*t_C_* = 42 minutes, *t_cyc_* = 73 minutes). At the transition, taken to occur at 40 minutes, the dashed line corresponds to an assumed proportional increase in each parameter as a function of the doubling time (*t_C_* = 0.46 τ + 23.3 minutes, *t_cyc_* = 0.50 τ + 52.7 minutes). (C) and (D) plot the same data as in (A) and (B), but as a function of growth rate, given by *λ* = *ln*(2)/τ.

In our model, we need to determine the rate of peptide elongation *r*_*t*_, which we consider as simply depending on the supply of amino acids (and, therefore, also amino-acyl tRNAs) through a parameter *r_AA_* in units of AA per second, and the rate of amino acid consumption by protein synthesis (*r*_*t*_ × *R* × *f*_*a*_). The balance between these two rates will determine the effective amino acid concentration in the cell [*AA*]_eff_. An important premise for this formulation is growing evidence that cells are able to modulate their biosynthesis activity according to nutrient availability (i.e. extent of chromosomal replication, transcriptional, and translation activity) through secondary-messenger molecules like (p)ppGpp (***Hauryliuk et al., 2015***; ***Zhu and Dai, 2019***; ***Kraemer et al., 2019***; ***Fernández-Coll et al., 2020***; ***Büke et al., 2020***). Given our observation that protein synthesis and energy production are not limiting, we assume that other molecular players required by ribosomes like elongation factors and GTP are available in sufficient abundance. In addition, experimentally, the relative number of tRNA and elongation factor EF-Tu per ribosome have been found to increase in poorer nutrient conditions ***Pedersen*** (***1978***); ***Dong et al.*** (***1996***); ***Klumpp et al.*** (***2013***)).

We begin by considering a coarse-grained description of peptide elongation, which includes 1) the time required to find and bind each correct amino-acyl tRNA, and 2) the remaining steps in peptide elongation that will not depend on the amino acid availability. These time scales will be related to the inverse of the elongation rate *r*_*t*_′,

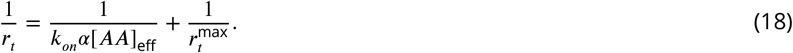

where we have assumed that the rate of binding by amino-acyl tRNA *k_on_* is proportional to [*AA*]_eff_ by a constant α. 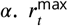 refers to the maximum elongation rate. This leads to a Michaelis-Menten dependence of the elongation rate *r*_*t*_ on the effective amino acid concentration [*AA*]_eff_ (***Klumpp et al., 2013***; ***Dai et al., 2016***). We can re-write this more succinctly in terms of an effective dissociation constant,

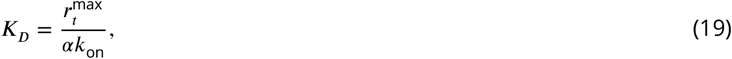

where the elongation rate *r*_*t*_ is now given by

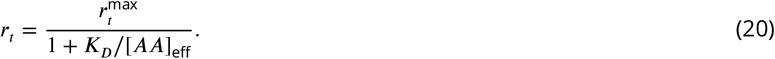

The rate of amino acid supply *r_AA_* will vary with changing nutrient conditions and the cell can maintain [*AA*]_eff_ by tuning the rate of amino acid consumption, *r*_*t*_ × *R* × *f*_*a*_. Thus, [*AA*]_eff_ is determined by the difference in the rate of amino acid synthesis (or import, for rich media) and/or tRNA charging, *r_AA_*, and the rate of consumption, *r*_*t*_ × *R* × *f*_*a*_. Over an arbitrary length of time t of cellular growth, the cell will grow in volume, requiring us to consider these rates in terms of concentration rather than absolute numbers, with [*AA*]_eff_ given by,

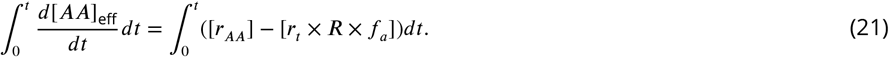

This considers the net change in amino acid concentration over a time from 0 to *t*, with the square brackets indicating concentrations per unit time. Integrating ***Equation 21*** yields.

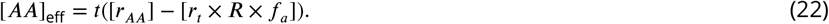

Alternatively, to connect to the experimental data in terms of absolute ribosome copy number *R* we can consider a unit volume *V*,

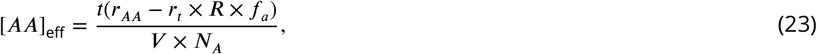

where *r_AA_* is in units of AA per unit time and *r*_*t*_ is in units of AA per unit time per ribosome. *N*_A_ refers to Avogadro’s number and is needed to convert between concentration and absolute numbers per cell. With an expression for [*AA*]_eff_ in hand, we can now solve ***Equation 20*** for *r*_*t*_ which is a quadratic function with a physically-meaningful root of

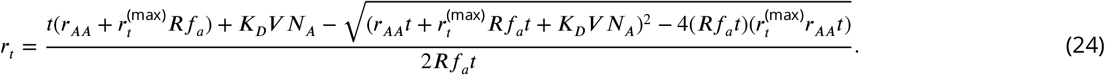

This is the key equation that allows us to calculate growth rate for any combination of *N*_pep_, *R*, *f_a′_*, and cell size *V* as a function of amino acid supply *r_AA_* (***Equation 3*** of the main text). We refer the reader to Section “A Minimal Model of Nutrient-Mediated Growth Rate Control “ of the main text for our exploration of this model in the context of the proteomic data.

We end this section by noting several distinctions of this formulation with previous work. The first, as noted in the main text, relates to the now seminal work of ***Scott et al.*** (***2010***), which provides a treatment of resource allocation that partitions of the proteome into sectors – including one for ribosome-associated proteins and one for metabolic proteins. As cells grow faster, there is a notable change in the mass fraction of these sectors, with an increase in ribosomal content that is predominantly achieved at the expense of a decrease in the metabolic sector. By including an additional constraint through the phenomenological parameter *v*, which characterizes the quality of the growth medium ***Scott et al.*** (***2010***); ***Klumpp et al.*** (***2013***); ***Klumpp and Hwa*** (***2014***), the authors derive a model of growth rate, dependent on optimal resource allocation. Here we have developed a model that considers the effect of changes in absolute protein abundance and ribosomal content, and consider how these influence the achievable growth rate. In addition, by accounting for the metabolic supply of amino acids directly though their availability in the cell (i.e. [*AA*]_eff_), we are able to consider how the balance between translation-specific metabolic capacity and translational capacity influences both the elongation rate *r*_*t*_ and growth rate *λ*.

The second and last point we note is that the recent works from ***Dai et al.*** (***2016***) and ***Klumpp et al.*** (***2013***) also employ a similar coarse-graining of translation elongation as we’ve considered above. Here, however, a notable distinction is that the authors consider the entire ternary complex (i.e. the complex of amino-acyl tRNA, EF-Tu, and GTP) as rate limiting. Further, through an assumed proportionality between ternary complex and ribosome abundance, they arrive at a formulation of elongation rate *r*_*t*_ that exhibits a Michaelis-Menten dependence on the ribosomal fraction Φ_*R*_. They demonstrate that all their measurements of elongation rate, even upon addition of sublethal doses of chloramphenicol (which cause an increase in both *r*_*t*_ and Φ_*R*_), can be collapsed onto a single curve described by this Michaelis-Menten dependence. There is always a benefit to increase their ribosomal fraction Φ_*R*_ on growth rate when nutrient conditions allow (see Section “Maximum Growth Rate is Determined by the Ribosomal Mass Fraction” on the main text), and this trend in the data in part follows from the tendency for cells to increase Φ_*R*_ and better maximize *r*_*t*_ as nutrient conditions improve. In addition, it does not account for the decrease in the fraction of actively translating ribosome *f*_*a*_ that was strikingly apparent at slow growth rates or in sublethal doses of chloramphenicol in the work of ***Dai et al.*** (***2016***). Through ***Equation 24*** we also account for changes in the fraction of actively translating ribosomes. Ultimately, we find that cells are able to maximize both Φ_*R*_, *r*_*t*_, and their growth rate only to the extent allowed by the nutrient conditions (i.e. via *r*_*AA*_) and through the maintenance of the cellular pool of amino acids [*AA*]_eff_, amino-acyl tRNA, GTP, as well as the synthesis of other key molecular constituents like EF-Tu.

